# A comparative analysis of planarian genomes reveals regulatory conservation in the face of rapid structural divergence

**DOI:** 10.1101/2023.12.22.572568

**Authors:** Mario Ivankovic, Jeremias N. Brand, Luca Pandolfini, Tom Brown, Martin Pippel, Andrei Rozanski, Til Schubert, Markus A. Grohme, Sylke Winkler, Laura Robledillo, Meng Zhang, Azzurra Codino, Stefano Gustincich, Miquel Vila-Farré, Shu Zhang, Argyris Papantonis, André Marques, Jochen C. Rink

**Author notes:** Authors contributed equally (Mario Ivankovic, Jeremias N. Brand).

## Abstract

The planarian *Schmidtea mediterranea* can regenerate its entire body from small tissue fragments and is studied as regeneration model species. The assembly and functional analysis of planarian genomes has proven challenging due its high A/T content (70% A/T), repetitive nature, and limited transferability of routine laboratory protocols due to their divergent biochemistry. Only few and often fragmented genome assemblies are currently available, and open challenges include the provision of well-annotated chromosome-scale reference assemblies of the model species and other planarians for a comparative genome evolution perspective. Here we report a haplotype-phased, chromosome-scale genome assembly and high-quality gene annotations of the sexual S2 strain of *S. mediterranea* and provide putative regulatory region annotations via optimized ATAC-seq and ChIP-seq protocols. To additionally leverage sequence conservation for regulatory element annotations, we generated chromosome-scale genome assemblies and chromatin accessibility data for the three closest relatives of *S. mediterranea*: *S. polychroa*, *S. nova*, and *S. lugubris*. We find substantial divergence in protein-coding sequences and regulatory regions, yet reveal remarkable conservation in ChIP-mark bearing open chromatin regions identified as promoters and enhancers in *S. mediterranea*. The resulting high-confidence set of evolutionary conserved enhancers and promoters provides a valuable resource for the analysis of gene regulatory circuits and their evolution within the taxon. In addition, our four chromosome-scale genome assemblies provide a first comparative perspective on planarian genome evolution. Our analyses reveal frequent retrotransposon-associated chromosomal inversions and inter-chromosomal translocations that lead to a degradation of synteny across the genus. Interestingly, we further find independent and near-complete losses of the ancestral metazoan synteny across *Schmidtea* and two other flatworm groups, indicating that platyhelminth genomes largely evolve without syntenic constraints. Our work provides valuable genome resources for the planarian research community and sets a foundation for the comparative genomics of planarians. We reveal a contrast between the fast structural evolution of planarian genomes and the conservation of their regulatory elements, suggesting a unique genome evolution in flatworms where gene positioning may not be essential.

## Introduction

Evolution acts on genomic changes to bring about the diversity of life. For example, single nucleotide changes in coding gene sequences duplicate the goldfish tail fin ^1^ or cause nose loss in humans ^2^; changes in gene regulatory regions are associated with profound evolutionary body plan changes ^3,4^, e.g. limb loss in snakes ^5^, and gene loss is emerging as important mechanism in trait evolution ^6,7^. On the other hand, the rapidly increasing number of sequenced genomes indicates that genome structure may also be evolutionarily constrained.

Synteny (the association of genes on a chromosome or linkage group) is deeply conserved across animals, with Metazoan Ancestral Linkage Groups (MALG) being conserved between Sponges, Cnidaria, and Bilateria and thus in numerous animal phyla ^8–10^. However, some groups like Nematodes and Drosophilids have lost this ancestral synteny, but intriguingly, as ancestral synteny disappears, new linkage groups emerged in these species ^11,12^. The finding that the arrangement of genes in the genome at the mega-base scale is important for gene regulation ^13^ provides a rationale for the evolutionary conservation of synteny. Indeed, studies mostly in vertebrates have shown that gene regulation is influenced by hierarchical levels of chromatin organization ^14^ and that the modulation of chromatin organization can give rise to genetic diseases ^15,16^, cancer ^17,18^ or even the origin of evolutionary novelties ^19–21^. However, not all taxa exhibit consistent genomic organization features or their significance remains ambiguous ^22^. Therefore, the current observations indicate that certain taxonomic groups may display specific patterns of genome evolution and that currently under-sampled clades may reveal additional patterns.

Planarians are one example of a large and so far, scarcely sequenced group of animals. As an order (Tricladida) within the diverse and species-rich phylum Platyhelminthes (flatworms), planarians are studied for their astonishing regenerative abilities and their abundant adult pluripotent stem cells (neoblasts) ^23,24^. As the only division-competent cell outside the reproductive system, neoblast proliferation and differentiation drive homeostatic tissue maintenance and give rise to the germ cell lineages of the hermaphroditic reproductive system of planarians ^25^. However, planarian reproductive strategies are highly diverse at the taxon level and range from sexual reproduction to various forms of parthenogenesis and asexual propagation by fission and regeneration ^26^. The formation of germ cells from somatic neoblasts and the “inheritance” of many parental neoblasts by a fission fragment raises profound questions regarding genome evolution, including the maintenance of “genetic self” within the pluripotent stem cell population of an individual ^27^. Given these profound questions regarding genome evolution and the rapid growth of the planarian research community in general, high-quality genomic resources are highly desirable.

Although the planarian model species *Schmidtea mediterranea* was amongst the early cohort of Sanger-sequenced genomes, the resulting assembly was highly fragmented ^28^. Only the advent of long-read sequencing achieved significant assembly contiguity ^29^, which recent Hi-C scaffolding extended to a chromosome-scale ^30^. The strong compositional bias (> 70% A/T), abundant repeats inclusive of giant > 30 kb Burro retroelements and inbreeding-resistant heterozygosity ^29^ associated with a large chromosomal inversion on Chromosome1 ^30^ are some of the reasons of why the *S. mediterranea* genome remains an assembly challenge. The extent by which these peculiarities are species-specific or general features of planarian genomes remains unknown, as the only other currently available planarian genome assembly is a highly fragmented assembly of *Dugesia japonica* ^31^. Additional planarian genome assemblies are therefore essential for gaining a comparative perspective and also as an entry point for probing the genetic basis of the rich phenotypic biodiversity within the group ^32^. Parallel efforts have initiated the functional annotation and analysis of *S. mediterranea* genomic features utilizing ATAC-seq and ChIP-seq ^33–41^. These efforts have provided first insights into the epigenetic state of planarian genes, the function of epigenetic regulators and gene regulatory elements in individual cell types. However, the systematic annotation of gene regulatory elements across the genome remains a challenge. In addition, the ferocious nuclease activity in planarian cell extracts and abundant contaminants in nucleic acid preparations still leave room for further optimizations of the functional genomics protocols within the community.

Here, we present four high-quality genome assemblies of the planarian model species *S. mediterranea* and its three closest known relatives, *S. polychroa*, *S. nova,* and *S. lugubris.* For the *S. mediterranea* research community, the haplotype-phased genome assembly and substantially improved gene annotations constitute important model system resources. Using improved ATAC-seq and ChIP-seq protocols, we further identify and annotate regulatory elements that are conserved across the genus and present attractive targets for functional investigations. In contrast, we find that the genome structure is poorly conserved within the genus and that *Schmidtea*, parasitic flatworms and the free-living early-branching flatworm *Macrostomum hystrix* have independently lost the Metazoan Ancestral Linkage Groups. Altogether, our study provides a first comparative perspective on the *S*. *mediterranea* genome and indicates that synteny may not constrain the structural evolution of planarian genomes and those of other flatworms.

## Results

### *Schmidtea mediterranea* genome

The current *S. mediterranea* reference genome (dd_Smes_g4 ^29^) is a haploid consensus assembly containing 481 contigs. Recently, a scaffolding of this genome (referred to here as schMedS2) has revealed substantial differences between the haplotypes, especially on Chromosome 1 ^30^. This makes a haplotype-phased assembly desirable. Towards this goal and to close the remaining sequencing gaps, we re-sequenced the *S. mediterranea* genome with Pacific Biosciences’ high-fidelity Circular Consensus Sequencing (CCS, also known as HiFi) reads and used Hi-C for scaffolding. The new assembly, designated S3, consists of two pseudo-haplotypes: S3h1 and S3h2, and a merged version of the two, referred to as S3BH for “S3 both haplotypes”. The total assembly sizes of both S3h1 and S3h2 are slightly larger than the previous dd_Smes_g4 assembly, with 840Mb and 820Mb compared to 774Mb (Table 1, Additional File 1: Section 1.1). The N50 values of 270Mb and 269Mb indicate high contiguity with 95% and 96% of the total assembly contained in the four largest scaffolds that closely match the known karyology in size and number (1n = 4, Table 1). The Hi-C contact maps indicate high contiguity in both phases, thus justifying the designation as a “chromosome-scale” assembly (Fig. 1A). Nevertheless, not all scaffolds are capped by telomere repeats and unincorporated contigs remain (S3h1 and S3h2 contain 662 and 432 contigs, respectively). The largest fraction of unincorporated contigs is comprised of various repeat sequences (satellite DNA, telomere repeats, rRNA clusters) that could not be placed during the assembly process. However, others contain annotated genes (S3h1: 505 genes on 216 contigs, S3h2: 509 genes on 197 contigs), pointing towards remaining localized assembly ambiguities.

**Figure 1.**
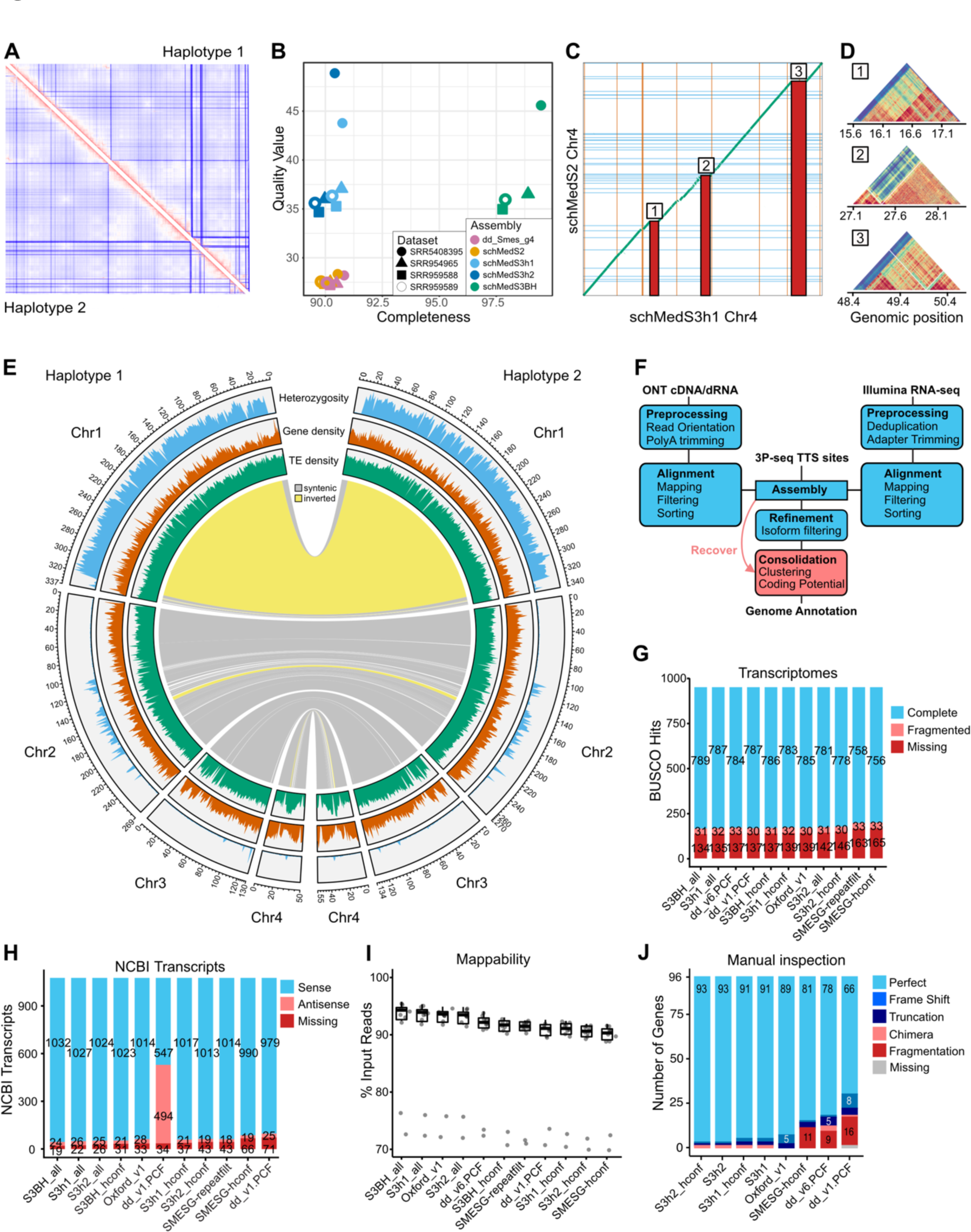
Quality control metrics and description of the *S. mediterranea* genome and annotation. **A:** HiC contact map of the reads used for scaffolding on S3h1 (upper right) and S3h2 (lower left), showing high contact intensity in red and low contact intensity in blue. **B:** Results of a Merqury analysis using the four indicated Illumina shot-gun datasets, none of which was used in the analysis. **C:** Dotplot representing a whole genome alignment between Chromosome 4, inferred with minimap2, of the previously scaffolded assembly (schMedS2) on the y-axis and the genome in this study (schMedS3h1). Blue lines indicate scaffold gaps in schMedS2 and red lines indicate scaffold gaps in schMedS3h1. Numbered red bars indicate alignment gaps >1Mb, which contain highly repetitive satellite DNA absent in the previous assembly. **D:** self-similarity heatmap, calculated with stainedglass, of the gaps indicated in C showing their high self-similarity, typical of centromeric or pericentromeric repeats. **E:** Comparison between the two pseudohaplotypes of schMedS3. The chord diagram in the center indicates synteny regions and inversions (in yellow) between the haplotypes. Density plots in the outer three circles show the distribution of transposable elements (TE), genes, and heterozygosity. **F:** Schematic representation of the hybrid gene annotation workflow. **G:** BUSCO assessment of all benchmarked annotations. **H:** Transcriptome completeness assessment using the 1054 *S. mediterranea* transcripts deposited in GenBank. **I:** Mappability of publicly available RNA-Seq datasets against benchmarked annotations. **J:** Results of a manual inspection of 96 gene models. The scores only reflect the best predicted transcript /locus. **G-J:** Benchmarked gene annotations: S3h1, S3h2, S3BH: this study; dd_v1 the non-stranded dd_Smes_v1 assembly of the sexual strain of *S. mediterranea* ^42^; dd_v6 the dd_Smed_v6 assembly of the asexual strain of *S. mediterranea* ^42^; SMESG the gene prediction on basis of the previous dd_smes_g4 *S. mediterranea* genome assembly ^42^; Oxford_v1 a composite annotation of Neiro et al. ^39^, García-Castro et al. ^45^, and SMESG.

**Table 1.** Summary statistics for the genome assemblies and annotation of the four *Schmidtea* species. Given is the number of contigs in the assembly, assembly length, GC content, N50 values, the percentage of assembly that is on chromosome scaffolds, and the number of bases that are not placed on a chromosome. For the annotation the number of genes and transcripts and their length and span are indicated.

|  | **schMedS3h1** | **schMedS3h2** | **schPol2** | **schNov1** | **schLug1** |
| --- | --- | --- | --- | --- | --- |
| **Genome assembly statistics** |  |  |  |  |  |
| # contigs | 662 | 432 | 1013 | 283 | 320 |
| Total length (bp) | 840,173,815 | 819,865,861 | 781,290,622 | 1,251,382,582 | 1,499,048,548 |
| GC (%) | 29.59 | 29.59 | 28.11 | 28.13 | 27.87 |
| N50 (bp) | 270,168,396 | 268,961,546 | 189,691,935 | 455,729,997 | 498,167,912 |
| chromosome scaff. (%) | 95 | 96 | 89 | 98 | 99 |
| unplaced (Mb) | 42 | 32.8 | 85.9 | 25 | 15 |
| **Gene annotation statistics** |  |  |  |  |  |
| # Genes | 58,735 | 58,551 | 41,915 | 41,539 | 48,029 |
| Median gene length | 1113 | 1104 | 1137 | 1005 | 1039 |
| Shortest gene | 83 | 83 | 68 | 85 | 81 |
| Longest gene | 379,876 | 379,876 | 399,802 | 422,721 | 513,405 |
| Total gene length | 395,812,229 | 391,333,147 | 360,601,815 | 508,048,891 | 610,689,353 |
| # Transcripts | 86,102 | 85,741 | 60,887 | 57,075 | 66,510 |
| Median transcript length | 936 | 930 | 950 | 842 | 860 |
| Shortest transcript | 83 | 83 | 68 | 85 | 61 |
| Longest transcript | 76,733 | 94,201 | 32,100 | 36,224 | 28,220 |
| Total transcript length | 111,559,651 | 110,732,822 | 79,151,531 | 68,678,530 | 79,879,797 |

To quantitatively evaluate the quality of the S3 assemblies compared to previous assemblies, we first assessed base-pair accuracy and assembly completeness via a Merqury analysis of four independent short-read gDNA datasets (see Methods). Fig. 1B shows that the two individual haplotype assemblies have similar completeness as the previous dd_Smes_g4 assembly (with S3h1 being slightly better than S3h2), yet both display a base-pair accuracy that is at least an order of magnitude higher than that of dd_Smes_g4. Naturally, S3BH showed similar high accuracy but also substantial improvement in completeness, now representing ∼ 98% of the short-read datasets (Fig. 1B). Genome completeness assessment using Benchmarking Universal Single-Copy Orthologs (BUSCO) showed that both S3h1 and S3h2 have higher completeness than dd_Smes_g4 and schMedS2 ^30^ (Additional File 1: Section 1.1). S3h1 again performed slightly better than S3h2, which is why we chose S3h1 as our focal assembly in all following analyses. In contrast, for analyses depending more on reference completeness (e.g., RNA-seq read mapping), we recommend the use of S3BH instead.

To independently assess the long-range contiguity of our assembly, we performed whole genome alignment between our phased assemblies and schMedS2. The result demonstrated a high degree of structural agreement between the two independent scaffolding efforts (Additional File 1: Section 1.2). Additionally, the S3 assembly successfully captured prominent repeat regions on all chromosomes that were absent in previous assemblies, and that likely contributed to the slightly larger size. These repeat regions often reach a length of > 1 Mb as exemplified by our visualization of these regions on Chromosome 4 (Fig. 1C). Closer analyses of the sequence blocks revealed that they are comprised of nested tandem repeats (Fig. 1D) and their successful reconstruction in the S3 assembly is likely a result of the much-increased base-pair accuracy of HiFi reads (Fig. 1B). Comparisons between the two S3 pseudohaplotypes further indicated major structural variations between them. Besides the three inversions on Chromosome 1 (Fig. 1E) that were already described previously ^30^, we detected one inversion on Chromosome 2 and one inversion on Chromosome 4. Heterozygosity was found to be largely restricted to Chromosome 1, again aligning with prior findings ^30^, and on Chromosome 2 partially associated with the inversion but also extending to other regions (Fig 1E). The mapping of the Hi-C data onto the diploid assembly (S3BH) revealed an abundance of uniquely mapping reads to regions with high heterozygosity (inverted region of Chromosome 1 and Chromosome 2), and the central regions of Chromosome 4 that harbor the inversion. The current assembly phasing is therefore likely an accurate representation of the haplotype divergence in these regions (Additional File 1: Section 1.3). By contrast, the much lower density of haplotype differences in other parts of Chromosome 2 and almost the entire Chromosome 3 is likely due to the extensive inbreeding of our genome strain (> 18 generations) and more frequent recombination events in non-inverted regions of the genome ^29,30^.

The distribution of genes across the chromosomes was largely uniform with no typical reduction in gene density toward the centromere (Fig. 1E). A notable exception to this is Chromosome 4 where we found a marked increase in transposon density and a concurrent decrease in gene density at the metacentric chromosome (Fig. 1E). Overall, the S3 phased genome assembly represents significant improvements over previous assemblies in terms of accuracy, completeness, and assembly contiguity and thus a strategic community resource for the analysis of gene function in the model species *S. mediterranea*.

### *Schmidtea mediterranea* genome annotation

We further sought to complement the new genome assembly with high quality gene annotations. Encountering increasingly diminishing returns on investment with our previous *de novo* gene prediction approaches (^42^; data not shown), we instead developed a new hybrid approach that merges Oxford Nanopore long-reads (ONT), Illumina short-reads, and 3P-seq of transcription termination sites (TTS) data ^43^ with genome-guided transcript assembly and thus leverages the benefits of direct gene isoform evidence with the base pair accuracy of our genome assembly (Fig. 1F). We generated separate annotation sets for both haplotypes, S3h1 and S3h2, as well as a combined annotation that contains the transcripts from both haplotypes (S3BH). Further, we subsetted each annotation for high-confidence transcripts/coding genes on basis of Open Reading Frame (ORF) length thresholds and a minimum coverage filter (see Methods). High confidence transcripts/genes are designated by a ‘hconf’ label to facilitate high specificity applications (e.g., promoter analyses), while the unfiltered annotation sets are recommended for applications requiring maximal mappability (e.g., RNA-seq mapping analyses). With a total of 58,739 and 58,551 gene loci and 21,401 and 21,310 high-confidence gene loci in S3h1 and S3h2, respectively, our new annotation sets are broadly in range with the *S. mediterranea* gene number estimates from previous studies ^29,42,44^.

To analyze the overall quality of the new annotations, we carried out systematic benchmarking comparisons against previous transcriptomes or *S. mediterranea* gene model predictions. As a measure for annotation efficiency (i.e. sensitivity), we first assessed BUSCO representation and completeness. S3BH had the highest number of complete BUSCOs (789) and the fewest missing BUSCOs (134) of all the annotations tested. Interestingly, the transcriptome or gene model-based BUSCO scores were consistently better than genome-mode BUSCO assessments (Fig. 1G, Additional File 1: Section 1.1), indicating that the detection method used by BUSCO is sub-optimal for planarian genomes. Moreover, the comparatively high number of “missing” BUSCOs reflects a high proportion of genuine gene losses in planarians (see below) and highlights the need for a group-specific BUSCO set. As a further completeness measure, we analyzed the representation of 1075 *S. mediterranea* genes available from NCBI Genbank. S3BH again demonstrated the highest completeness with 1056 genes represented compared to 1004 in the current community reference, the dd_v6 transcriptome (Fig. 1H). In addition, 3 of the 19 transcripts “missing” in the S3BH annotations were false positives (e.g. mitochondrial transcripts, Additional File 1: Section 1.4), thus yielding an overall annotation completeness better than 98 %. Similarly, when quantifying the mappability of published RNA-seq data sets as a global measure of gene annotation completeness, S3BH also outperformed the existing *S. mediterranea* gene annotations, (Fig. 1I) inclusive of a recent expanded gene annotation sets^39,45^.

To assess the specificity of our gene annotations, we manually inspected and compared the representation of 96 genes amongst the different annotations. The test set consisted of often lowly expressed signaling pathway components and 50% randomly chosen genes to provide an unbiased representation of planarian genes. Each gene in the dataset was scored for the presence or absence in the particular annotation and for commonly encountered gene model mistakes, including truncated, fragmented, frame-shifted, or chimeric transcript predictions (Fig. 1J). Collectively, the S3 annotations scored the highest fraction of error-free gene models, with the annotations containing gene models of all test genes and predicted transcripts with intact ORF representations of 93 out of 91 test genes. The identical scores of the “all” versus “high confidence” S3 annotations reflects the inclusion of all 96 test genes in the “high confidence” category and thus provides an important verification of this annotation subset (see above). Although the S3 predictions still harbor a low proportion of chimeric, truncated, or frame-shifted transcripts (see Discussion and Additional File 1: Section 1.5), they nevertheless represent a significant specificity improvement over the current annotations, e.g. the dd_v6 transcriptome with only 78/96 error-free transcript representations and a significantly higher proportion of fragmented transcripts.

Overall, the S3 annotations therefore represent the most complete and accurate gene annotations of the model species *S*. *mediterranea* to date. On the basis of our results, we recommend using S3h1 for gene loci analyses and the diploid assembly as a mapping reference. The annotations are scheduled for inclusion in ENSEMBL Metazoa and will be incorporated in a future PlanMine update.

### Promoter and enhancers annotation in the *S. mediterranea* genome

The annotation of the cis-regulatory elements is a further critical element in understanding the biology of an organism. To explore and annotate regulatory elements in the S3 genome assembly, we sought to first identify accessible chromatin regions using ATAC-seq. With the high nuclease activity and abundant polysaccharides (mucus components) in planarian tissue extracts as persistent experimental challenges ^29^, we modified an existing Omni ATAC-seq protocol ^46^ to minimize clumping of nuclei and to deplete free and mitochondrial DNA contamination (Additional File 1: Section 2.1, see Methods section). To annotate regions of accessible chromatin, we applied our protocol to whole intact (wt) or x-irradiated individuals (x-ray) of the broadly studied asexual strain of *S. mediterranea* and generated a high-confidence ATAC-seq peak set, only incorporating peaks present in at least three biological replicates of the experiment (Fig. 2A). The distribution of nucleosome-free fragments and progressively fewer mono-, di- and trinucleosomal fragments in our ATAC-seq libraries ^47^(Fig. 2B, top), the typical transcription start site (TSS) enrichment profiles of nucleosomal-free and mono-nucleosomal fragments (Fig. 2B, middle) and the size distribution of the ATAC-seq peaks (Fig. 2B, bottom) collectively indicate the high quality of our ATAC-seq data, as well as TSS annotations of the S3 gene models. Altogether, the merged peak set comprises 55,585 high-confidence peaks with a mean length of 668bp (Fig. 2B, Additional File 2: Table S1).

**Figure 2.**
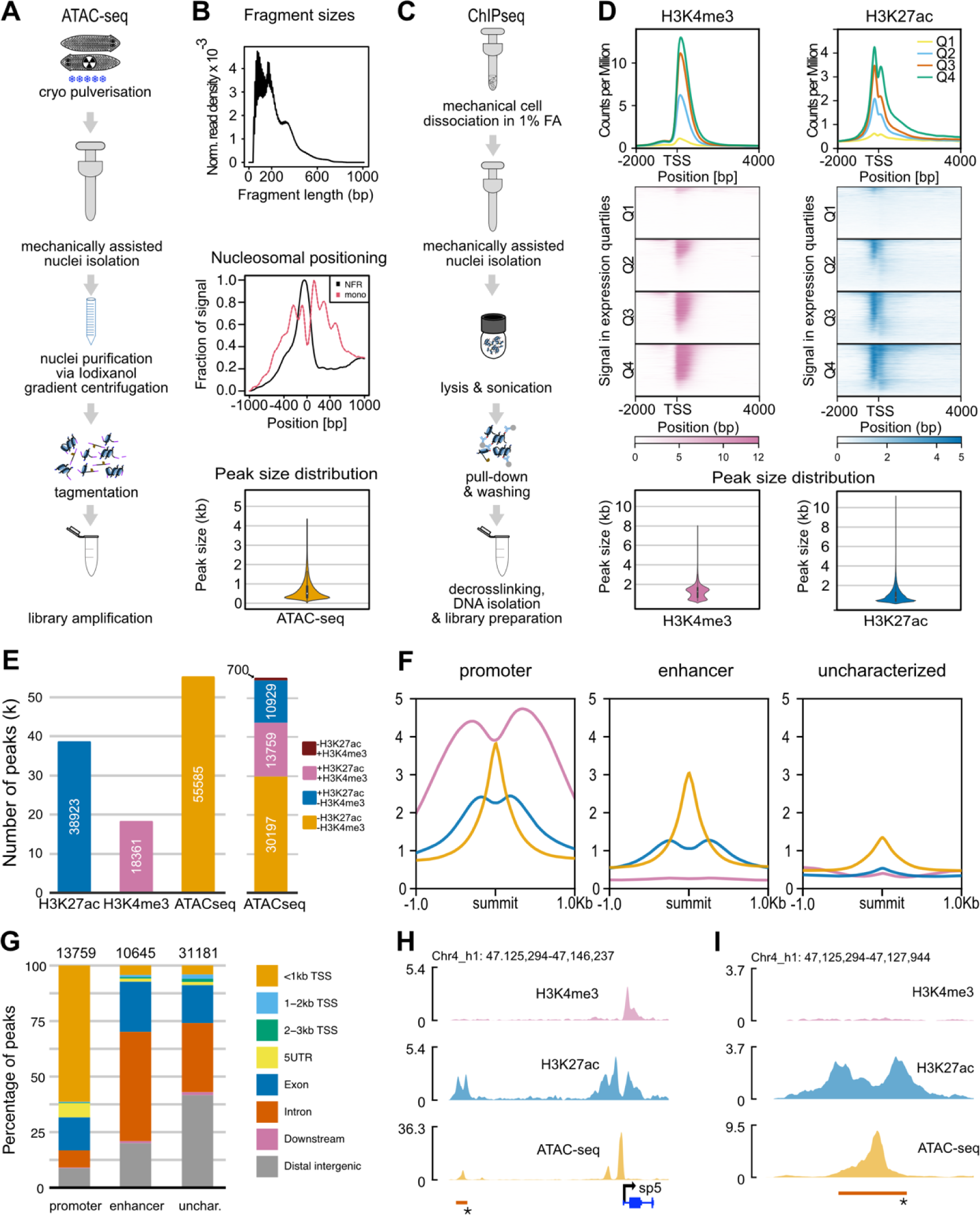
Profiling chromatin regulatory landscape in *Schmidtea mediterranea* using ATAC-seq and ChIP-seq. **A:** Schematic illustrating the workflow used to generate ATAC-seq libraries. **B:** top: Graph depicting a typical fragment size distribution. Increased concentration around 100 and 200 bp are reflecting the presence of nucleosome-free and mono-nucleosome-bound fragments. Middle: Representative TSS enrichment plot displaying an abundance of nucleosome-free fragments at transcription start sites (TSS), while mono-nucleosome fragments are depleted at TSS but enriched at flanking regions. Bottom: Genome-wide peak size distribution of ATAC-seq peaks. **C:** Schematic illustrating the workflow used to generate ChIP-seq libraries. **D:** Top: Average coverage profiles and heat maps of H3K27ac and H3K4me3 ChIP-seq data around TSS of genes with different expression levels showing the correlation between RNA expression quantiles and their correlation with histone modification ChIP-seq signal intensity heatmaps. Bottom: Genome-wide peak size distribution of H3K4me3 and H3K27ac ChIP-seq peaks. **E:** Bar plot displaying the number of peaks called for H3K4me3 and H3K27ac ChIP-seq as well as ATAC-seq (left). Stacked bar plot showing ATAC-seq peak intersection with histone marks (right). **F:** Profile of H3K4me3 and H3K27ac ChIP-seq marks centered on the three classes of ATAC-seq peaks. **G:** Stacked barplots showing the distribution of putative promoters, putative enhancers, and uncharacterized ATAC-seq peaks in relation to the high-confidence gene annotations. **H:** Example of a putative promoter with H3K4me3 and H3K27ac signal at the TSS of the sp5 gene. **I:** Example of a putative promoter (red bar) with H3K4me3 and H3K27ac signal upstream of the sp5 gene.

To further sub-categorize these accessible chromatin regions, we thought to leverage the known enrichment of specific histone marks at gene regulatory sequences (reviewed in ^48^). To do so, we developed a ChIP-seq protocol that utilizes isolated nuclei from fixed tissue (Fig. 2C). As shown in Additional File 2: Section 2.2, our protocol yielded high signal-to-noise ChIP-seq signals with both the H3K4me3 mark (TSS and gene body of actively transcribed genes ^49^) and H3K27ac (enhancers and TSS ^50^). In addition, the levels of H3K27ac and H3K4me3 signal in the vicinity of the TSS correlated with the corresponding gene expression levels as a further indicator of data quality and evolutionary conservation of histone mark distribution in planaria (Fig. 2D, middle). In total, our ChIP-seq data set consists of 18,361 H3K4me3 peaks with a mean size of 1,184 bp and 38,923 H3K27ac peaks with a mean size of 891bp (Fig. 2D bottom, Fig. 2E; Additional File 2: Table S2-S3).

Intersecting our ChIP-seq peak sets with the 55,585 ATAC-seq peaks revealed significant co-occurrence of ATAC-seq signal and both H3K4me3 signal (permutation test; 10,000 permutations, observed overlap: 15,465, permuted overlap: 2,251, Z-score: 15,465, P-value < 0.0001, Fig. 2E) and H3K27ac signal (permutation test; 10,000 permutations, observed overlap: 25,550, permuted overlap: 4,017, Z-score: 25,550, P-value < 0.0001, Fig. 2E). In line with the conserved functions of the examined histone marks, we designated the 13,759 ATAC-seq peaks with associated H3K27ac and H3K4me3 as ‘putative promoters’ and the 10,645 ATAC-seq peaks with only H3K27ac signal as ‘putative enhancers’. The remaining 30,481ATAC-seq peaks without H3K27ac or H3K4me3 signal and 700 ATAC-seq peaks with only H3K4me3 signal were designated ‘uncharacterized accessible chromatin’ (Fig. 2E; see Additional File 2: Tables S1 for details on the classification of each called ATAC-seq peak). Consistent with this functional categorization, we found that ‘putative promoters’ collectively displayed a sharp ATAC-seq peak centered within “valleys” of both H3K4me3 and H3K27ac signals (Fig. 2F, Additional File 1: Section 2.3) and that > 61.4% were located within 1 kb upstream of a TSS annotation (Fig. 2G). In contrast, ‘putative enhancers’ collectively displayed the ATAC-seq peak in a “valley” of surrounding H3K27ac signal (Fig. 2F, Additional File 1: Section 2.3) and 80% were either intronic (49.2%), exonic (22.6%), or otherwise associated with a gene model. As expected, “Uncharacterized” ATAC-peaks lacked histone mark enrichment and displayed generally lower ATAC-seq signals (Fig. 2F).

The fact that only 20% of enhancers were designated as “distal intergenic” indicates that the regulatory elements in the *S. mediterranea* genome may be closely associated with genes. The Smed-sp5 gene locus (Fig. 2H, I) illustrates the above distribution of chromatin mark features, specifically a putative promoter immediately upstream of the TSS and a putative enhancer ∼3 kb upstream of the TSS (Fig. 2I). Overall, the 13,759 putative promoters and 10,645 putative enhancers resulting from our ATAC-seq and ChIP-seq integration likely represent regulatory regions. However, with 21,401 genes in the S3h1 hconf annotations (Additional File 2: Table S4), it is clear that our whole-animal averaged peak annotations only cover a subset of regulatory regions, likely those of constitutively expressed genes or those active in the most abundant cell types.

### Genomes and annotations for *S. polychroa*, *S. nova*, and *S. lugubris*

To orthogonally verify our peak annotations and to address the extent by which the remaining 30,197 ATAC-seq peaks without ChIP marks might represent false negatives, we turned to the principle that the sequence of important regulatory elements is often conserved over evolutionary time ^51^. Our genome sequencing and annotation pipeline made sequencing additional planarian genomes technically feasible. Since multiple lines of evidence indicate unusually high sequence divergence within planarians and between flatworms in general ^32,52–55^ we sequenced and assembled the genomes of the three closest known relatives of *S. mediterranea*. Namely, all known members of the genus, *S. polychroa*, *S. nova*, and *S. lugubris* (Fig. 3A). All sequenced strains were diploid and displayed the expected karyotypes with 3 or 4 chromosomes (not shown; ^56–59^). The Hi-C maps of the assemblies indicated that the genomes are well scaffolded (Fig. 3B) and of similarly high assembly qualities as for *S. mediterranea* (Fig. 1A). In addition, the BUSCO scores (Fig. 3C) suggested a comparable completeness to the *S. mediterranea* S3 assembly (Fig. 1G). Interestingly, the assemblies of *S. nova* (1,251 Mb) and *S. lugubris* (1,499 Mb) were substantially larger than those of *S. mediterranea* (840 Mb) and *S. polychroa* (781 Mb) (Table 1).

**Figure 3.**
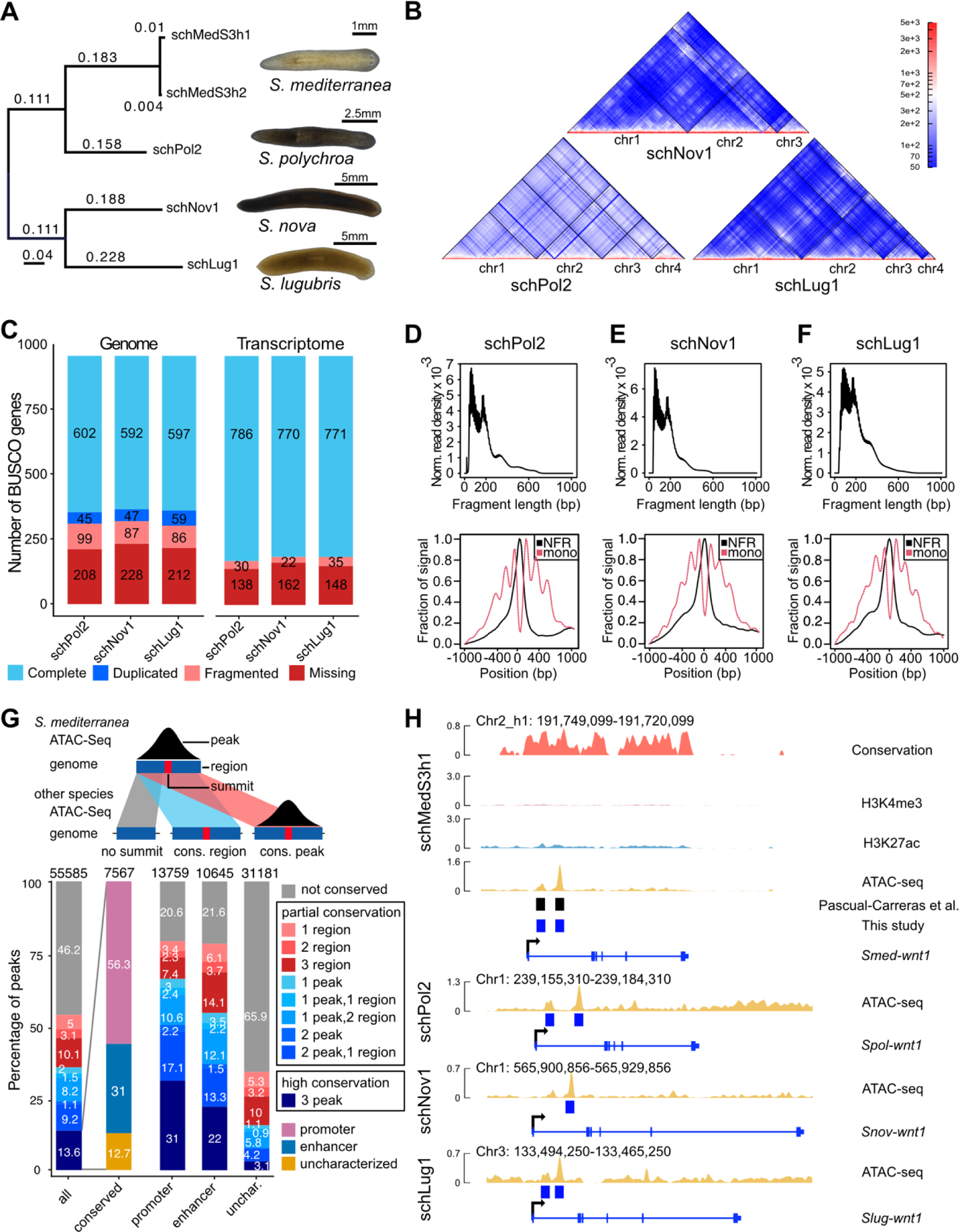
Genome and ATAC-seq quality metric and assessment of regulatory element conservation. **A:** Phylogenetic tree of the genus *Schmidtea* was inferred using four-fold degenerate site positions across our whole genome alignments. Branch lengths indicate expected substitutions per site. **B:** HiC contact maps of the genome assemblies of *S. polychroa* (schPol2), *S. nova* (schNov1), and *S. lugubris* (schLug1). **C:** BUSCO assessment of the genome assemblies (left three bars) and the transcriptome annotations (right bars) of the species in B. Note, that the transcriptome assessment was run on the transcript level, leading to extensive duplication and for visualization Duplicate and Complete were combined. **D-F:** On top the ATAC-seq fragment-size distribution and below the pileup of nucleosomal free (NFR) and mono-nucleosomal (mono) reads at transcription start sites (TSS) of the species in B. **G:** Schematic diagram showing the definition of a conserved region (sequence conservation without ATAC-seq signal) and conserved peak (sequence conservation and ATAC-seq signal). Barplots below indicate the conservation status in *S. mediterranea* of all accessible chromatin and the element category of the highly conserved elements revealing that 87.3% of them are associated with ChIP marks. Three barplots to the right show the conservation status of putative promoters, putative enhancers, and uncharacterized ATAC-seq peaks. **H:** Example of highly conserved regulatory elements at the Wnt1 gene, showing sequence conservation inferred using PhyloP based on the whole genome alignment, as well as, H3K4me3 ChIP, H3K27ac ChIP, and ATAC-seq signal for *S. mediterranea* on the top, followed by a track indicating the location of putative enhancers identified by Pascual-Carreras et al. ^115^. Track below shows the position of the two corresponding putative enhancers we identified and their conservation across the genus *Schmidtea* illustrated by the ATAC-seq signal and called peaks in the genomes of *S. polychroa* (schPol2), *S. nova* (schNov1), and *S. lugubris* (schLug1) shown below. Note, that only a single peak was called in *S. nova* and the peak was therefore classified as a conserved region in that species.

To annotate the new genomes, we again used our hybrid transcriptome assembly strategy (Fig. 1F) yet without 3P-seq TTS evidence and coverage-based “high confidence” filter due to the lack of extensive RNA-seq data for these species. The annotation statistics indicated similar gene numbers and gene length distributions as for *S. mediterranea* (Table 1, Additional File 2: Table S4). Additionally, the BUSCO annotation completeness was improved compared to the run-in genome mode and achieved comparable results to the *S. mediterranea* assembly (Fig. 3C, Additional File 2: Table S5-6). Nevertheless, we identified 125 BUSCOs that are consistently missing across all four high quality genomes and thus provide further illustration of the previously noted substantial gene loss in planarians ^29^ (Additional File 2: Table S5-6). The analysis of four-fold degenerate site divergence revealed considerable sequence divergence between the 4 *Schmidtea* species. *S. polychroa* differed from *S. mediterranea* by 0.3 substitutions per site, a distance analogous to that between humans and horses ^60^. Both *S. nova* and *S. lugubris* show a divergence to *S. mediterranea* of ∼0.6 substitutions at four-fold degenerate site, similar to the distance between humans and shrews ^60^ (Fig. 3A). Overall, our additional high-quality genome assemblies and annotations provide an interesting comparative perspective on the *S. mediterranea* genome, especially because the four assemblies are considerably more distant than what one might expect of close sister species in other taxonomic branches.

### Conservation of regulatory regions

We first explored the conservation of the putative *S. mediterranea* gene regulatory element annotations in the new genomes. Besides sequence conservation alone, we opted to also assess the conservation of chromatin accessibility as a proxy for functional conservation in the different species. We therefore collected ATAC-seq data sets in *S. polychroa*, *S. lugubris* and *S. nova* under the same biological conditions as previously employed for *S. mediterranea* (wt and x-ray, see Methods for further detail). As shown in Fig. 3D, the quality control analysis of all ATAC-seq data in the other *Schmidtea* species confirmed the robust and species-independence of our revised protocol (Fig. 3D-F). Merging of the replicates and biological conditions resulted in 36,729, 60,352, and 77,926 ATAC-seq peaks for *S. polychroa*, *S. nova*, and *S. lugubris*, respectively (Additional File 2: Table S7-9). To assess the conservation of *S. mediterranea* ATAC-seq peaks relative to these data, we assessed sequence conservation under each peak through whole genome alignment liftover, and chromatin state conservation by overlapping the whole genome alignment liftover with ATAC-seq data in the receiving species. Peaks from *S. mediterranea* with only sequence conservation in the receiving species were designated as conserved regions. In contrast, conserved regions that additionally overlapped with an ATAC-seq peak in the receiving species were designated as conserved peaks (Fig. 3G). This allowed us to categorize each *S. mediterranea* peak according to its conservation status across the genus with the most informative categories being ‘not conserved’, ‘partial conservation’ (all categories except not conserved) and ‘high conservation’ (3 peaks, Fig. 3G, Additional File 2: Table S1), indicating peaks that showed conservation in all three recipient species. Across all 55,585 ATAC-seq peaks annotated in *S. mediterranea*, 13.6% were highly conserved, 40.2% partially conserved and 46.2% were not conserved (Fig. 3G). Interestingly, 87.3% of the highly conserved ATAC-seq peaks also had additional ChIP-seq support (Fig. 3G). Furthermore, when considering all ATAC-seq peaks with additional ChIP-seq support, we found that 79.4% of the putative promoters and 78.4 % of the putative enhancers displayed at least partial conservation, thus confirming that these peak sets are indeed likely to be enriched for functionally relevant regions under purifying selection (Additional File 1: Section 3). As expected, the 30,197 “uncharacterized” ATAC-seq peaks were collectively much less conserved, with only 3.1% of high conservation and 34.1% of partial conservation (Chi-squared = 9745.1, df = 4, p-value < 2.2e-16, all post-hoc tests p < 0.001, Additional File 1: Section 3). Although the partially conserved sub-set might, therefore, indeed include functionally relevant sequences, the nature and functional significance of the majority of *Schmidtea*-specific ATAC-only peaks remains unclear. In line with the considerable sequence divergence between the four *Schmidtea* species, the ∼20% ATAC-seq peaks that are *S. mediterranea* specific but ChIP-annotated further highlight a significant degree of regulatory divergence within the genus.

Owing to the comparatively recent advent of functional genomics in the field, only two gene-enhancer sequences have been partially characterized so far ^40,61^. To gauge the practical utility of our regulatory element analysis, we first examined the putative enhancers of *S. mediterranea Wnt1* identified by ^40^. As shown in Fig. 3H, we also identify two ATAC-seq peaks in the first intron, which we categorize as putative intronic enhancers due to their overlap with the H3K27Ac signal. In addition, our conservation analysis identifies one region as “highly conserved”, with prominent ATAC-seq peaks in all species, and one region as having a conserved peak in *S. polychroa* and *S. lugubris*, but only a conserved region in *S. nova* (Fig. 3H). Interestingly, the prominence of the ATAC-seq peaks in all four species and the H3K27ac peak in *S. mediterranea* contrast with low H3K4me3 signals at the *S. mediterranea* TSS (Fig. 3H, top). While the latter is consistent with the highly specific expression of *Wnt1* in very few cells at the tail tip of intact animals, the former might indicate that the regulatory regions of *Wnt1* are constitutively “poised” in a much broader range of cells in order to allow its dramatic upregulation at any *S. mediterranea* wound site ^62^. In contrast, the proposed head-specific enhancer sequence of *nou-darake* (ndk) was not detected by our analysis, even though multiple other putative regulatory sequences in the vicinity of the gene were annotated (Additional File 1: Section 3). While this might well reflect the limitation of our current whole animal data sets with respect to detecting regulatory regions that are only active in a small number of cells, our comparative genomics approach adds a valuable layer of information to the *S. mediterranea* genome annotations (Additional File 2: Table S1). By summarizing both the ChIP-seq and conservation status of all *S. mediterranea* ATAC-peaks, Table S1 represents a valuable resource for the reconstruction of regulatory circuits in the model species and their possible evolutionary divergence across the genus *Schmidtea*.

### Genome architecture & synteny

The availability of the four chromosome-scale genome assemblies also provided a first opportunity for exploring other features of genome evolution within the taxon. As noted before, *S. lugubris* and *S. nova* had substantially larger genomes than *S. polychroa* and *S. mediterranea* (Fig. 4A). Transposable element annotations revealed that a large proportion of the increase in genome size can be attributed to an expansion of transposable elements, in particular, DNA and LTR/Gypsy elements (Fig. 4A). Furthermore, in *S. lugubris* and *S. nova*, the total gene span was 54% and 28% larger compared to *S. mediterranea*. This increase was primarily due to the increased length of protein-coding genes and, specifically, the expansion of introns, at least in parts due to transposon insertions (Table 1, Additional File 2: Table S4). Therefore, the larger assembly sizes of *S. lugubris* and *S. nova* reflect genuine expansions of genome size due to transposable element expansions. The elucidation of the specific transposon families that mediated the expansions will be an interesting topic for future investigations.

**Figure 4.**
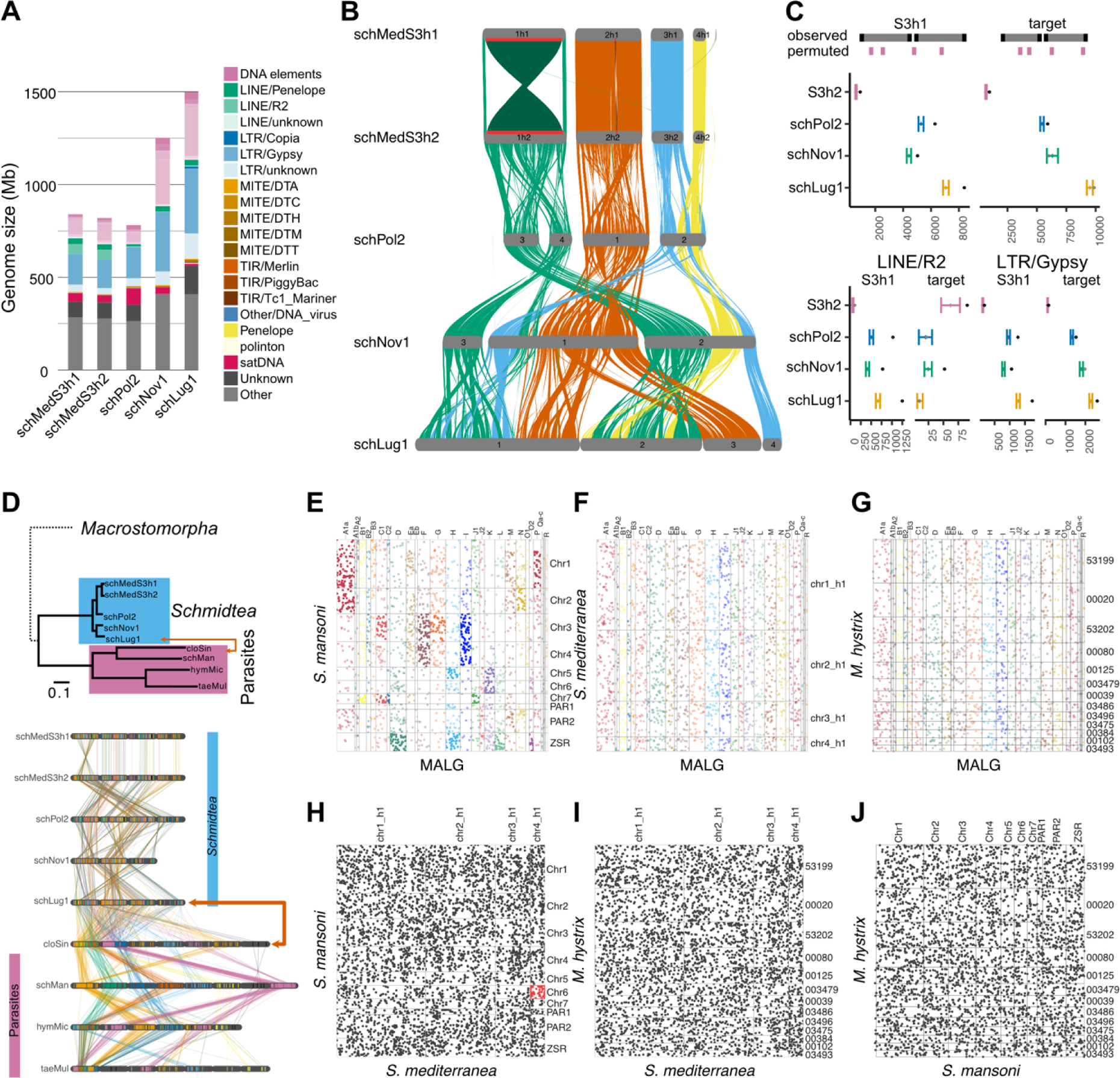
Genome content and synteny analysis of *Schmidtea*, parasite and a Macrostomorph genome. **A:** Genome assembly size for all four *Schmidtea*. Barplot colors indicate the proportion of the genome that is made up of various types of repetitive elements. **B:** Gene order based synteny analysis between four *Schmidtea* species. Ribbons are colored based on the chromosome location in *S. mediterranea*. Red bars and darker shading in schMedS3h1 indicate the inversions on Chromosome 1 and 2 that distinguish haplotype 1 and 2. **C:** Enrichment analysis of 10kb windows flanking synteny breakpoints inferred using GENESPACE. Tests are based on comparisons of the observed value (black dots) to 1000 permutations of random placement of an equal number of 10kb windows in the reference (colored mean and standard deviations). Black dots outside the permuted range indicate statistically significant enrichment. Shown on top are results for all transposable elements followed by LINE/R2 and LTR/Gypsy elements below since they returned the most relevant results. For a full table including all tested elements, see Additional File 1: Section 4.2 and Additional File 2: Table S12. **D:** On top the phylogenetic relationship between *Schmidtea* and the parasites based on protein divergence. The early-branching Macrostomorpha are indicated with a dotted line. Below a synteny analysis between *Schmidtea* and four species of parasitic flatworms. BUSCO genes are represented with bars colored based on the chromosome location in *Schistosoma mansoni*. **E-G:** Dotplots between metazoan ancestral linkage groups (MALG) and (E) *Schistosoma mansoni*, (F) *Schmidtea mediterranea*, and (G) *Macrostomum hystrix*. MALGs that are significantly enriched in one or more chromosomes according to a Fisher Exact test are indicated in dark color, while MALGs without enrichment are plotted in light colors. No enrichment was detected for any MALG in *S. mediterranea* and *M. hystrix*. **H-J:** Synteny analysis between *Schistosoma mansoni*, *Schmidtea mediterranea*, and *Macrostomum hystrix*. Boxes indicate chromosome combinations and dots represent one-to-one orthologs inferred with ODP. No chromosome combination was enriched except for 57 orthologs between *S. mansoni* and *S. mediterranea* (shown in red).

Next, we assessed the synteny between the four genomes using GENESPACE ^63^. The visualization of the syntenic blocks revealed a large number of rearrangements between the genomes (Fig. 4B). Already between the two pseudo-haplotypes of *S*. *mediterranea*, the previously noted large inversion on Chromosome 1 and the smaller inversion on Chromosome 2 stand out as prominent structural rearrangements (indicated with red bars and dark shading in Fig. 4B). Comparisons with the other genomes revealed a striking history of frequent structural rearrangements encompassing inversions and inter-chromosomal translocations. For instance, the gene content of *S. mediterranea*’s Chromosome 1 equates to *S. polychroa*’s chromosomes 3 and 4, and *S. polychroa*’s Chromosome 2 is equivalent to *S. mediterranea*’s chromosomes 3 and 4, suggesting splits or fusions between all the involved chromosomes (Fig. 4B). Moreover, the chromosomal reduction in *S. nova* from 4 to 3 implies a complex fusion event between multiple chromosomes. The rapidly decreasing size and increasing number of syntenic blocks in the assembly comparisons quantitatively confirmed the chromosomal fragmentation apparent in Fig. 4B (See Additional File 1: Section 4.1 for more details and Additional File 2: Table S14 for the inferred syntenic blocks). Interestingly, we found that 10kb windows flanking the synteny breakpoints in our species panel were significantly enriched in LTR/Gypsy retrotransposons in all assemblies except for *S. nova*, and enriched in LINE/R2 retrotransposons in all assemblies except *S. nova* and *S. lugubris* (Fig. 4C), thus providing a first indication that transposable elements may play a role in the frequent structural rearrangements as is the case for LINE elements in humans ^64^ (Additional File 1: Section 4.2 and Additional File 2: Table S15). Overall, our synteny analysis therefore revealed a surprising amount of structural genome rearrangements within the taxon *Schmidtea*, including frequent inter-chromosomal translocations and the consequent erosion of gene order within chromosomes.

Intrigued by these findings, we next asked whether this feature is unique to *Schmidtea* or if synteny is generally poorly conserved amongst flatworms. To address this, we selected chromosome-scale genome assemblies of four parasitic flatworms (Neodermata; comprising the two Trematoda species *Clonorchis sinensis* and *Schistosoma mansoni* and two Cestoda species *Taenia multiceps* and *Hymenolepis microstoma*) that we collectively refer to as “parasites” in the following. Additionally, we included a highly contiguous – but not chromosome-scale – genome assembly of *Macrostomum hystrix* in some of the following analyses, as a representative of the early branching flatworm group Macrostomorpha. As expected on basis of the large evolutionary distances involved, the protein-sequence divergence of the parasite genomes was much higher than the divergence within the *Schmidtea* taxon (top, Fig. 4D). To obtain an overview of synteny conservation across such evolutionary distances, we first compared the chromosome-scale assemblies using the location of single-copy genes on basis of *de novo* BUSCO annotations. As shown in Fig. 4D, large genomic blocks of BUSCOs (color blocks; parallel lines) were clearly conserved across the parasites, despite several large-scale genomic rearrangements. Intriguingly, between the parasites and the *Schmidtea* genomes (red arrows), the relative positions of the same BUSCO genes appeared largely randomized. Of note, the loss of synteny also includes the BUSCO content of the parasite sex chromosome, which was previously hypothesized to be predominantly homologous to Chromosome 1 of *S. mediterranea* ^30^. We quantitatively confirmed this finding using a 9-fold expanded gene set using Orthofinder and Chi-squared test analyses, again finding the maintenance of measurable synteny within the *Schmidteas* and the parasites, but the near-total degradation of synteny between the two groups (e.g., small residual effect sizes with all <0.09, and with the Chi-square test not even reaching the significance threshold for the schMedS3h1-hymMic and schMan-schNov1 combinations; Additional File 1: Section 5.3). The analysis of chromosome gene complement conservation between *Schmidtea*, the parasites, and the early-branching flatworm *Macrostomum hystrix* using 1:1 orthologue annotations inferred using the ODP tool (see Methods) also revealed the near complete absence of synteny between the two groups (Additional File 1: Section 5.4). Overall, this analysis therefore confirmed a profound loss of synteny and chromosomal gene content between the genus *Schmidtea*, parasitic flatworms and the early branching flatworm *Macrostomum hystrix*.

This result also raised the question which of the flatworm groups better conformed to the 28 ancestral metazoan linkage groups identified by Simakov et al. ^8^, which we here refer to as Metazoan Ancestral Linkage Groups (MALG). We, therefore, identified orthologs of the MALG using the ODP tool to determine if they were associated with particular chromosomes of our test genomes. Half of the 28 MALGs were statistically significantly enriched in *Schistosoma mansoni* on specific chromosomes (Fig. 4E, Additional File 1: Section 4.5). However, all linkage groups were either split between several chromosomes and/or fused and mixed with other linkage groups (See our detailed description of these mixing events in Additional File 1: Section 4.5). A similar pattern emerged in the other parasites with 12, 6, and 12 MALGs partially conserved in *Clonorchis sinensis*, *Hymenolepis microstoma*, and *Taenia multiceps*, respectively. Notably, despite the conservation of synteny between the parasites, the statistical tests did not find conservation of the same MALGs in each species (Additional File 1: Section 4.5). This suggests that the parasites retain detectable traces of the MALG, even though they have been largely obscured by a history of pervasive chromosomal rearrangements. In sharp contrast, we failed to detect any traces of MALG conservation across the *Schmidtea* genomes (Fig. 4F, Additional File 1: Section 4.5). MALGs were similarly undetectable in the genome assembly of *Macrostomum hystrix* (Fig. 4G, Additional File 1: Section 4.5). Intriguingly, our ODP analysis further revealed the lack of synteny conservation between *Macrostomum*, *Schmidtea*, and the parasites, therefore suggesting that these taxa represent entirely independent genome architectures (Fig. 4H-J, Additional File 1: Section 4.5).

Overall, our data reveal gene shuffling between the different flatworm groups, indicating that synteny may not pose a functional constraint in the evolution of flatworm genomes.

## Discussion

Here, we present and analyze four high-quality genomes of planarians flatworms in the genus *Schmidtea*, including a richly annotated chromosome-scale and haplotype phased assembly of the model species *S. mediterranea*. The new S3 assembly significantly improves over our previous *S. mediterranea* assembly ^29^ in terms of assembly contiguity, completeness, and sequence accuracy (See Fig. 1A-B). The haplotype-phasing of the assembly (Fig. 1A,E) will benefit many research applications in the model species such as primer design, variant calling, motif enrichment analysis, or the design of guide RNAs for the development of Cas9 genome modifications as one of the current frontiers in the field. In addition, the increased accuracy of the HiFi reads over previous sequencing chemistries resolved many tandem repeat stretches in the *S. mediterranea* genome that can reach > 1 Mb in length and that were collapsed in the previous assembly (Fig. 1C-D; Table 1). Similarly relevant for the planarian research community are the S3 genome gene annotations, which represent a significant improvement over existing *S. mediterranea* gene and protein predictions (Fig. 1G-J), including the dd_v6 transcriptome that is currently used as a reference by many labs in the community ^42^. Furthermore, we provide a global annotation of regulatory regions in the *S. mediterranea* genome out of the intersection between ATAC-seq and ChIP-seq data that complements similar recent efforts ^39–41^. The additional dimension of evolutionary sequence conservation in the three *S. mediterranea* sister species provides a valuable tool for motif conservation analysis or the prioritization of suspected regulatory regions for functional analysis. Altogether, this amounts to a valuable starting point for the reconstruction of gene regulatory circuits in *S. mediterranea*.

Even though the S3 assembly and associated annotation tracks significantly expand the available genome resources for the planarian research community, it is important to stress that the *S. mediterranea* genome resources remain a work in progress. Remaining challenges include missing telomeres on some chromosome arms of the S3 assembly and ∼400 unincorporated contigs containing repeats, telomere fragments, and about 500 annotated genes. Like the assembly, the S3 gene annotations still require refinement. One persistent error category are so-called proximity chimeras, e.g., fusions between closely spaced genes. One such example is the erroneous fusion between the Activin inhibitor *follistatin* to the gene immediately upstream (Additional File 1: Section 1.5), which is likely caused by <500 bp distance between the stop codon of the upstream gene and the TSS of *follistatin* and frequently overlapping transcripts of both genes (not shown). Similarly, the isoforms in the current annotations are uncurated and should be interpreted as indicators of splicing diversity only. Further improvements of the genome assembly and the gene annotations will increasingly depend on manual curation efforts. Further improvements of the genome assembly and gene annotations, as well as, the integration of the increasing number of regulatory element annotations ^39–41^ will increasingly depend on manual curation efforts. The integration of manual curation into PlanMine ^42^ and other community resources (e.g. ^65–68^) therefore represents a strategic objective for the community.

Beyond *S. mediterranea* resources, the four high-quality *Schmidtea* genomes in our study present a first opportunity to explore patterns of genome evolution within the genus and in planarians in general. Consistent with previous transcriptome-based analyses ^32^ the four genomes emphasize the extent of sequence divergence within planarians. Even though we sequenced the *S. mediterranea* sister species *S. polychroa* and its two closest known relatives, the genome assemblies revealed third base divergences equivalent to ∼30 Mio years (*S. mediterranea* vs. *S. polychroa*) or ∼70 Mio years (*S. mediterranea* vs. *S. nova*) of vertebrate genome evolution ^60^. The calibration of sequence divergence rates remains challenging due to the extremely poor planarian fossil record ^69^ and we, therefore, cannot distinguish between ancient divergence or much increased rates of molecular evolution in planarians as the explanation of the sequence divergence between the *Schmidtea* species. However, as discussed below, our analyses collectively favor the latter interpretation.

The numerous inversions, translocations and chromosome fusion and fission events that our genome comparisons revealed provide striking examples of the structural divergence between the *Schmidtea* genomes. Already apparent between the two haplotypes of the S3 assembly (Fig. 1E), large-scale structural rearrangements also dominate the *Schmidtea* species genome comparisons. In fact, the unbalanced chromosomal translocation between the sexual and asexual biotypes of *S. mediterranea* ^70^ and previous taxonomic classification of *Schmidtea* into biotypes based on karyotypes ^59^ have already hinted at frequent structural rearrangements in *Schmidtea* species. Our chromosome-scale assemblies confirm the hypothesis of Benazzi and Puccinelli ^58^, that Chromosome 1 of *S. nova* resulted from a fusion of chromosomes 1 and 3 of *S. lugubris*, but we additionally show that *S. lugubris* Chromosome 1 also shares synteny with Chromosome 3 of *S. nova* and that the gene complement of *Schmidtea* chromosomes is generally unstable. These multiple ‘hidden’ rearrangements are similar to what has recently been uncovered in some holocentric lineages, i.e., Lepidoptera and beaksedges ^71^. However, unlike in the Lepidoptera, we discovered an enrichment of the abundant LTR/Gypsy elements and LINE/R2 elements near the synteny breakpoints (Additional File 1: Section 5.2). This finding is consistent with recent results in plant genomes (beaksedges ^72^) and may hint at the underlying mechanisms for the rearrangements (see below). The consequence of the large scale and high frequency of structural rearrangements in the genomes is a striking degradation of synteny across the *Schmidtea* genomes. For example, we find that synteny between *S. mediterranea* and *S. lugubris* is fragmented into 272 identified syntenic blocks with a median size of 1.5 Mb and covering only 48.7% of the assembly (Additional File 1: Section 4.1; Additional File 2: Table S14). Altogether, these data indicate a substantial loss of synteny within the taxon.

This result also raises the question of whether the structural genome instability is specific to *Schmidtea* versus a general feature of flatworm genome evolution. The striking qualitative randomization of BUSCO genomic positions (Fig. 4D) and the quantitative analysis approach of Metazoan ancestral linkage groups (MALG; (Fig. 4H-J) ^8^) demonstrates the near-complete absence of genomic synteny between *Schmidtea*, the analyzed parasite genomes and even the genome of the early-branching flatworm *Macrostomum hystrix*. These results imply that the genome architectures of the sampled flatworm clades have evolved largely independently, which also raises concerns_regarding the presumptive synteny between the parasite sex chromosomes and chromosome 1 of *S. mediterranea* ^30^. Moreover, our finding that MALGs have been lost independently in *Schmidtea* and *M. hystrix*, but weakly retained in the parasites (also see ^73^) is remarkable for several reasons. The remnants of MALGs only in the parasitic flatworms is unexpected, given that they are often assumed to represent the most derived taxonomic group ^74^ on basis of their phylogenetic position ^52,53^, compacted genomes ^73^ and obligate parasitic life cycles ^75^. A broader taxon sampling of flatworm genomes will therefore be required to place the evolution of parasitism within the evolutionary history of the phylum. In addition, the loss of MALGs *per se* is uncommon, given that MALGs are defined on basis of their conservation across metazoan genomes ^8,9,76,77^ and some are even conserved in unicellular relatives of animals ^10^. Although, MALG losses have already been noted in other taxonomic groups ^11,12,78^, the obliteration of ancestral linkage groups at the base of these lineages appears to have been followed by the establishment and retention of new clade-specific linkage groups (e.g. Nigon elements in Nematodes ^12^, Muller elements in Drosophilids ^11^, ALGs in Bryozoa ^78^). Interestingly, the group-specific linkage groups remain even in clades that contain species with drastic genome/karyotype rearrangements, suggesting selection for the maintenance of linkage groups rather than mechanistic constraints on inter-chromosomal rearrangements ^10^. In contrast, our results indicate that the dispersal of MALGs in flatworms was not accompanied by the emergence of clade-specific linkage groups and that gene order has been and continues to evolve independently within the different taxa. Altogether, this amounts to the provocative proposition that synteny may not matter in flatworms.

The importance of topological constraints on gene expression in many systems ^15–21^ might imply that planarians and other flatworms achieve gene expression specificity by different means. Consistently, topologically associated domains (TADs) are not apparent in our initial Hi-C analysis (Fig. 1A, Fig. 3B) and the finding that 80% of our enhancer annotations in the *S. mediterranea* genome are located in introns, exons, or within 1 kb of the closest gene body (Fig. 2G, Additional File 1: Section 2.3), may reflect a particularly tight association of regulatory elements with their target genes (a genome architecture similar to a tunicate species complex with rapid chromosome-arm restricted gene shuffling ^79^). However, cell or tissue type-specific Hi-C data sets and quantitative comparisons of enhancer distributions with well-annotated genomes will be required to understand the extend by which these observation reflect technical artefacts versus genuine mechanistic differences in the control of gene expression in planarians.

A further interesting question raised by our study are the mechanistic causes of the frequent genome rearrangements in *Schmidtea*. Both free-living and parasitic flatworms survive gamma-irradiation doses well beyond lethal levels in vertebrates ^80–83^, which implies the existence of efficient double-strand break repair pathways that are also known to mediate Robertsonian translocations ^84–86^. The above-mentioned enrichment of LTR/Gypsy and LINE/R2 elements near the synteny breakpoints that we discovered (Additional File 1: Section 5.2), might indicate a role of these abundant retrotransposons as templates for strand invasion during the repair of double-strand breaks. Whether double-strand break repair pathways mediate the frequent chromosomal rearrangements and ultimately drive the likely rapid structural evolution of planarian genomes is therefore a further interesting topic for future analysis. Finally, the possibility that parallel somatic evolution and selection phenomena amongst the many thousand pluripotent somatic stem cells of a single planarian might contribute to the extraordinary rates of sequence divergence raises profound questions regarding the maintenance of genetic “self” and the evolution of multicellularity ^27^. In summary, understanding the mechanistic links between the unusual patterns of genome evolution of flatworms and their unusual biology is a worthwhile research endeavor.

## Methods

### Samples

All animals used for these analyses were derived from long-term laboratory cultures maintained at the Max Planck Institute of Molecular Cell Biology and Genetics in Dresden and the Max Planck Institute for Multidisciplinary Sciences in Göttingen. The animals were maintained in planarian water supplemented with gentamycin sulfate at 50µg/ml at 20C and fed with organic calve liver as described previously ^87^. We used the laboratory strain of the sexual biotype of *S. mediterranea* originating from Sardinia that was also used for the previous genome project (S2F17, derived from S2F2, internal ID: GOE00500). Functional data was generated from the standard laboratory strain of the asexual biotype of *S. mediterranea* (CIW4, internal ID: GOE00071). The *S. nova* strain (internal ID: GOE00023) was collected at 51,0717710; 13,7421400, in Dresden, Germany on 2013-04-14. The *S. lugubris* strain (internal ID: GOE00057) was collected at 52.942432, -1.113739 in Nottingham, UK (JCR). The S. *polychroa* strain (internal ID: GOE00227) was collected at 43.71249; 16.72605 near the Village of Gala, Croatia. Animals were starved for 10 days prior to experiments. For the x-ray irradiation treatment, animals were irradiated with 60 Gray using a Precision Cellrad Cell Irradiation System (10-130 KV, Precision X-Ray, USA).

### DNA extraction

High molecular weight DNA was extracted as previously described with modifications ^29^. Briefly, planarians were treated with a 0.5 w/v N-acetyl-L-cysteine (NAC) stripping solution, augmented with 20 mM HEPES-NaOH at a pH of 7.25. The pH was carefully adjusted to approximately 7 using 1 M NaOH and monitored using a 0.5% w/v phenol-red solution. Planarians were submerged in 10 ml of this freshly prepared NAC solution and agitated vigorously, for instance, on a rotator, for 10-15 minutes at room temperature. After this, a quick rinse with distilled water was done before proceeding to the DNA extraction phase.

For DNA isolation, only wide-bore pipette tips were utilized to handle the high molecular weight DNA, ensuring minimal shear forces. About 20 mucus-stripped planarians, roughly 1 cm in size and having been starved for 1-3 weeks, were placed into a 50 ml tube. They were then lysed using 15 ml of cold GTC buffer (containing 4 M guanidinium thiocyanate, 25 mM sodium citrate, 0.5% w/v N-Lauroylsarcosine, and 7% v/v β-mercaptoethanol) for 30 minutes on ice, with the tube being inverted every 10 minutes to promote tissue dissociation. The lysate was mixed with an equal volume of phenol/chloroform/isoamyl alcohol (in a 25:24:1 ratio), buffered with 10 mM Tris (pH 8.0) and 1 mM EDTA. This mixture was centrifuged at 4,000 x g for 20 minutes at 4°C, after which the upper aqueous phase was carefully collected into a new tube. The phenol/chloroform extraction step was repeated 1-2 times, or until the interphase vanished. Any remaining phenol was removed by mixing the aqueous phase once with an equivalent volume of chloroform, followed by centrifugation. To this cleared aqueous phase, an equal volume of ice-cold 5 M NaCl was added and mixed. After a 15-minute incubation on ice, the sample was centrifuged at maximum speed for 10 minutes at 4°C to pellet any contaminants. The nucleic acid-rich supernatant was moved to a new tube, precipitated using 0.7-1 volumes of isopropanol, and centrifuged at 2,000 x g for 30-45 minutes at room temperature. The DNA pellet was then washed with 70% ethanol, centrifuged at 2,000 x g for 5 minutes, briefly air-dried, and finally resuspended in 50 μl TE buffer and left to dissolve overnight at 4°C.

During post-purification with CTAB at room temperature, contaminants were removed from the isolated DNA. The DNA was treated with RNase A (4 mg/ml) for an hour at 37°C, and NaCl concentration was adjusted using a 2% CTAB/1.4 M NaCl solution. After mixing with chloroform and centrifuging at 12,000–16,000 x g for 15 minutes, the clear phase was extracted. The DNA was then precipitated using isopropanol, washed in 70% ethanol, and resuspended in TE buffer overnight at 4°C.

It is known that DNA can be removed from crude lysates by streptomycin complexation ^88,89^. Compared to other aminoglycoside antibiotics, streptomycin has negligible affinity for acidic mucopolysaccharides, proteins and RNA ^90^. We therefore employed streptomycin precipitation to specifically separate DNA from remaining contaminants. CTAB-purified DNA samples in a 1.5 ml low DNA binding tube (Eppendorf) were mixed with 0.1-0.2 volumes of 50 mg/ml streptomycin sulfate in nuclease-free H2O. For highly viscous samples, the sample was diluted by additional TE buffer before streptomycin addition. The mixture was carefully shaken or flicked to avoid DNA shearing. The DNA-streptomycin complex was allowed to form for at least 15 min at RT and was subsequently precipitated by centrifugation at 4,000xg for 30 min at RT. The supernatant was removed without disturbing the pellet. Excess streptomycin was washed off using 1 ml of PEG/NaCl-based wash buffer (10% (w/v) PEG-8000, 1.25 M NaCl, 10 mM Tris-HCl pH 8.0, 1 mM Na2EDTA pH 8.0, 0.05% (v/v) Tween-20) for 15 min at RT which keeps the DNA precipitated ^91^. After centrifugation for 5 min at 4,000 x g at RT, the supernatant was removed, the pellet briefly washed with 1 ml of 70% ethanol and pelleted as before. After removal of 70% ethanol the pellet was re-suspended in 100 µl of DNA pre-dialysis buffer (10 mM Tris-HCl pH 9.0, 2 M NaCl, 1 mM Na2EDTA, pH 9.0). A dialysis membrane (Millipore: VSWP 04700 (mean pore size = 0.1 μm)) was hydrated on a 100-fold volume of DNA dialysis buffer (10 mM Tris-HCl, pH 9.0, 0.1 mM Na2EDTA, pH 9.0). The sample was carefully transferred onto the dialysis membrane and dialyzed for 4-6 h at RT and then carefully collected using a wide-bore pipette tip. The quality and quantity of the DNA were verified using pulse field gel electrophoresis run using the Pippin PulseTM device (SAGE Science), and the Qubit™ fluorometer.

### PacBio High Fidelity (HiFi) library preparation and sequencing

For all species the genomic DNA entered library preparation using the PacBio HiFi library preparation protocol “Preparing HiFi Libraries from Low DNA Input Using SMRTbell Express Template Prep Kit 2.0”. Briefly, all gDNA was sheared to 14 to 22 kb with the MegaRuptor device (Diagenode) and 12 to 18 µg sheared gDNA was used for library preparation. Depending on the gDNA input amount and performance during library preparation, the PacBio libraries were either size selected for fragments either larger than 3 kb with Ampure beads or for fragments larger than 8 to 10 kb using the BluePippin™ device. The size-selected libraries were prepared for loading following the instructions generated by the SMRT Link software (PacBio, version 10) and the ‘HiFi Reads’ application. The Sequel^®^ II Binding Kit 2.2 (PacBio, USA) was used to prepare the libraries for loading, using the Sequel^®^ II DNA Internal Control Complex 1.0 (PacBio). All libraries ran on SMRT™ Cells 8 M (PacBio) using the Sequel^®^ II Sequencing Kit 2.0 (PacBio) on the Sequel^®^ II Sequencer (PacBio).

### Phased genome assembly of *S. mediterranea*

Circular consensus sequences from ∼30x coverage PacBio reads were called using pbccs (v6.0.0) and reads with quality > 0.99 (Q20) were taken forward as “HiFi” reads. Additionally, we generated 1,000 Million Hi-C reads from extracted nuclei of whole animals using the Arima-HiC+ Kit. PacBio HiFi and Hi-C reads were used to assemble phased contigs with hifiasm v0.7. Next, Hi-C reads whose mapping quality no less than 10 (-q 10) were further utilized to scaffold the contigs from each haplotype by SALSA v2) following the hic-pipeline (https://github.com/esrice/hic-pipeline), which includes filtering procedures such as removal of experimental artifacts from Hi-C alignments, fixing of Hi-C pair mates, and removal of PCR duplicates, etc. Four chromosome-level scaffolds could be observed in both haplotypes after scaffolding. However, Hi-C heatmap revealed evidence of misplacement of contigs in terms of positions and orientations. These errors were then manually curated based on the interaction frequency indicated by the intensity of Hi-C signals.

### Genome assembly of *S. polychroa*, *S. nova*, and *S. lugubris*

Circular consensus sequences from PacBio reads were called using pbccs v(6.0.0) and reads with quality > 0.99 (Q20) were taken forward as “HiFi” reads. To create the initial contig assemblies for Schmidtea nova, canu v2.1 was used with parameters: maxInputCoverage=100 -pacbio-hifi. For Schmidtea polychroa and Schmidtea lugubris hifiasm (v0.14.2) was used to create initial contigs with purging parameter: -l 2. Next, alternative haplotigs were then removed using purge-dups (v1.2.3) using default parameters and cutoff as they were correctly estimated by the program. To initially scaffold the contigs into scaffolds, SALSA v2 (v2.2) was used after mapping HiC reads to the contigs. The VGP Arima mapping pipeline was followed: https://github.com/VGP/vgp-assembly/tree/master/pipeline/salsa using bwa-mem (v0.7.17), samtools (v0.10) and Picard (v2.22.6). False joins in the scaffolds were then broken and missed joins merged manually following the processing of HiC reads with pairtools (v0.3.0) and visualization matrices created with cooler (v0.8.11).

Following scaffolding, the original PacBio subreads were mapped to the chromosomes using pbmm2 (v1.3.0) with arguments: --preset SUBREAD -N 1 and regions +/- 2kb around each gap were polished using gcpp’s arrow algorithm (v1.9.0). Those regions in which gaps were closed and polished with all capital nucleotides (gcpp’s internal high confidence threshold) were then inserted into the assemblies as closed gaps.

Lastly, the PacBio HiFi (CCS reads with a read quality exceeding 0.99) were aligned to the genomes using pbmm2 (v1.3.0) with the arguments --preset CCS -N 1. DeepVariant (v1.2.0) was used to detect variants in the alignments to the assembled sequence. Only the homozygous variants (GT=1/1) that passed DeepVariant’s internal filter (FILTER=PASS) were retained using bcftools view (v1.12) and htslib (v1.11). The genome was then polished by creating a consensus sequence based on this filtered VCF file, as detailed in the VGP assembly pipeline (https://github.com/VGP/vgp-assembly/tree/master/pipeline/freebayes-polish).

### Bacterial and mitochondrial sequence removal

We used FCS-GX (https://github.com/ncbi/fcs) to screen the genome assemblies for any potential bacterial and fungal sequences. Contigs that were flagged with ‘EXCLUDE’ because they contained a fungal or bacterial hit were removed. Based on our findings we removed 42 contigs from the schMedS3h1, 12 contigs from the schMedS3h2, 5 contigs from the schPol2, and 7 contigs from the schLug1 assembly. Furthermore, we removed one contig from the schMedS3h1 assembly because it represented the mitochondrial genome.

### *De novo* repeat discovery and annotation

We annotated transposable elements using the Extensive de novo TE Annotator (EDTA) workflow (v2.1.0, ^92^). This approach augments the standard Repeat Modeler workflow with additional tools specifically targeted at LTR, Helitron, and TIR-Elements. We used parameters: ‘--species others --step all --sensitive 0 -anno 1’ and provided the previously manually curated repeat library generated for *S. mediterranea* ^29^ as a curated library. Additionally, Transposable element protein domains (Neumann et al., 2019) found in the assembled genomes were annotated using the DANTE tool available from the RepeatExplorer2 Galaxy portal (https://repeatexplorer-elixir.cerit-sc.cz/galaxy/) exploiting the REXdb database ^93^ (Viridiplantae_version_3.0).

To identify the overall repetitiveness of the genomes we performed *de novo* repeat discovery with RepeatExplorer2 ^94^. For *S. mediterranea* we used a repeat library obtained from the RepeatExplorer2 analysis of shotgun whole-genome Illumina paired-end sequencing (NCBI accession: SRR5408395). Since for *S. polychroa*, *S. nova*, and *S. lugubris* no Illumina data was available, we generated pseudo paired-end reads from 2Gb of CCS reads as input for RepeatExplorer2. All clusters representing at least 0.005% of the genomes were manually checked, and the automated annotation was corrected if needed. Contigs from the annotated clusters were used to build a repeat library. To minimize potential conflicts due to the occasional presence of contaminating sequences in the clusters, only contigs with average read depths ≥ 5 were included and all regions in these contigs that had read depths < 5 were masked. Genome assemblies were then annotated using custom RepeatMasker search with options ‘-xsmall -no_is -e ncbi -nolow’. Output from RepeatMasker was parsed using custom scripts (https://github.com/kavonrtep/repeat_annotation_pipeline) to remove overlapping and conflicting annotations.

Tandem repeat annotations were performed using TAREAN tool available from the RepeatExplorer2 output. Consensus monomers were then used as bait to annotate the presence and overall distribution of satellite DNA repeats in the assembled genome using the annotation tool available in Geneious R9 ^95^.

Since we noticed that a few highly repetitive regions were not annotated we additionally used Satellite Repeat Finder (srf, ^96^, commit: faf9c19) for annotation. We first generated a k-mer distribution of the genome assembly using kmc (^97^, v 3.2.1) and then used srf in combination with minimap2 (^98^, v2.24) to identify regions containing regions with high k-mer abundance. We then manually inspected all regions where srf resulted in an additional annotation and added them to the RepeatExplorer2 annotation.

### Long read Oxford Nanopore sequencing

Several adult animals of different statuses and sizes (i.e. starved for either 2 weeks and 1 month), regenerating fragments at several stages (from 0 to 7 days after cut) and isolated heads were collected in order to ensure the best transcriptomic diversity. Total RNA was extracted from snap-frozen planarian tissue using the protocol described in ^99^After the phenol-chlorophorm extraction step, RNA was purified using a Clean & Concentrator-25 kit (Zymo). Since read size distribution in Nanopore sequencing is usually biased towards the shorter transcripts, in order to partially counteract this effect we employed the manufacturer’s protocol variant optimised for the enrichment of transcripts longer than 200nts. RNA quality and quantity were assessed using Bioanalyzer RNA 6000 Nano Kit (Agilent). The poly-A+ fraction of RNA was isolated using Oligo d(T)_25_ Magnetic Beads (New England BioLabs Inc.) following the commercial protocol for Mammalian Cells provided by the manufacturer. Briefly, 14µg of total RNA were diluted into 250µL of Lysis/Binding buffer and used as input of the isolation procedure. 50µL of oligo-dT beads were employed for each round. After 2 rounds of isolation, the resulting poly-A+ RNA fraction (corresponding to 0.7-2% of the starting amount) was then purified again on a Clean & Concentrator-5 Column Kit (Zymo) and eluted in 10µL of molecular-grade water.

The direct RNA and cDNA libraries for Oxford Nanopore Sequencing were prepared using the SQK-RNA002, SQK-PCS109, andSQK-PCS111 kits, starting from 100ng and 4ng of poly-A+ RNA, respectively, following the manufacturer’s instructions. Sequencing was performed on the Oxford Nanopore Technologies (ONT) platform using a MinION and a PromethION P24 device. The prepared library was loaded onto a R9.4.1 flow cell, and sequencing was initiated following the manufacturer’s instructions. Real-time data acquisition was monitored using the ONT sequencing software MinKNOW.

### Genome annotation

The transcript annotation was generated by a hybrid genome-guided approach relying on both 1) dedicated long-read Nanopore cDNA/dRNA sequencing runs and 2) Illumina short-read and poly-adenylation (3P-seq) data obtained by publicly available datasets.

After read quality trimming, deduplication, filtering, and mapping (using HISAT2 ^100^and minimap2 ^98^ for short and long reads, respectively), a draft transcriptome was generated using Stringtie2 ^101^, then it was further refined using FLAIR ^102^ and a collection of custom scripts to filter high confidence isoforms. For details of the procedure and a step-by-step guide to the genome annotation analysis see Additional File 3. To designate a high-confidence gene set we applied additional filters using the repeat annotation, analysis of transcript expression across all nanopore data using the Nanocount program (Gleeson et al., 2021; 10.1093/nar/gkab1129), and a requirement for BLAST homology of at least 75% identity and 75% coverage against either the dd_smed_v6 or dd_smes_v1 transcriptomes, along with considerations for the open reading frame (ORF) length. We excluded all transcripts that overlapped more than 75% with a repeat annotation. For those transcripts with an ORF of 100 amino acids or more, we set a minimum expression threshold of 0.001 Transcripts per Million (TPM). In the case of transcripts with ORFs smaller than 100 but at least 50 amino acids in length, we included them in the high-confidence set under two conditions: if they had a BLAST hit and expression of at least 1 TPM, or in the absence of a BLAST hit, if their expression was at least 10 TPM.

### Benchmarking of *S. mediterranea* annotations

We compared our gene annotations with several transcriptomes that are commonly used in the field. Namely, dd_v1 a non-stranded *de novo* transcriptome assembly of the sexual strain of *S. mediterranea* (dd_Smes_v1, ^42^); dd_v6 a *de novo* transcriptome assembly of the asexual strain of *S. mediterranea* (dd_Smed_v6, ^42^); SMESG an *ab initio* gene prediction on basis of the previous dd_smes_g4 *S. mediterranea* genome assembly ^42^; Oxford_v1 a composite annotation of Neiro et al. ^39^ and García-Castro et al. ^45^ in combination with SMESG. We assessed the BUSCO content of the assemblies using BUSCO (v5.0.0) on the transcript level and merging the Complete and Duplicated category. We assessed how published transcripts were represented in the transcriptomes by aligning them to all transcripts available in NCBI using minimap2. To assess the mappability of these transcriptomes, we used sequencing data generated post-July 2020, which was after the creation of the benchmarked annotations. This approach was taken to avoid any bias that might arise from including reads used in the benchmarking process in the data creation. We utilized 13 sequencing libraries that varied in read lengths (152 – 302bp), sequencing platforms, and biological conditions (encompassing whole worms, dissociated cells, sorted X1 cells, and regenerating wound regions) to provide a comprehensive representation of common conditions in the field. The mapping efficiency was assessed using the BWA tool. Finally, we mapped all transcripts to the schMedS3h1 assembly using minimap2 and manually inspected 96 gene models for frame shifts, truncations, chimeras, and fragmentation.

### Nuclei isolation for ChIP-seq and HiC

For *S. mediterranea* 100 worms of ∼ 7 mm were treated with NAC for 10 min and afterwards rinsed with dH2O. The animals were transferred into a 15mL dounce tissue grinder and all excess water was removed. 10 mL modified cell buffer containing 1% formaldehyde (+PI + 10mM NaButyrate) was added and a timer set to 10 min. The tissue was homogenized using pestle A till resistance was minimal and incubated on a rocking shaker. Cross-linking was stopped by adding of glycine (125 mM glycine per 1% FA in 10 ml fixation buffer) and incubated for 5 minutes at RT. The sample was centrifuge in a swing bucket centrifuge for 10 min at 1000 g at 4°C. From this point on all steps were conducted at 4°C unless stated otherwise. The supernatant was gently removed and the pellet was resuspended in 10 mL of Buffer A (+PI + 10mM NaButyrate). Nuclei release was supported by mechanical disruption using pestle B till resistance was minimal. The sample was transferred into a 15mL tube and incubated for 15 min on ice on a rocking shaker. After pelleting at 1000 g for 10 min at 4°C, the supernatant was gently removed and the pellet was resuspended in 10 ml of Buffer B (+PI + 10mM NaButyrate) and incubated shaking vertically for 15 min on ice. The sample was then filtered through a 50μm mesh and an aliquot of 100μL was used for nuclei counting. The remaining solution was centrifuged at 1000 g for 15 min at 4°C, then the supernatant removed. The nuclei pellet was resuspend in 1 mL buffer B and the nuclei were distributed into 2.0*107 aliquots for ChIP-seq experiments or 1.1*106 nuclei aliquots for HiC. Finally, the samples were centrifuged at 1050 g for 10 min at 4°C, supernatant was removed and the pellets snap frozen and stored at -80°C.

### SDS-PAGE and Western Blot

Pulldown samples were directly mixed with 6x Laemmli buffer and decrosslinked and denatured for 15 min at 95°C. The unbound fraction was concentrated by acetone precipitation. Four volumes of cold (-20°C) acetone were added to the sample, mixed, and incubated for 60 min at -20°C. Then the sample was centrifuged for 10 min at 15,000 × g and the supernatant was removed. The protein pellet was finally resuspended in the same volume as the pulldown sample, mixed with 6x Laemmli buffer and decrosslinked and denatured at 95°C for 15 min. Samples were run on NuPAGE Novex 4–12% Bis-Tris protein gels in 1× MOPS running buffer, transferred onto Nitrocellulose membranes in transfer buffer (1× MOPS with 20% MeOH), blocked in 1× PBS with 0.1% Tween20 and 5% w/v nonfat dry milk and incubated with primary antibody (H3K4me3 milipore 07-473 Lot#3381394) diluted 1:5.000 in 1× PBS with 0.1% Tween20 and 5% nonfat dry milk. Membranes were washed with washing buffer (1× PBS with 0.1% Tween20) prior to incubation with fluorescent secondary antibodies (anti-Rabbit IRDye 800CW, LICOR) diluted 1:20.000 in blocking solution. Membranes were washed with washing buffer, followed by a final wash step in 1× PB without Tween20. Stained membranes were dried and imaged on a LI-COR Odyssey imager.

### Chromatin immunoprecipitation and sequencing (ChIP-seq)

The frozen nuclei pellet was resuspended in 1mL of lysis buffer +PI +0.5mM NaButyrate and incubated for 25 min at 4°C while rocking. Then the sample was transferred into a miliTUBE with AFA fiber (Covaris Part# 520135) and topped up with lysis buffer. The sample was sonicated using the Covaris sonicator S220 at 140 Peak Power, 5,0 Duty Factor and 200 Cycles/Burst for 15 min at max 8°C, then transferred into a fresh 1,5mL tube and centrifuged for 5 min at 4°C (10 000g) to pellet debris. The supernatant was split into aliquots of 150µL and subsequently used for pulldown experiments. For input one aliquot was topped up with Buffer B to 475 µL and 20 μL 5M NaCl and 5 μL proteinase K (20 mg/mL) was added and incubated at 65°C overnight to reverse cross-links. The next day the sample was removed from the thermoblock and cooled down to RT. Subsequently 2 μl RNase (10mg/mL) was added and incubated for 1h at 37°C. DNA was isolated using the PCI-DNA extraction method. Therefore, the sample was transferred into a phase lock tube, 500 μL PCI-mixture was added to the sample, vortexed and centrifuged at full speed for 5 min at RT. 500μL of pure Chloroform was added, vortexed and centrifuge at full speed for another 5 min at RT. The upper aqueous phase was transferred into a 1,5mL tube, 500 μl isopropanol (-20°C), 100 μl 3M sodium acetate (NaOAc) and 2 μl glycogen were added and then incubated for 60 min at -20°C. After incubation the sample was centrifuged at full speed for 15 min at 4°C, before the supernatant was removed. The pellet was washed with 500 μl 96% ethanol (-20°C) and centrifuged at full speed for 10 min at 4°C. A second washing step was performed with 500 μl 70% ethanol (-20°C) and then centrifuged at full speed for 10 min at 4°C. After removing the supernatant the pellet was dried at 55°C. The DNA was resuspended into 50 μl EB buffer and stored at -20°C before library preparation.

For pull-down one aliquot was used per target. 2 Volumes of Dilution Buffer were added to the sample and topped up to 1mL with RIPA buffer (+PI +0.5mM NaButyrate). 2,5µ α-H3K4me3 (milipore 07-473 Lot#3381394) or 5µL α-H3K27ac (active motif #39133 Lot#16119013) was added and incubated for 1h at 4°C with gentle agitation (6rpm). Subsequently 30µL Magna ChIP Protein A Magentic Beads were added to the sample and incubate at 4°C with gentle agitation (6rpm) overnight. Beads were accumulate using a magnetic rack. Washing was performed by adding 1 mL washing solution, incubating with gentle agitation for 5min at 4°C and removing solution. The following buffers were used for washing chronologically RIPA, HiSalt, LiCl, TE (2 times). After the final wash, the TE buffer was thoroughly removed and the bead-bound complexes released by incubating in 100µL elution buffer at 4°C for 30min. The supernatant was transferred into a fresh tube and continued with reversing cross-links and DNA isolation, as stated above.

Immuno-precipitated DNA samples at an input amount of 2-100 ng were subjected to Illumina fragment library preparation using the NEBnext Ultra II DNA library preparation chemistry (New England Biolabs, E7370L). In brief, DNA fragments were end-repaired, A-tailed and ligated to unique-dual indexed Illumina Truseq adapters. Resulting libraries were PCR-amplified for 15 cycles using universal primers (Primer 1: CAAGCAGAAGACGGCATACGAGAT and Primer 2: AATGATACGGCGACCACCGA*G; *: Phosphothioate bond), purified using XP beads (Beckman Coulter) with a bead to library ratio of 1:1. The libraries were size selected using XP beads with a 0.6:1 right and 1:1 left bead to library ratio and if needed subjected to an extra 0.8:1 bead to library ratio to remove the left over adaptor dimers. They were checked for their quality and quantified using Fragment Analyzer (Agilent). Final libraries were subjected to 100-bp-paired-end sequencing on the Illumina NovaSeq6000 and 75-bp-paired-end and single-end on the NextSeq500 platform to a depth of 30-70 million fragments per library.

### Chromatin conformation capture (HiC)

Chromatin conformation capturing was done using the ARIMA-HiC High Coverage Kit (Article Nr. A101030-ARI) following the Arima documents (User Guide for Animal Tissues, Part Number A160162 v00). 1×10^6^ nuclei from each species was crosslinked and went into lysis step. The crosslinked gDNA was digested with a cocktail of four restriction enzymes. The 5’-overhangs were filled in and labelled with biotin. Spatially proximal digested DNA ends were ligated, and the ligated biotin containing fragments were enriched and went for Illumina library preparation, following the ARIMA user guide for Library preparation using the Kapa Hyper Prep kit (ARIMA Document Part Number A160139 v00). The barcoded HiC libraries ran on an S4 flow cell of an Illumina NovaSeq 6000 with 2×150 cycles.

### Assay for transposase-accessible chromatin with sequencing (ATAC-seq)

The ATAC-seq experimental protocol is a modification of the protocol from ^46^. For each biological replicate of the wt and X-ray analysis 10 worms of ∼ 7 mm were treated with NAC for 10 min and afterwards rinsed with dH2O. The animals were transferred into Covaris tissueTUBEs (TT05M TX), snap frozen in liquid nitrogen and crushed with setting 2 using the Covaris CryoPrep (CP02) device. Snap freezing and crushing was repeated a second time, before the powder was stored at -80°C. For nuclei isolation two corresponding samples were processed simultaneously (wt and X-ray). The sample was resuspended in 1,5 mL of cold 1× unstable homogenization buffer and transferred into a pre-chilled tissue grinder. The powder was further homogenized with a dounce homogenizer and pestle B (clearance 0.0005-0.0025 in.) on ice with 10 strokes and afterwards filtered through a 50μm mesh. 1,5 mL of 50% Iodixanol Solution (with freshly added Spermidine and Spermine) was added and well mixed.

For gradient centrifugation 2000 μl of a 40% Iodixanol solution was transferred into a 15mL tube. 1000 μl of 30% Iodixanol solution was slowly layered on top of the 40% mixture. Finally, the 25% Iodixanol-nuclei solution was carefully added on top of the 30% mixture. Separation was performed by centrifuge at 3,000 RCF with brake off at 4 °C for 20 min. Then the nuclei band was collected and transferred to a fresh tube. 2 volumes of ATAC-RSB-Buffer + 1mM DTT were added before nuclei were counted. For each biological replicate, 3 libraries were prepared for tagmentation. Therefore, 5*104 nuclei were transferred in each tube and centrifuged at 900g for 7 min at 4 °C. The supernatant was discarded, and the pellet was resuspended in 25 μL ATAC-seq Reaction Mix +1mM DTT, then 25μL commercial buffer from Illumina Tagment DNA TDE1 Enzyme and Buffer kit was added and the mixture was incubated for 30 min at 37°C 1000RPM. Tagmentation was stopped by proceeding with the Zymo PCR Clean and Concentrate kit. Finally, tagmented DNA was eluted in 10 μL and used for library amplification.

A total of 10 µl of purified tagmented DNA was indexed and pre-amplified for initial 5 PCR cycles with 1× KAPA HiFi HotStart Readymix and 100 nM unique dual index P5 and P7 primers compatible with Illumina Nextera DNA barcoding, under the following PCR conditions: 72°C for 5 min, 98°C for 30 s, thermocycling for 5 cycles at 98°C for 10 s, 63°C for 30 s and 72°C for 1 min. Subsequently, a qPCR on the LightCycler 480 (Roche) was performed with 1 µl of the pre-amplified material to determine the remaining PCR cycle numbers (7-13) to avoid saturation and potential biases in library amplification (see Buenrostro et al. 2015). Purification and double-sided size selection of amplified libraries was done with AMPure XP beads (Beckmann Coulter; starting with a 1.55x volume of XP bead purification, followed by a 0.6x/1.55x double-sided size selection, an additional 0.6x/1.55x double sided size selection was performed if needed), and checked for their quality and quantity on a Fragment Analyzer (Agilent). Libraries were sequenced on the Illumina NovaSeq 6000 with PE 100 bp reads to a depth of 40-150 M read pairs.

### ATAC-seq and ChIP-seq read processing and mapping

ATAC-seq and ChIP-seq reads were trimmed using trim_galore (0.6.6) in --paired mode and QC was done using fastqc (0.11.9) prior to mapping. Genomes were indexed using bwa index. Due to the size limitations for BAI indexing we split Chromosome 1 at position 414,900,000 and 166,500,000 of *S. lugubris* and *S.nova*, respectively. Of note, in the proximity of these positions no gene annotations were observed. Trimmed reads were mapped to the corresponding genome using bwa (0.0.17) with the following command “bwa mem -M”. Mapped reads were filtered using samtools fixmate -m and sort, before PCR duplicates were removed using picard (2.25.5) with the following setting MarkDuplicates REMOVE_DUPLICATES=True VALIDATION_STRINGENCY=LENIENT. Finally reads were filtered using samtools view samtools view -h -b -q 20 -f 3. Libraries sequenced on the different sequencing Chips, were merged after this step using samtools merge and subsequently sorted an index as previously described.

### ChIP-seq analysis

Peak calling for ChIP-seq data was performed using MACS2 ^103^ (2.2.7.1) running macs2 callpeak -t *SAMPLE* -c *INPUT* -f BAMPE --nomodel --bdg --keep-dup all -g 6.44e8. Effective genome size was calculated using the number of uniquely mappable bases using a k-mer based approach with the khmer software ^104^ and a k-mer size of 32 and a read length of 100bp. Peak annotations were generated with the ChIPseeker library ^105^ (1.33.2) utilizing the annotatePeak function, defining Promoter as 3000 – 0 upstream of the TSS. Profile plots and heatmaps were generated using deeptools ^106^ (3.5.1) package using different tools and modified manually for better readability. Signal tracks were generated using SparK.py (v2.6.2, ^107^). Intersections of different regions were done using bedtools (2.30.4). To quality control the ChIP-seq data we assigned the associated genes into quartiles of gene expression in a dataset of 7mm sized wild type asexual *S. mediterranea*. To quantify expression we used RNA-seq reads, quality trimmed using trimmomatic, mapped and quantified using STAR and normalized to transcripts per million per sample. We then used the mean of both samples as the expression estimate of that gene.

### ATAC-seq analysis

ATAC-seq QC was performed using the ATAC-seq QC library (1.16.0 and 1.22.0, ^108^). Fragment size distribution was calculated and visualized with the fragSizeDist function. For the distribution of fragments corresponding to nucleosomal-free and mononulceosomal regions, the enrichedFragments function was used. Technical replicates were merged using samtools merge prior to further processing. To eliminate biases in peak numbers between contrasting samples (wt vs X-ray and head vs tail) due to differences in sequencing depths samples have been subsampled to the lowest library size before peak calling. Peak calling was performed with MACS2 using the following parameters: -f BAMPE --keep-dup all and species-specific effective genome size with the -g parameter. Peak sets of the same biological condition (e.g. head region) were combined using ChIP-R ^109^ (1.23.5) with the parameter -- minentries 3. Summit information was added to the generated files by averaging the positions of the 3 original summits using a custom script. Final consensus peak sets for each species were generated using bedtools intersect by adding condition-specific peaks to the wild-type peak set. Signal tracks were generated using SparK.py.

### Evolutionary conservation of ATAC-seq peaks

Existing methodologies for regulatory region conservation assessment (e.g. ^110^), were observed to inadequately deal with the high extent of fragmentation and duplication in planarian genomes. Additionally, they do not leverage ATAC-seq data in the recipient species to filter duplicated peak regions. Hence, a custom protocol was developed using a series of scripts to address the inadequacies associated with peak fragmentation and duplication. The base dataset was constituted by the set of consensus ATAC-seq peaks (see ATAC-seq data analysis). The *S. mediterranea* pseudohaplotype 1 served as the frame of reference in the subsequent steps. The peak region (RGN) and a 51bp extension of the summit (SUM) for each species were processed using halLiftover ^111^. This allowed for an independent assessment of the presence of the summit in the recipient species. Any RGN liftover regions less than 20bp in size were disregarded, and fragmented RGN liftover regions separated by less than 100bp were merged. RGN liftover regions that overlapped with the corresponding SUM liftover were identified as ‘conserved regions’. Those RGN liftover regions without a SUM overlap were classified as ‘not conserved’. The conserved regions were characterized by a median size of 555bp, in contrast to the short, highly fragmented size of the non-conserved liftovers (Additional File 1: Section 3), indicating the efficacy of the developed pipeline in filtering out low-confidence liftovers. Conserved regions were then examined for an overlap with the ATAC-seq signal in the recipient species. A liberal scoring criterion was employed, treating any overlap as a hit and marking conserved regions with an overlap as a ‘conserved peak’. Finally, the results of the conservation assessment in each species were used to annotate each *S. mediterranea* pseudohaplotype 1 peak. The highest conservation status was given to peaks that showed conservation in all three recipient species.

### Synteny analysis

Since we expected gene order to be largely preserved across Schmidtea, we used the R package GENESPACE (v1.0.8, ^63^) to infer gene-order based syntenic blocks. GENESPACE uses a combination of Orthofinder and a reimplementation of the MCScanX algorithm ^112^. We then visualized the resulting syntenic blocks using the built-in riparian plot function.

To understand the conservation of synteny across increasingly larger phylogenetic distances, we employed three tools and four approaches. First, we annotated BUSCO genes present in Schmidtea and the parasites using BUSCO (v5.0.0, ^113^)and then identified those genes that were present in each species as a single copy gene. We then visualized the location of the BUSCOs in each assembly by coloring them according to the chromosomal location in *Schistosoma mansoni*. Second, we used Orthofinder (v2.5.5, ^114^) with default parameters and with protein sequences representing the longest isoform of the entire protein coding fraction of the *Schmidtea* and the parasites annotations as input. We then processed the resulting orthogroups for each pairwise combination always only retaining pairwise single copy orthologs. Thus, for each pair the number of orthologs used differed. We visualized the distribution of orthologs using a dotplot and conducted chi-squared tests against the null hypothesis that orthologs are randomly distributed across the chromosomes. A significant p-value indicates that there is a clustering of orthologs based on the chromosome combination (i.e. preservation of synteny). Given potential violations of chi-square test assumptions by our genomic data, we not only conducted a conventional significance test based on the chi-square distribution but also employed a permutation test with 100,000 permutations to assess significance. We calculated the effect size ‘Cramer’s V’ to determine the amount of clustering. Cramer’s V is 0 for a random distribution and 1 for a perfect correlation between chromosomes. Third, we used the reciprocal-best-blast and Fisher’s exact tests approach as implemented in the ODP tool ^10^ (v0.3.0) to infer pairwise orthologs between species and test for synteny conservation. Given these analyses test for a direct association between two chromosomes/scaffolds, we included the highly contiguous, albeit not chromosome-scale, genome assembly of *Macrostomum hystrix*. Finally, we assessed if ancestral metazoan linkage groups (MALG), i.e. linkage groups that are present in bilaterians, cnidarians, and sponges, defined in ^8^ were preserved in the flatworm genomes using the ODP tool. The tool uses hidden-markov models to identify homologs to the MALG proteins and tests for their enrichment on particular chromosomes using Fisher’s exact test. We then summarized the results using the chromosome tectonics algebra defined in ^8^.

### Synteny breakpoint enrichment

We investigated whether synteny breakpoints were disproportionately associated with specific repetitive sequences using the R package GenomicRanges (v.1.46.1). For each syntenic block, delineated by GENESPACE, across *Schmidtea* species, we established 10kb flanking windows in both genomes involved in the pairwise combinations. Subsequently, we evaluated their enrichment by contrasting them against 1,000 iterations of random placements of an equivalent number of windows. We performed a two-tailed test to assess if the observed elements were present in higher or lower quantities than expected from the random iterations. To account for multiple testing, we adjusted the p-value for all tested elements with at least 10 member on average (this threshold was used to prevent loss of statistical power when testing elements that were exceedingly rare) utilizing the False Discovery Rate method.

## Data Access

The code repositories and sequence data associated with this paper are currently under preparation for release. These resources will be made publicly available following the peer-review process of our work.

The Neodermata genomes and annotations are publicly available at https://parasite.wormbase.orghttps://parasite.wormbase.org under the following accessions: *Clonorchis sinensis*: PRJNA386618; *Schistosoma mansoni*: PRJEA36577; *Taenia multiceps*: PRJNA307624; *Hymenolepis microstoma*: PRJEB124. The *Macrostomum hystrix* genome is available from the ENA database under the following accession: GCA_950097015. the *Macrostomum hystrix* gene annotations are available from the Zenodo repository under the following accession: https://doi.org/10.5281/zenodo.7861770.

## Acknowledgements

We thank Heino Andreas, Rick Kluivert, Jens Krull, and the MPI-NAT animal services staff for worm care and the maintenance of the species collection. We thank Tobias Boothe for picture acquisition. We thank Juliana G. Roscito for sequencing support and close collaboration during method development. We thank Hanh Vu for access to *S. mediterranea* RNA-seq data. We thank the following facilities for their support: the DRESDEN Concept Genome Center, part of the MPI-CBG and the technology platform of the CMCB at the TU Dresden, supported by DFG (INST 269/768-1); the Genomics Core Facility of the European Molecular Biology Laboratory in Heidelberg; and the IIT Genomics Facility. We are grateful to Diego Vozzi, Yeraldin Castillo Spelorzi, and Edoardo Henzen for their invaluable support to the Nanopore sequencing experiments. JCR received funding from the European Research Council (ERC) under the European Union’s Horizon 2020 research and innovation program (grant agreement number 649024), from the German Research Foundation (project RI 2449/51), from the Behrens-Weise Foundation, and from the Max Planck Society. JNB was supported by Swiss National Science Foundation Grant P500PB_206673. LP, AC and SG were supported by intramural funding of the Istituto Italiano di Tecnologia. AP was supported by the SPP2202 Priority program (Project No. 422389065).

