## Additional File 1 for "A comparative analysis of planarian genomes reveals regulatory conservation in the face of rapid structural divergence"

#### Table of Contents

*1 Genomic resources* ... *2*

1.1 Assembly quality assessment ... 2

1.2 Comparison to schMedS2 ... 3

1.3 Phasing efficiency ... 4

1.4 Gene annotation benchmarking ... 5

1.5 Chimeric gene annotations ... 6

*2 Regulatory element annotation* ... *7*

2.1 ATAC-seq quality control ... 7

2.2 ChIP-seq quality control ... 8

2.3 ATAC-seq peak classification ... 9

*3 Regulatory region conservation* ... *10*

*4 Synteny* ... *14*

4.1 GENESPACE based synteny ... 14

4.2 Synteny breakpoint inspection ... 15

4.3 Orthofinder based synteny ... 16

4.4 ODP based synteny ... 19

4.5 MALG conservation ... 27

*5 References* ... *30*

### 1 Genomic resources

#### 1.1 Assembly quality assessment

Table 1 Summary statistics for genome assemblies of the *S. mediterranea* sexual strain. dd_Smes_g4: Our previous diploid consensus assembly [1]; schMedS2: chromosome-scale scaffolding of most contigs of the dd_Smes_g4 assembly[2], schMedS3h1 and schMedS3h2: haplotype phased assemblies (this study).

| | **dd_Smes_g4** | **schMedS2** | **schMedS3h1** | **schMedS3h2** |
|---|---|---|---|---|
| # contigs | 481 | 4 | 662 | 432 |
| Total length (bp) | 773,939,492 | 763,446,644 | 840,173,815 | 819,865,861 |
| GC (%) | 29.63 | 29.6 | 29.59 | 29.59 |
| N50 (bp) | 3,854,845 | 265,042,666 | 270,168,396 | 268,961,546 |
| chr-scaffold (%) | - | 99 | 95 | 96 |
| unplaced (Mb) | - | 10.5 | 42 | 32.8 |

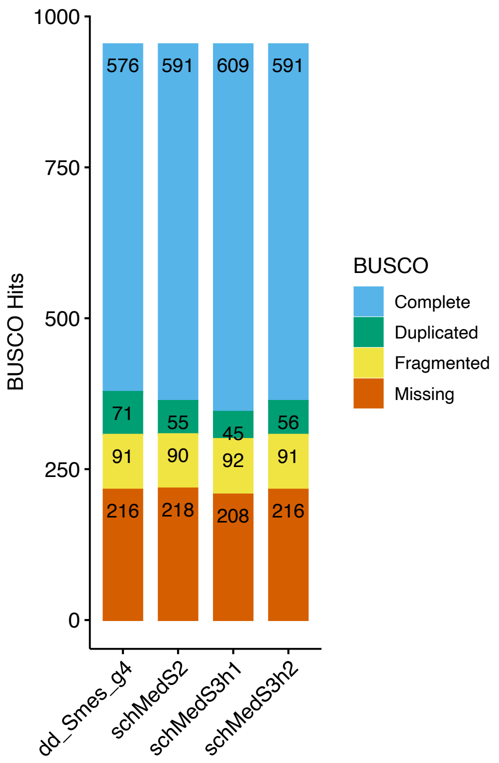


Figure 1 BUSCO score of our previous assembly dd_Smes_g4, the schMedS2 assembly, and the new phased assembly, showing increased completeness in the new assembly.

#### 1.2 Comparison to schMedS2

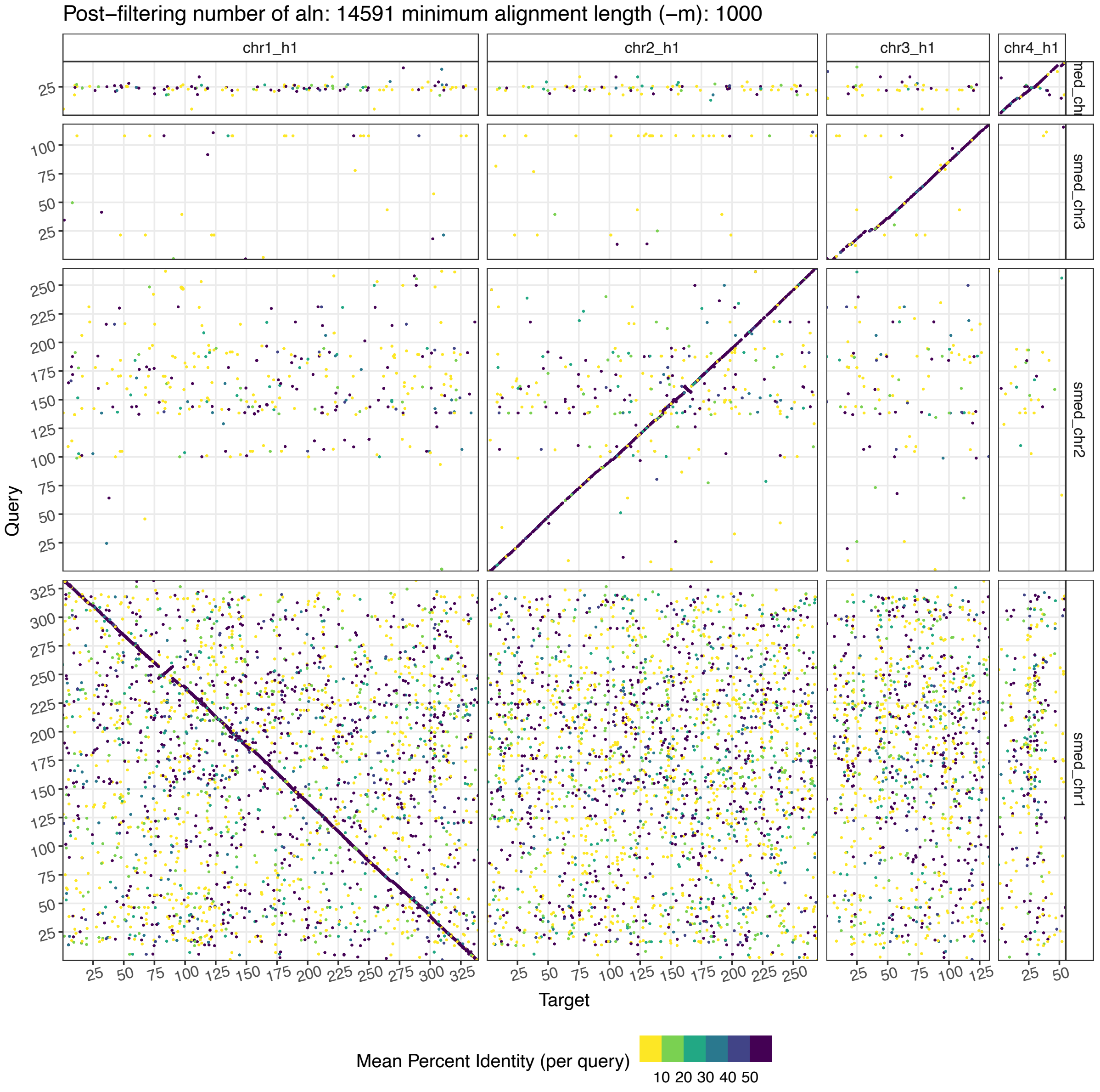


Figure 2 Dotplot alignment between schMedS3h1 from this study and schMedS2 from [2]. Dots represent alignments of at least 1000bp and the color indicates alignment quality.

### 1.3 Phasing efficiency

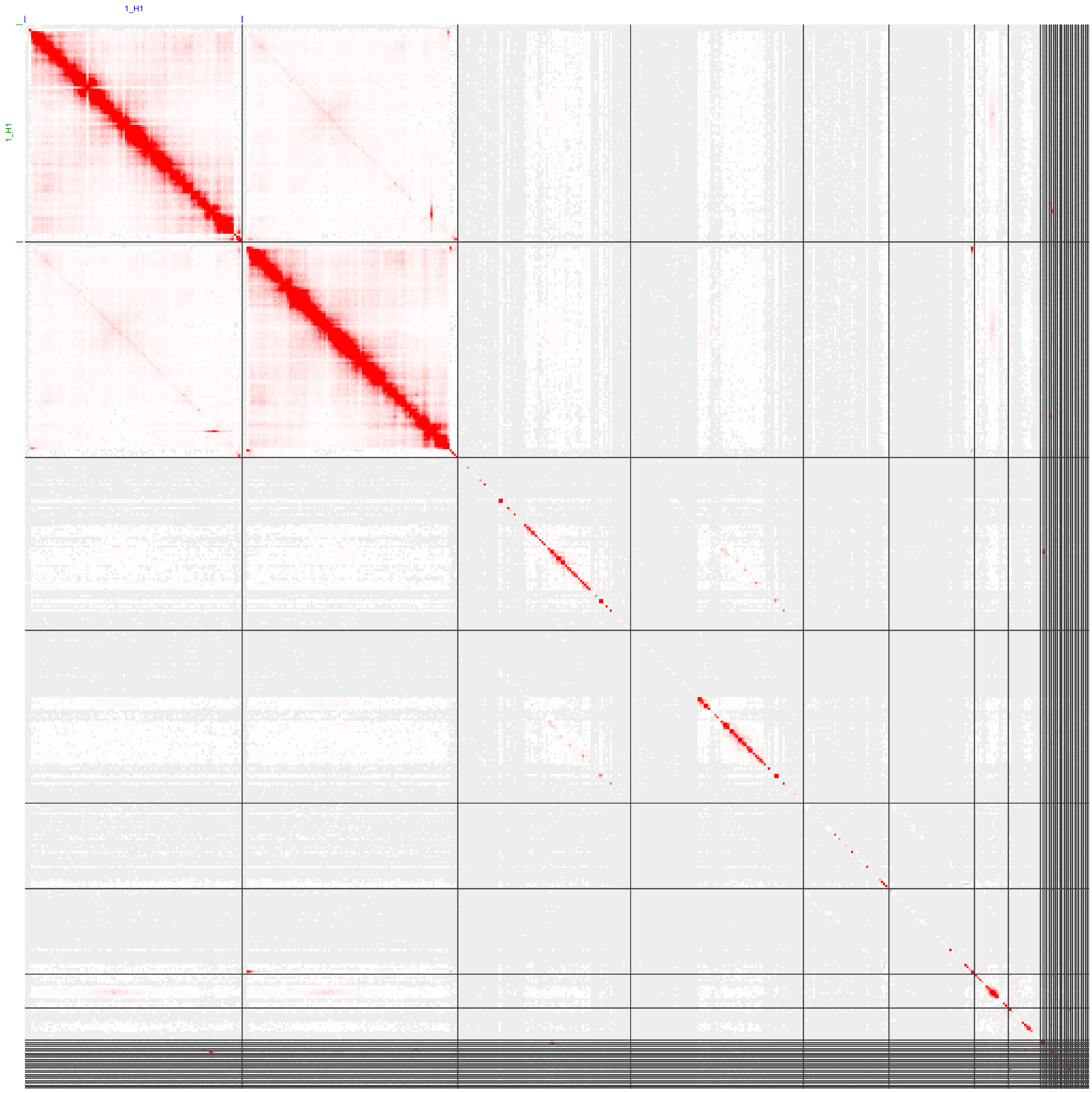


Figure 3 HiC map of the diploid assembly schMedS3BH showing a high number of uniquely mapping (red cells) on Chromosome 1, a large proportion of Chromosome 2 and Chromosome 4, and only few regions of Chromosome 3.

### 1.4 Gene annotation benchmarking

Table 2 Summary of the 16 genes deposited in NCBI that were not detected in the S3BH annotation. Two genes could not be mapped, likely due to assembly gaps. The other 14 mapped uniquely to genome and eight of them were present in some of our previous annotations (SMEST, dd_Smed_v6, or dd_Smes_v1), indicating errors in the S3BH gene preditions.

| Missing gene | Annotation | Map location | Predicted in x other annotations: | Category |
|---|---|---|---|---|
| KT163545.1 | Smed slc15a-9 (slc15a-9) | no hit | n.a. | Assembly gap |
| KX018976.1 | Smed NPYR-16 (npyr-16) | no hit | n.a. | Assembly gap |
| BK007012.1 | TPA_inf: Smed cerebral peptide prohormone like-1 | unique | 2 | missing prediction |
| BK007023.1 | TPA_inf: Smed secreted peptide prohormone-12 | unique | 1 | missing prediction |
| FJ471488.1 | Smed noggin-like protein 6 (nlg6) | unique | 3 | missing prediction |
| FJ588606.1 | Smed cyclinB-like protein | unique | 0 | missing prediction |
| KT163558.1 | Smed slc16a-13 (slc16a-13) | unique | 0 | missing prediction |
| KT163560.1 | Smed slc16a-15 (slc16a-15) | unique | 0 | missing prediction |
| KT163565.1 | Smed slc16a-20 (slc16a-20) | unique | 0 | missing prediction |
| KT163605.1 | Smed slc22a-7 (slc22a-7) | unique | 3 | missing prediction |
| KT163651.1 | Smed slc25a-33 (slc25a-33) | unique | 0 | missing prediction |
| KT163655.1 | Smed slc26a-1 (slc26a-1) | unique | 3 | missing prediction |
| KT163750.1 | Smed slc47a-3 (slc47a-3) | unique | 0 | missing prediction |
| KX018920.1 | Smed GCR123 (gcr123) | unique | 2 | missing prediction |
| KX018937.1 | Smed GCR141 (gcr141) | unique | 1 | missing prediction |
| KX018982.1 | Smed NPYR-8 (npyr-8) | unique | 2 | missing prediction |

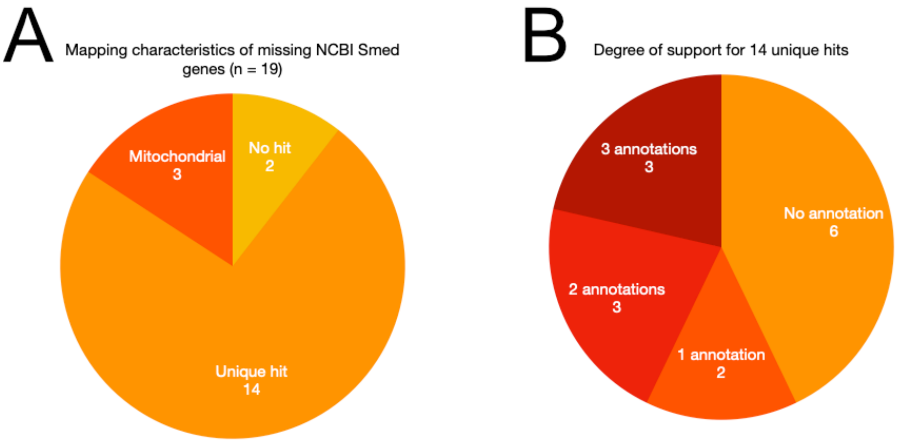


Figure 4 Details on the 19 transcripts deposited in NCBI that were not present in the S3BH annotation. **A** Mapping characteristic of the missing transcripts. **B** Details on whether the 14 uniquely mapped transcripts were present in some of the previous annotations (SMEST, dd_Smed_v6, or dd_Smes_v1).

### 1.5 Chimeric gene annotations

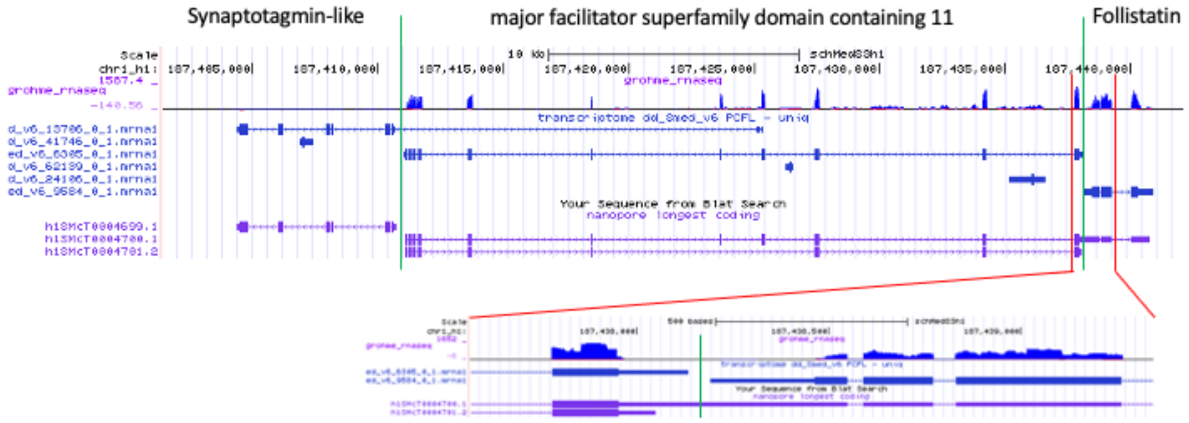


Figure 5 Genome browser view showing the chimeric merger of the gene annotations of Frz receptor and the Activin inhibitor follistatin. Note, the close proximity, same strandedness and similar RNA-seq coverage of both transcripts, making this a particularly challenging annotation problem.

### 2 Regulatory element annotation

#### 2.1 ATAC-seq quality control

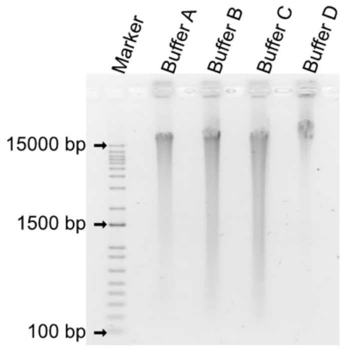


Figure 6 Impact of buffer composition on DNA integrity using native samples. Buffer A (10mM Tris- HCl pH7,5, 10mM NaCl, 3mM MgCl2, 0,1% Igepal), buffer B (10mM Hepes- NaOH pH7,9, 1,5mM MgCl, 10 mM KCl, 1mM EDTA, 0,05% Igepal, 0,5 mM DTT (added right before use)), buffer C: (10mM Hepes- NaOH pH7,9, 60 mM KCl, 1mM EDTA, 0,05% Igepal, 0,5 mM DTT (added right before use)) and buffer D (10mM Tris- HCl pH7,5, 10mM NaCl, 3mM MgCl2, 0,1% Igepal, 0,5mM Spermine, 0,25mM Spermidine, 0,5 mM DTT (added right before use)). Using buffer D resulted in sufficient isolation of high molecular weight DNA.

#### 2.2 ChIP-seq quality control

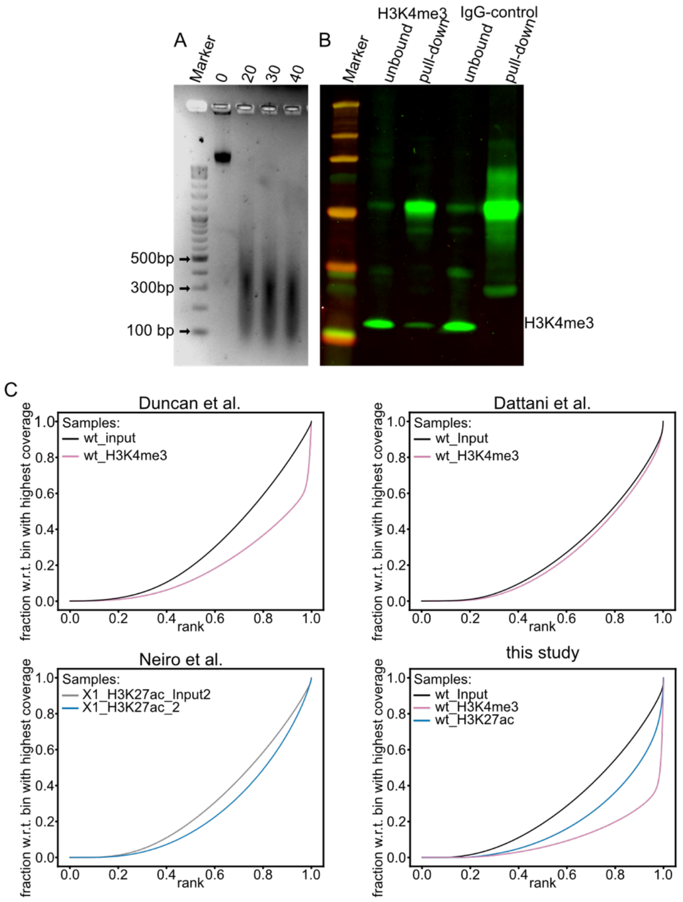


Figure 7 A: Identification of optimal sonication settings for chromatin fragmentation using the Covaris S2 sonicator with AFA tubes. (5%Duty Cycle, Intensity 4, 200 Cycles/Burst, 30 sec/cycle). Displayed are DNA samples from crosslinked nuclei without sonication (0), 20 cycles, 30 cycles and 40 cycles. B: Western blot validation of H3K4me3 ChIP-seq pulldown. H3K4me3 pulldown enriched for protein of interest (left), while IgG control did not show any H3K4me3 enrichment (right), confirming sufficient purification. C Fingerprint plot displaying ChIP enrichment in different experiments. A diagonal line indicates uniformly distributed reads along the genome (no enrichment), while a steep curve indicates strong enrichment.

#### 2.3 ATAC-seq peak classification

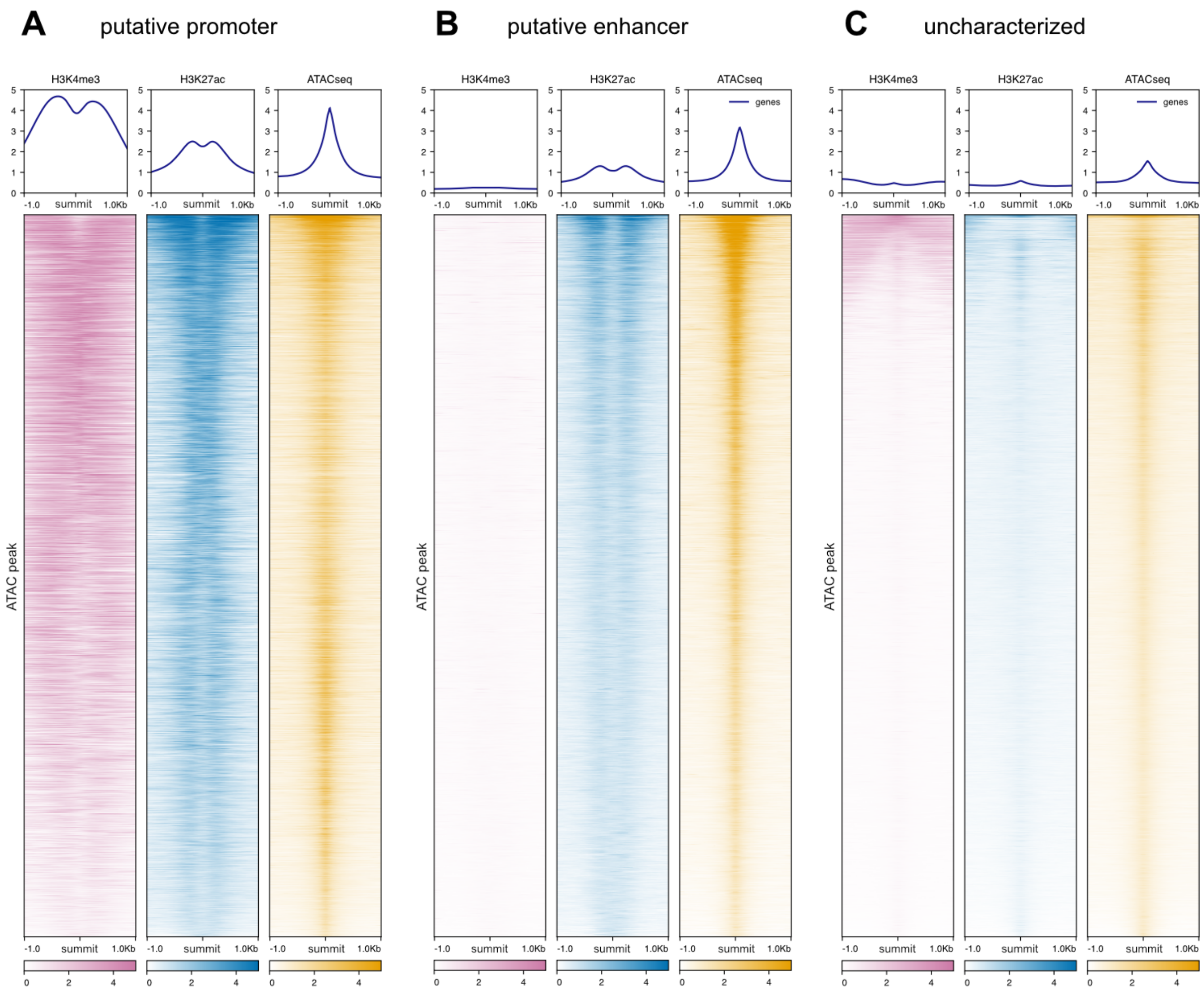


Figure 8 Heatmap of ChIP signal at the summits of ATACSeq peaks. A: H3K4me3 and H3K27ac signal at the summits of putative promoters. B: H3K4me3 and H3K27ac signal at the summits of putative enhancers. C: H3K4me3 and H3K27ac signal at the summits of the remaining uncharacterized ATACSeq peaks.

Table 3 Genomic location of ATAC-seq peaks depending on their classification. Annotation is based on the closest gene. Application of a Chi-square test showed that annotation was not equally distributed across the categories. We used Z-score-based post-hoc tests with a Bonferroni correction for multiple testing to determine the enrichment of element types.

| | **putative promoter** | | | | **putative enhancer** | | | | **uncharacterized** | | | |
|---|---|---|---|---|---|---|---|---|---|---|---|---|
| | N | % | Z | p | N | % | Z | p | N | % | Z | p |
| <1kb TSS | 8444 | 61.4 | 150.8 | < 0.001 | 454 | 4.3 | -41.6 | <0.001 | 1263 | 4.1 | -98.1 | <0.001 |
| 1-2kb TSS | 48 | 0.3 | -11.7 | < 0.001 | 101 | 0.9 | -4 | 0.0019 | 599 | 1.9 | 13.3 | <0.001 |
| 2-3kb TSS | 39 | 0.3 | -9.8 | < 0.001 | 72 | 0.7 | -3.8 | 0.0037 | 448 | 1.4 | 11.5 | <0.001 |
| 5UTR | 877 | 6.4 | 31.5 | < 0.001 | 147 | 1.4 | -9 | <0.001 | 445 | 1.4 | -20.2 | <0.001 |
| Exon | 2044 | 14.9 | -9.6 | < 0.001 | 2402 | 22.6 | 15.1 | <0.001 | 5312 | 17 | -3.6 | 0.0066 |
| Intron | 1066 | 7.7 | -62.9 | < 0.001 | 5240 | 49.2 | 51.7 | <0.001 | 9705 | 31.1 | 13.7 | <0.001 |
| Downstream | 38 | 0.3 | -10 | < 0.001 | 99 | 0.9 | -1.1 | 1.0000 | 432 | 1.4 | 9.6 | <0.001 |
| Distal intergenic | 1203 | 8.7 | -61.2 | < 0.001 | 2130 | 20.0 | -23.5 | <0.001 | 12977 | 41.6 | 71.8 | <0.001 |

### 3 Regulatory region conservation

Table 4 Conservation of ATACSeq peaks depending on their characterization. Application of a Chi-square test showed that conservation was not equally distributed across the categories (X-squared = 9745.1, df = 4, p-value < 2.2e-16). We used Z-score-based post-hoc tests with a Bonferroni correction for multiple testing to determine the enrichment of element types. All comparisons were highly statistically significant (all p-values <0.001), indicating enrichment for conserved peaks in putative promoters and putative enhancers (positive Z-scores) but enrichment for no conservation in uncharacterized peaks.

| | **putative promoter** | | **putative enhancer** | | **uncharacterized** | |
|---|---|---|---|---|---|---|
| | N | Z-score | N | Z-score | N | Z-score |
| highly conserved | 4261 | 68.4 | 2343 | 28.1 | 963 | -81.8 |
| partially conserved | 6659 | 22.7 | 6006 | 38 | 9661 | -49.9 |
| not conserved | 2839 | -69.4 | 2296 | -56.7 | 20557 | 105.3 |
| % highly conserved | 31 | | 22 | | 3.1 | |
| % min. partially conserved | 79.4 | | 78.4 | | 34.1 | |

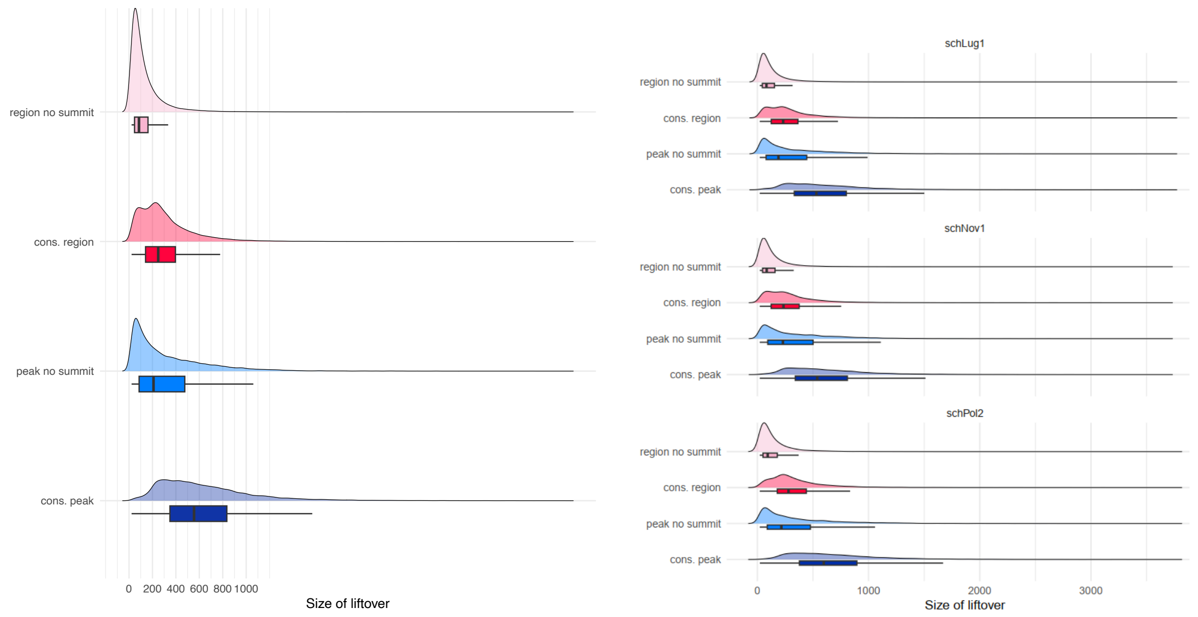


Figure 9 Size of *S. mediterranea* ATAC-seq peak liftover depending on the conservation classification. Left side: Summary across all three liftover species showing liftovers that were categorized as not conserved (region no summit and peak no summit), were short compared to conserved peaks. Right side: Same plot but split by species.

Table 5 Size distribution of schMedS3h1 liftover by category.

| | median size | SD |
|---|---|---|
| region no summit | 85 | 141.4 |
| cons. region | 249 | 246.3 |
| peak no summit | 209 | 329.0 |
| cons. peak | 555 | 410.2 |

Table 6 Size distribution of schMedS3h1 liftover by category and species.

| species | type | median size | SD |
|---|---|---|---|
| *S. lugubris* | region no summit | 82 | 133.3 |
| | cons. region | 232 | 228.2 |
| | peak no summit | 190 | 315.8 |
| | cons. peak | 531 | 394.0 |
| *S. nova* | region no summit | 84 | 136.3 |
| | cons. region | 235 | 234.4 |
| | peak no summit | 230 | 328.6 |
| | cons. peak | 539 | 393.4 |
| *S. polychroa* | region no summit | 93 | 158.7 |
| | cons. region | 280 | 268.2 |
| | peak no summit | 216 | 346.5 |
| | cons. peak | 597 | 436.8 |

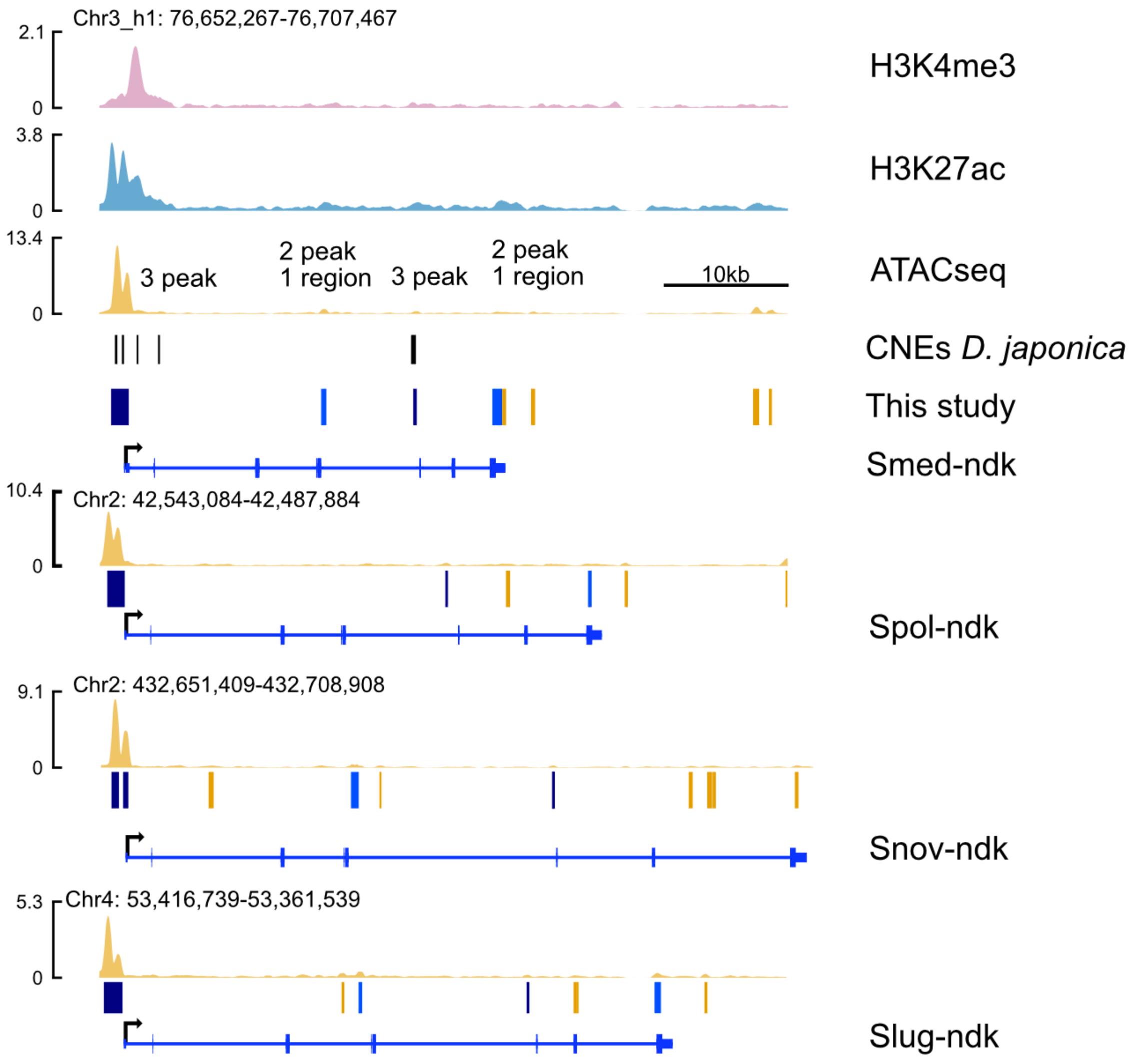


Figure 10 Example of a highly conserved regulatory elements at the nou-darake gene (ndk), showing H3K4me3 ChIP, H3K27ac ChIP, and ATAC-seq signal for *S. mediterranea* on the top, followed by a track indicating the mapping of conserved non-coding elements identified in Dugesia japonica and below the ATAC-seq signal in the genomes of *S. polychroa* (schPol2), *S. nova* (schNov1), and *S. lugubris* (schLug1). Blue lines indicate conserved regulatory elements.

### 4 Synteny

#### 4.1 GENESPACE based synteny

Table 7 Syntenic blocks identified between schMedS3h1 and the other assemblies using GENESPACE. Given is the number of syntenic blocks with their median, standard deviation, minimum size, maximum size, and the genome coverage in Mb and as a percentage.

| genome | blocks | median (Mb) | SD (Mb) | min (Mb) | max (Mb) | block coverage (Mb) | block coverage (%) |
|---|---|---|---|---|---|---|---|
| schMedS3h2 | 18 | 12.5 | 59.1 | 0.7 | 192.9 | 797.8 | 97.2 |
| schPol2 | 198 | 2.2 | 4.3 | 0.1 | 33.4 | 752.4 | 96.3 |
| schNov1 | 166 | 2.1 | 5.8 | 0.2 | 33.4 | 747.5 | 59.7 |
| schLug1 | 272 | 1.5 | 3.2 | 0.1 | 26.7 | 730.8 | 48.7 |

#### 4.2 Synteny breakpoint inspection

Table 8 Enrichment analysis of 10kb windows flanking synteny breakpoints inferred using GENESPACE. Tests are based on comparisons to 1000 iterations of random placement of an equal number of 10kb windows in the reference. Given are results for all transposable elements followed by the LTR/Gypsy and LINE/R2 elements since they returned the most relevant results. For a full table including all tested elements, see Additional File 2: Table S12. P values are adjusted for multiple testing using the Benjamini-Hochberg procedure and printed in bold when <0.05.

| reference | target | observed | expected | SD | padj_larger |
|---|---|---|---|---|---|
| schMedS3h1 | schLug1 | 8387 | 7065 | 204 | **0.000** |
| schMedS3h1 | schMedS3h2 | 921 | 621 | 56 | **0.007** |
| schMedS3h1 | schNov1 | 5035 | 4414 | 157 | **0.008** |
| schMedS3h1 | schPol2 | 6288 | 5283 | 182 | **0.000** |
| schMedS3h2 | schMedS3h1 | 960 | 668 | 63 | **0.011** |
| schPol2 | schMedS3h1 | 5913 | 5435 | 152 | **0.000** |
| schNov1 | schMedS3h1 | 6007 | 6319 | 474 | 0.998 |
| schLug1 | schMedS3h1 | 9884 | 9503 | 256 | 0.303 |
| LTR/Gypsy | | | | | |
| schMedS3h1 | schLug1 | 1741 | 1277 | 58 | **0.000** |
| schMedS3h1 | schMedS3h2 | 164 | 110 | 16 | **0.020** |
| schMedS3h1 | schNov1 | 1059 | 792 | 43 | **0.000** |
| schMedS3h1 | schPol2 | 1269 | 958 | 51 | **0.000** |
| schMedS3h2 | schMedS3h1 | 148 | 99 | 16 | **0.033** |
| schPol2 | schMedS3h1 | 1533 | 1320 | 64 | **0.000** |
| schNov1 | schMedS3h1 | 1980 | 1799 | 82 | 0.076 |
| schLug1 | schMedS3h1 | 2600 | 2275 | 76 | **0.000** |
| LINE/R2 | | | | | |
| schMedS3h1 | schLug1 | 1249 | 664 | 52 | **0.000** |
| schMedS3h1 | schMedS3h2 | 103 | 59 | 14 | **0.007** |
| schMedS3h1 | schNov1 | 777 | 412 | 40 | **0.000** |
| schMedS3h1 | schPol2 | 1024 | 506 | 45 | **0.000** |
| schMedS3h2 | schMedS3h1 | 90 | 62 | 16 | 0.180 |
| schPol2 | schMedS3h1 | 22 | 20 | 11 | 0.708 |
| schNov1 | schMedS3h1 | 52 | 25 | 6 | 0.057 |
| schLug1 | schMedS3h1 | 9 | 11 | 5 | 0.995 |

### 4.3 Orthofinder based synteny

As described in the main text, we used single-copy orthologs derived using Orthofinder to test for synteny conservation using dotplots and Chi-square tests. The results for the comparison of *S. mediterranea* haplotype 1 against the other *Schmidtea* and the parasites as well as *Schistosoma mansoni* against the same are shown below (Table 9-10). Additionally, we compared *Schmidtea* and the parasites to Amphioxus, a representative of ancestral vertebrate linkage groups. This revealed that while we can replicate previous findings that its synteny is conserved with the cnidarian *Nematostella vectensis*, it is lost compared to members of the genus *Schmidtea* (Table 11). Interestingly, macrosynteny between Amphioxus and *S. mansoni* and the other parasites was degraded with all effect sizes <= 0.22 (Table 11), but some synteny appeared conserved for specific chromosomes (e.g., Amphioxus chromosomes 3 and 4).

Table 9 Test for macrosynteny using a chi-square test of the distribution of one-to-one orthologs between the target and query genome assemblies. Given is the number of orthologs, the degrees of freedom, the chi-squared statistic, the effect size Cramer's V which is 0 for a random distribution and 1 for a perfect correlation and the p-value calculated assuming the chi-square distribution, or 100,000 permutations of the data.

| target | query | orthologs | df | chisq | V | p-val | perm. p-val |
|---|---|---|---|---|---|---|---|
| schMedS3h1 | schMedS3h2 | 11718 | 9 | 34436.9 | 0.99 | **<1e-16** | **< 1e-05** |
| schMedS3h1 | schLug1 | 9479 | 9 | 10233.4 | 0.6 | **<1e-16** | **< 1e-05** |
| schMedS3h1 | schPol2 | 9458 | 9 | 18508.7 | 0.81 | **<1e-16** | **< 1e-05** |
| schMedS3h1 | schNov1 | 9321 | 6 | 7150.1 | 0.62 | **<1e-16** | **< 1e-05** |
| schMedS3h1 | cloSin | 3440 | 18 | 43.1 | 0.06 | **0.000774** | **0.00087** |
| schMedS3h1 | schMan | 2602 | 27 | 45.3 | 0.08 | 0.0151 | 0.015 |
| schMedS3h1 | hymMic | 2996 | 15 | 15.2 | 0.04 | 0.438 | 0.44052 |
| schMedS3h1 | taeMul | 2823 | 15 | 39.3 | 0.07 | **0.000569** | **0.00055** |

Table 10 Test for macrosynteny using a chi-square test of the distribution of one-to-one orthologs between the target and query genome assemblies. Given is the number of orthologs, the degrees of freedom, the chi-squared statistic, the effect size Cramer's V which is 0 for a random distribution and 1 for a perfect correlation and the p-value calculated assuming the chi-square distribution, or 100,000 permutations of the data.

| target | query | orthologs | df | chisq | V | p-val | perm. p-val |
|---|---|---|---|---|---|---|---|
| schMan | schMedS3h2 | 2587 | 27 | 46.9 | 0.08 | 0.0102 | 0.01036 |
| schMan | schPol2 | 2555 | 27 | 43.1 | 0.07 | 0.0255 | 0.02538 |
| schMan | schNov1 | 2506 | 18 | 24.4 | 0.07 | 0.142 | 0.13971 |
| schMan | schLug1 | 2572 | 27 | 54.8 | 0.08 | 0.0012 | 0.00129 |
| schMan | cloSin | 3075 | 54 | 10406.1 | 0.75 | <1e-16 | < 1e-05 |
| schMan | hymMic | 2547 | 45 | 7789.5 | 0.78 | <1e-16 | < 1e-05 |
| schMan | taeMul | 2365 | 45 | 6998.4 | 0.77 | <1e-16 | < 1e-05 |

Table 11 Test for macrosynteny using a chi-square test of the distribution of one-to-one orthologs between the target and query genome assemblies. Given is the number of orthologs, the degrees of freedom, the chi-squared statistic, the effect size Cramer's V which is 0 for a random distribution and 1 for a perfect correlation and the p-value calculated assuming the chi-square distribution, or 100,000 permutations of the data.

| target | query | orthologs | df | chisq | V | p-val | perm. p-val |
|---|---|---|---|---|---|---|---|
| BraLan | NemVec | 5624 | 252 | 32122.4 | 0.64 | **<1e-16** | < 1e-05 |
| BraLan | schMedS3h1 | 3918 | 54 | 75.5 | 0.08 | **0.0282** | 0.0282 |
| BraLan | schMedS3h2 | 3890 | 54 | 77 | 0.08 | **0.0217** | 0.0210 |
| BraLan | schPol2 | 3788 | 54 | 67.2 | 0.08 | 0.107 | 0.1061 |
| BraLan | schNov1 | 3760 | 36 | 41.6 | 0.07 | 0.241 | 0.2403 |
| BraLan | schLug1 | 3831 | 54 | 67.7 | 0.08 | 0.0994 | 0.0993 |
| BraLan | cloSin | 3289 | 108 | 988.1 | 0.22 | <1e-16 | < 1e-05 |
| BraLan | schMan | 2577 | 162 | 1060.4 | 0.21 | <1e-16 | < 1e-05 |
| BraLan | hymMic | 2953 | 90 | 575 | 0.2 | <1e-16 | < 1e-05 |
| BraLan | taeMul | 2883 | 90 | 626.6 | 0.21 | <1e-16 | < 1e-05 |

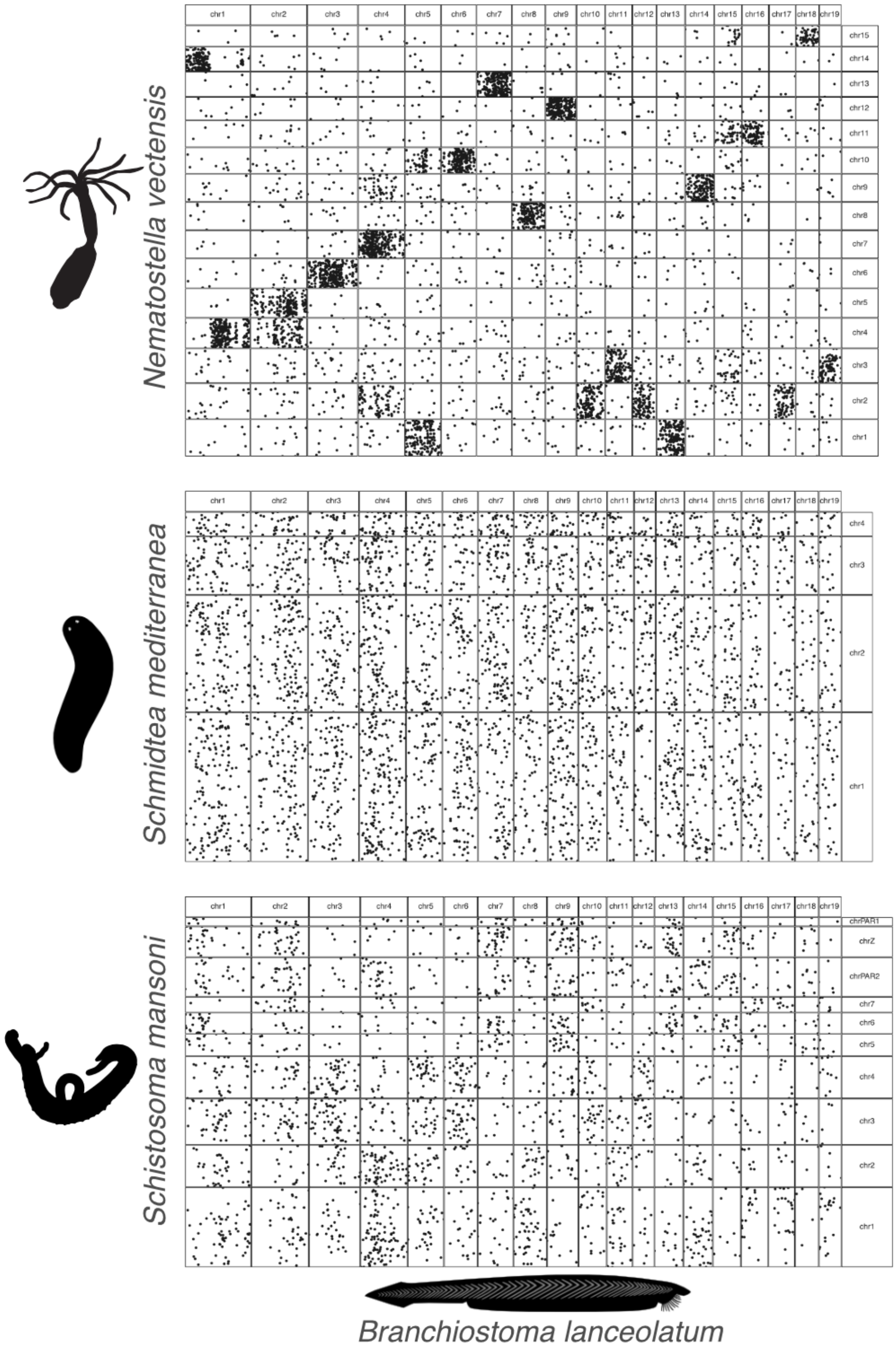


Figure 11 Oxford dotplot showing orthologous genes between Amphioxous, a representative of vertebrate ancestral linkage groups and the cnidarian *Nematostella vectensis*, *Schmidtea mediterranea* haplotype 1, and the parasitic flatworm *Schistosoma mansoni*.

### 4.4 ODP based synteny

To evaluate the synteny among the investigated genomes, we utilized the ODP tool. In the comparisons among the parasites, there was a marked conservation of synteny, as evidenced by significant p-values from Fisher's exact tests (Table 12). This conservation was evident in the pronounced clustering of orthologs by chromosomes, and even within chromosomal segments. Similarly, within *Schmidtea*, we observed substantial synteny conservation, with by significant p-values from Fisher's exact tests (Table 13) and the distinct clustering of orthologs by chromosome and gene order.
However, in pairwise comparisons between *Schmidtea* and the parasites, there was no evident clustering of orthologs by chromosomes (Table 14). The results of Fisher's exact tests indicated no association for cloSin. For taeMul, an association was observed between LG6 and chromosome 2 of both schNov1 and schLug1, though this was based on a limited gene set and was accompanied by a high p-value (Table 14). For hymMic, a more pronounced association was found: scaffold HMN_01_pilon aligned with a chromosome in schMedS3h1, schNov1, and schPol2, while HMN_06_pilon was associated with chr4 in schMedS3h1. Despite these associations involving a greater number of genes, the p-values remained elevated, consistent with the observed lack of clear clustering in the dotplot. For schMan a similar picture emerges but with statistically significant associations of at least one chromosome with each *Schmidtea* species (Table 14). Lastly, there was no significant association of any Machtxv2 scaffold with a chromosome of any other species included in this study.

Table 12 Results of Fisher's exact tests for synteny conservation between the parasite genomes in the study.

| genome1 | genome2 | scaf1 | scaf2 | p-value | count |
|---|---|---|---|---|---|
| cloSin | hymMic | 1 | HMN_01_pilon | 7.66E-243 | 1177 |
| cloSin | hymMic | 1 | HMN_03_pilon | 9.36E-216 | 664 |
| cloSin | hymMic | 2 | HMN_04_pilon | 2.42E-133 | 610 |
| cloSin | hymMic | 2 | HMN_02_pilon | 1.53E-76 | 475 |
| cloSin | hymMic | 2 | HMN_05_pilon | 9.67E-72 | 447 |
| cloSin | hymMic | 1 | HMN_06_pilon | 3.46E-11 | 303 |
| cloSin | hymMic | 3 | HMN_04_pilon | 1.53E-159 | 238 |
| cloSin | hymMic | 5 | HMN_02_pilon | 5.00E-169 | 232 |
| cloSin | hymMic | 4 | HMN_01_pilon | 3.53E-90 | 228 |
| cloSin | hymMic | 6 | HMN_05_pilon | 6.59E-158 | 212 |
| cloSin | hymMic | 7 | HMN_06_pilon | 1.11E-148 | 170 |
| cloSin | schMan | 1 | SM_V9_1 | 1.15E-29 | 905 |
| cloSin | schMan | 2 | SM_V9_3 | 0.00E+00 | 820 |
| cloSin | schMan | 1 | SM_V9_2 | 1.92E-294 | 817 |
| cloSin | schMan | 1 | SM_V9_4 | 7.26E-265 | 787 |
| cloSin | schMan | 2 | SM_V9_ZSR | 0.00E+00 | 658 |

| | | | | | |
|---|---|---|---|---|---|
| cloSin | schMan | 2 | SM_V9_PAR2 | 1.56E-46 | 433 |
| cloSin | schMan | 1 | SM_V9_6 | 5.22E-123 | 410 |
| cloSin | schMan | 3 | SM_V9_1 | 9.05E-210 | 355 |
| cloSin | schMan | 5 | SM_V9_PAR2 | 2.69E-288 | 311 |
| cloSin | schMan | 6 | SM_V9_1 | 1.13E-180 | 305 |
| cloSin | schMan | 4 | SM_V9_5 | 0.00E+00 | 305 |
| cloSin | schMan | 7 | SM_V9_7 | 0.00E+00 | 260 |
| cloSin | schMan | 2 | SM_V9_PAR1 | 4.29E-75 | 163 |
| cloSin | taeMul | 1 | LG1 | 3.80E-78 | 1638 |
| cloSin | taeMul | 2 | LG3 | 7.86E-178 | 406 |
| cloSin | taeMul | 3 | LG1 | 9.76E-21 | 220 |
| cloSin | taeMul | 5 | LG1 | 2.75E-21 | 207 |
| cloSin | taeMul | 4 | LG5 | 1.56E-280 | 207 |
| cloSin | taeMul | 6 | LG4 | 9.03E-255 | 186 |
| cloSin | taeMul | 1 | LG7 | 6.68E-35 | 151 |
| cloSin | taeMul | 7 | LG2 | 8.63E-215 | 144 |
| cloSin | taeMul | 1 | LG6 | 5.15E-31 | 120 |
| hymMic | schMan | HMN_03_pilon | SM_V9_1 | 0.00E+00 | 675 |
| hymMic | schMan | HMN_04_pilon | SM_V9_3 | 0.00E+00 | 626 |
| hymMic | schMan | HMN_01_pilon | SM_V9_4 | 6.29e-313 | 602 |
| hymMic | schMan | HMN_01_pilon | SM_V9_2 | 1.03E-305 | 593 |
| hymMic | schMan | HMN_02_pilon | SM_V9_PAR2 | 0.00E+00 | 523 |
| hymMic | schMan | HMN_05_pilon | SM_V9_ZSR | 5.85E-159 | 314 |
| hymMic | schMan | HMN_06_pilon | SM_V9_6 | 1.90E-292 | 288 |
| hymMic | schMan | HMN_04_pilon | SM_V9_1 | 5.22E-03 | 274 |
| hymMic | schMan | HMN_01_pilon | SM_V9_5 | 2.31E-89 | 228 |
| hymMic | schMan | HMN_05_pilon | SM_V9_1 | 3.97E-05 | 227 |
| hymMic | schMan | HMN_02_pilon | SM_V9_ZSR | 4.26E-29 | 172 |
| hymMic | schMan | HMN_06_pilon | SM_V9_7 | 2.10E-137 | 159 |
| hymMic | schMan | HMN_05_pilon | SM_V9_PAR1 | 3.15E-65 | 104 |
| hymMic | taeMul | HMN_01_pilon | LG1 | 8.86E-24 | 1314 |
| hymMic | taeMul | HMN_04_pilon | LG1 | 2.96E-146 | 1001 |
| hymMic | taeMul | HMN_02_pilon | LG1 | 5.37E-105 | 828 |
| hymMic | taeMul | HMN_03_pilon | LG1 | 2.13E-104 | 776 |
| hymMic | taeMul | HMN_05_pilon | LG3 | 0.00E+00 | 527 |
| hymMic | taeMul | HMN_05_pilon | LG4 | 7.78E-227 | 283 |
| hymMic | taeMul | HMN_01_pilon | LG5 | 7.48E-143 | 282 |
| hymMic | taeMul | HMN_06_pilon | LG2 | 5.17E-259 | 213 |
| hymMic | taeMul | HMN_06_pilon | LG6 | 1.35E-197 | 168 |
| schMan | taeMul | SM_V9_1 | LG1 | 2.53E-05 | 850 |

| | | | | | |
|---|---|---|---|---|---|
| schMan | taeMul | SM_V9_3 | LG1 | 9.61E-69 | 542 |
| schMan | taeMul | SM_V9_2 | LG1 | 2.59E-62 | 533 |
| schMan | taeMul | SM_V9_4 | LG1 | 3.69E-62 | 532 |
| schMan | taeMul | SM_V9_PAR2 | LG1 | 3.04E-28 | 475 |
| schMan | taeMul | SM_V9_ZSR | LG3 | 3.09E-204 | 282 |
| schMan | taeMul | SM_V9_5 | LG5 | 6.99E-276 | 200 |
| schMan | taeMul | SM_V9_1 | LG4 | 1.14E-91 | 200 |
| schMan | taeMul | SM_V9_7 | LG2 | 4.91E-223 | 149 |
| schMan | taeMul | SM_V9_6 | LG6 | 3.96E-186 | 119 |
| schMan | taeMul | SM_V9_PAR1 | LG3 | 6.47E-74 | 92 |

Table 13 Results of Fisher's exact tests for synteny conservation between the Schmidtea genomes in the study. schMedS3h2 was omitted for readability since it had qualitatively identical results to schMedS3h1.

| genome1 | genome2 | scaf1 | scaf2 | p-value | count |
|---|---|---|---|---|---|
| schLug1 | schMedS3h1 | chr2 | chr1_h1 | 0.00E+00 | 3104 |
| schLug1 | schMedS3h1 | chr1 | chr2_h1 | 1.56E-55 | 2508 |
| schLug1 | schMedS3h1 | chr3 | chr2_h1 | 0.00E+00 | 2051 |
| schLug1 | schMedS3h1 | chr2 | chr4_h1 | 0.00E+00 | 1413 |
| schLug1 | schMedS3h1 | chr1 | chr3_h1 | 2.66E-33 | 1403 |
| schLug1 | schMedS3h1 | chr4 | chr3_h1 | 0.00E+00 | 1069 |
| schLug1 | schNov1 | chr2 | chr2 | 0.00E+00 | 4621 |
| schLug1 | schNov1 | chr1 | chr1 | 0.00E+00 | 4410 |
| schLug1 | schNov1 | chr3 | chr1 | 0.00E+00 | 2076 |
| schLug1 | schNov1 | chr1 | chr3 | 0.00E+00 | 1681 |
| schLug1 | schNov1 | chr4 | chr2 | 0.00E+00 | 1033 |
| schLug1 | schPol2 | chr2 | chr3 | 0.00E+00 | 3096 |
| schLug1 | schPol2 | chr1 | chr1 | 5.44E-64 | 2563 |
| schLug1 | schPol2 | chr1 | chr4 | 0.00E+00 | 2320 |
| schLug1 | schPol2 | chr3 | chr1 | 0.00E+00 | 2055 |
| schLug1 | schPol2 | chr2 | chr2 | 3.67E-12 | 1476 |
| schLug1 | schPol2 | chr4 | chr2 | 0.00E+00 | 1056 |
| schMedS3h1 | schNov1 | chr2_h1 | chr1 | 0.00E+00 | 4495 |
| schMedS3h1 | schNov1 | chr1_h1 | chr2 | 2.05E-219 | 3168 |
| schMedS3h1 | schNov1 | chr1_h1 | chr3 | 0.00E+00 | 1664 |
| schMedS3h1 | schNov1 | chr4_h1 | chr2 | 0.00E+00 | 1404 |
| schMedS3h1 | schNov1 | chr3_h1 | chr1 | 3.67E-16 | 1311 |
| schMedS3h1 | schNov1 | chr3_h1 | chr2 | 8.32E-05 | 1084 |
| schMedS3h1 | schPol2 | chr2_h1 | chr1 | 0.00E+00 | 4619 |
| schMedS3h1 | schPol2 | chr1_h1 | chr3 | 0.00E+00 | 3172 |
| schMedS3h1 | schPol2 | chr3_h1 | chr2 | 0.00E+00 | 2515 |
| schMedS3h1 | schPol2 | chr1_h1 | chr4 | 0.00E+00 | 2342 |
| schMedS3h1 | schPol2 | chr4_h1 | chr2 | 0.00E+00 | 1439 |
| schMedS3h1 | schMedS3h2 | chr1_h1 | chr1_h2 | 0.00E+00 | 6670 |
| schMedS3h1 | schMedS3h2 | chr2_h1 | chr2_h2 | 0.00E+00 | 6171 |
| schMedS3h1 | schMedS3h2 | chr3_h1 | chr3_h2 | 0.00E+00 | 3458 |
| schMedS3h1 | schMedS3h2 | chr4_h1 | chr4_h2 | 0.00E+00 | 1646 |
| schNov1 | schPol2 | chr1 | chr1 | 0.00E+00 | 4537 |
| schNov1 | schPol2 | chr2 | chr3 | 0.00E+00 | 3160 |
| schNov1 | schPol2 | chr2 | chr2 | 1.67E-271 | 2488 |
| schNov1 | schPol2 | chr3 | chr4 | 0.00E+00 | 1634 |

Table 14 Results of Fisher's exact tests for synteny conservation between the Schmidtea genomes and the parasite genomes in the study. schMedS3h2 was omitted for readability since it had qualitatively identical results to schMedS3h1.

| genome1 | genome2 | scaf1 | scaf2 | p | count |
|---|---|---|---|---|---|
| cloSin | schLug1 | NA | NA | NA | NA |
| cloSin | schMedS3h1 | NA | NA | NA | NA |
| cloSin | schNov1 | NA | NA | NA | NA |
| cloSin | schPol2 | NA | NA | NA | NA |
| hymMic | schLug1 | NA | NA | NA | NA |
| hymMic | schMedS3h1 | HMN_01_pilon | chr3_h1 | 6.04E-03 | 259 |
| hymMic | schMedS3h1 | HMN_06_pilon | chr4_h1 | 6.03E-03 | 64 |
| hymMic | schNov1 | HMN_01_pilon | chr3 | 1.30E-02 | 171 |
| hymMic | schPol2 | HMN_01_pilon | chr4 | 1.09E-02 | 238 |
| schMan | schLug1 | SM_V9_5 | chr1 | 1.70E-02 | 130 |
| schMan | schLug1 | SM_V9_1 | chr2 | 2.09E-02 | 425 |
| schMan | schLug1 | SM_V9_6 | chr2 | 3.15E-05 | 129 |
| schMan | schLug1 | SM_V9_4 | chr3 | 1.48E-02 | 118 |
| schMan | schMedS3h1 | SM_V9_6 | chr4_h1 | 1.17E-05 | 57 |
| schMan | schNov1 | SM_V9_6 | chr2 | 4.63E-04 | 148 |
| schMan | schNov1 | SM_V9_4 | chr3 | 2.89E-02 | 93 |
| schMan | schPol2 | SM_V9_1 | chr3 | 4.01E-02 | 297 |
| schMan | schPol2 | SM_V9_5 | chr4 | 1.26E-04 | 66 |
| taeMul | schLug1 | LG6 | chr2 | 1.63E-02 | 47 |
| taeMul | schMedS3h1 | NA | NA | NA | NA |
| taeMul | schNov1 | LG6 | chr2 | 1.82E-02 | 55 |
| taeMul | schPol2 | NA | NA | NA | NA |

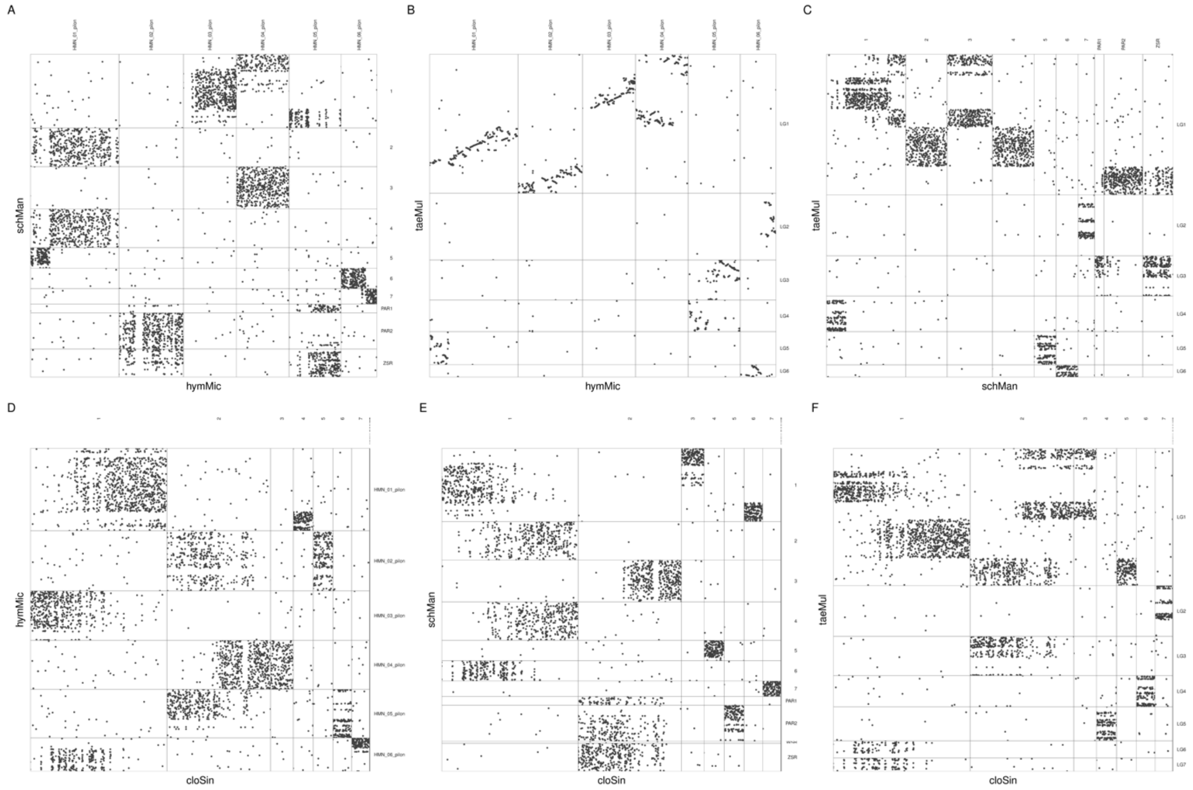


Figure 12 Oxford dotplot between the parasites species included in this study.

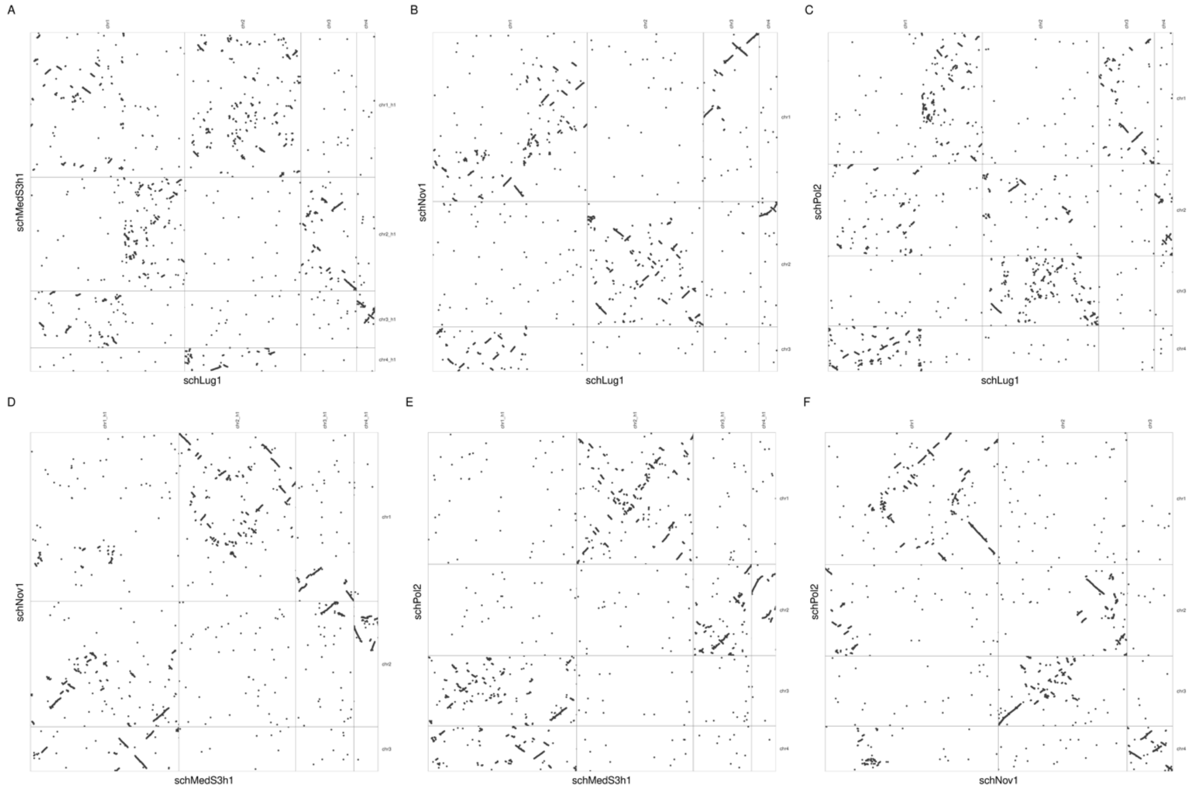


Figure 13 Oxford dotplot between the Schmidtea species included in this study.

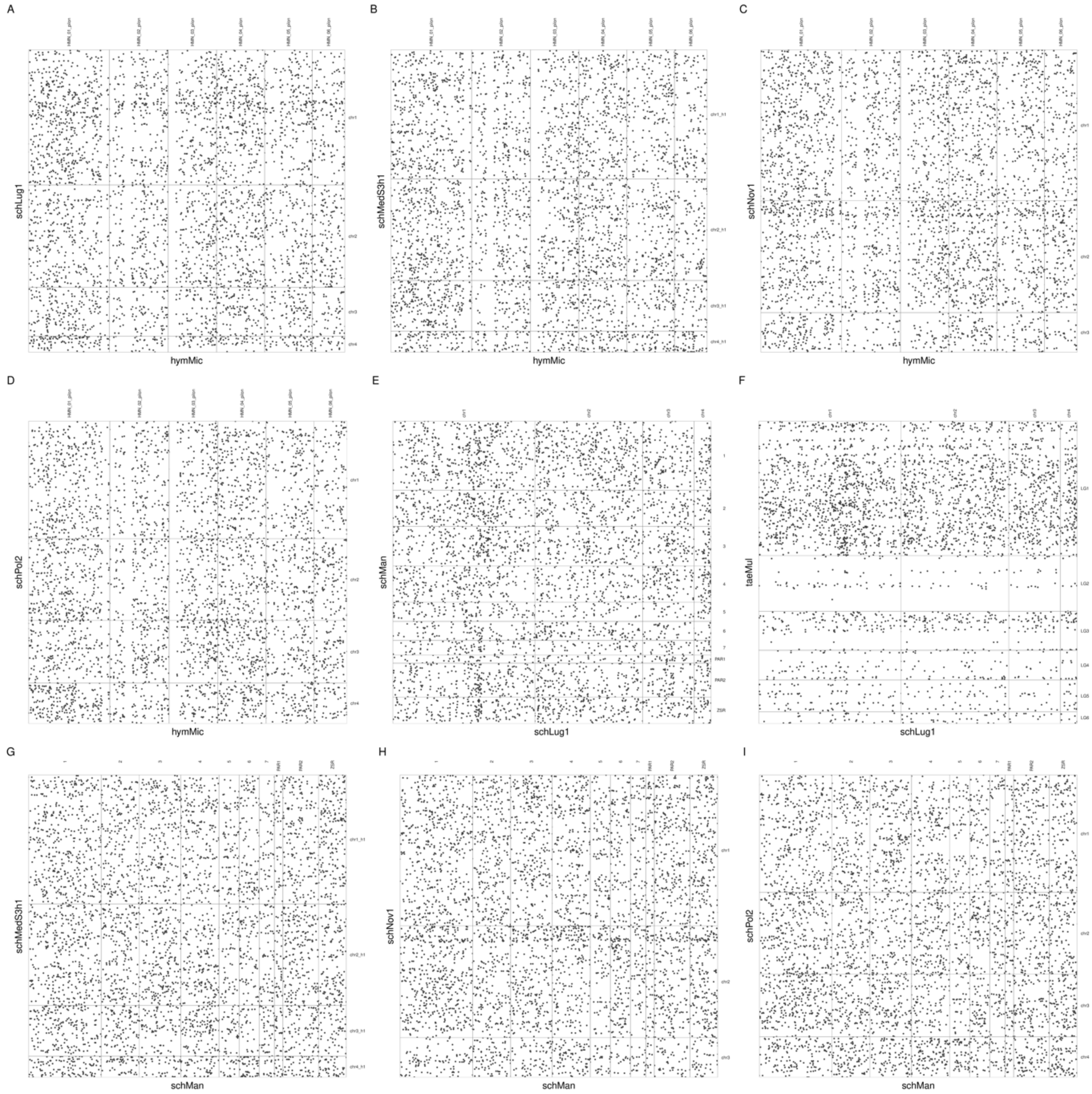


Figure 14 Oxford dotplot between the Schmidtea and parasites species included in this study.

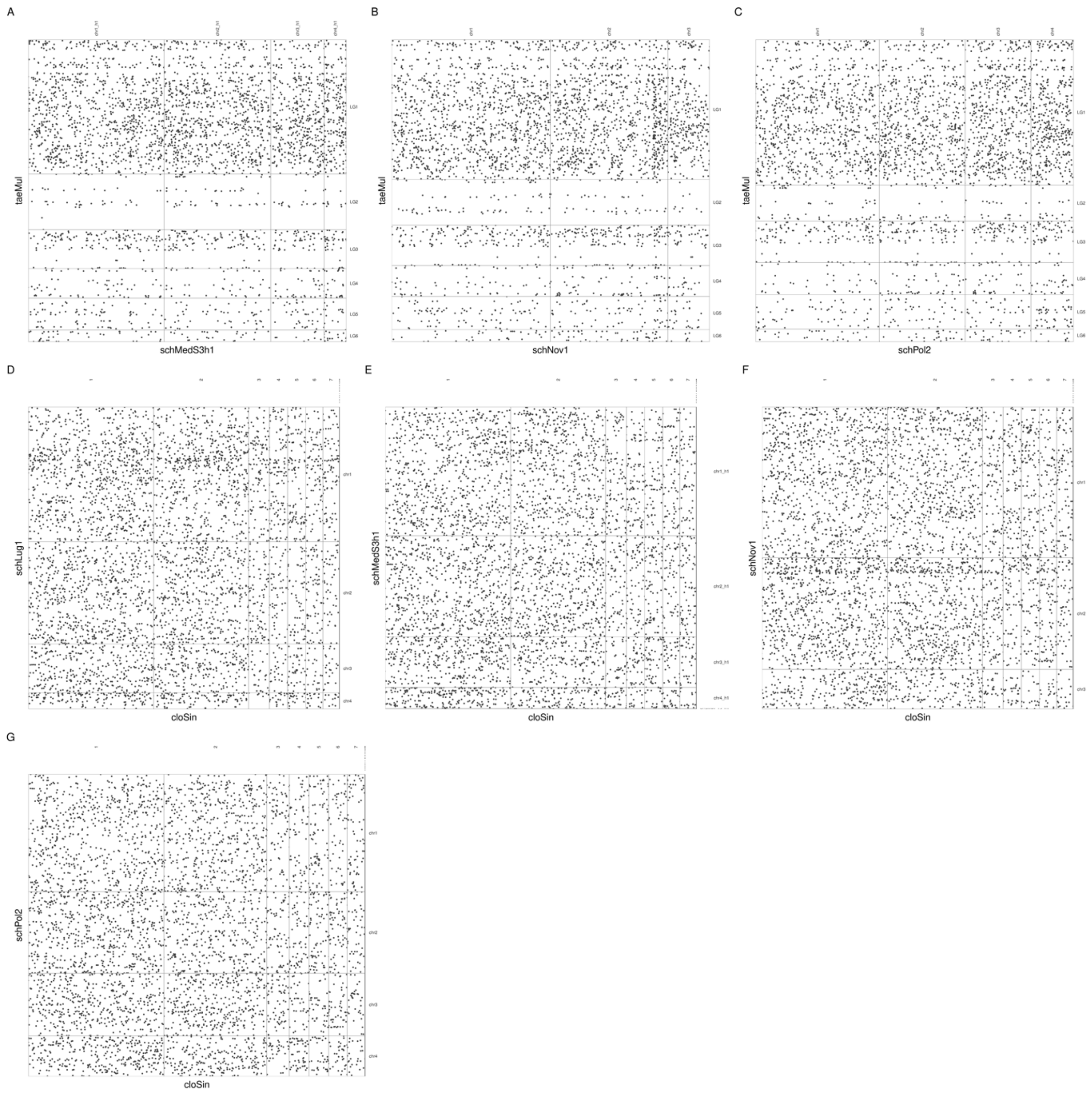


Figure 15 Oxford dotplot between the Schmidtea and parasites species included in this study.

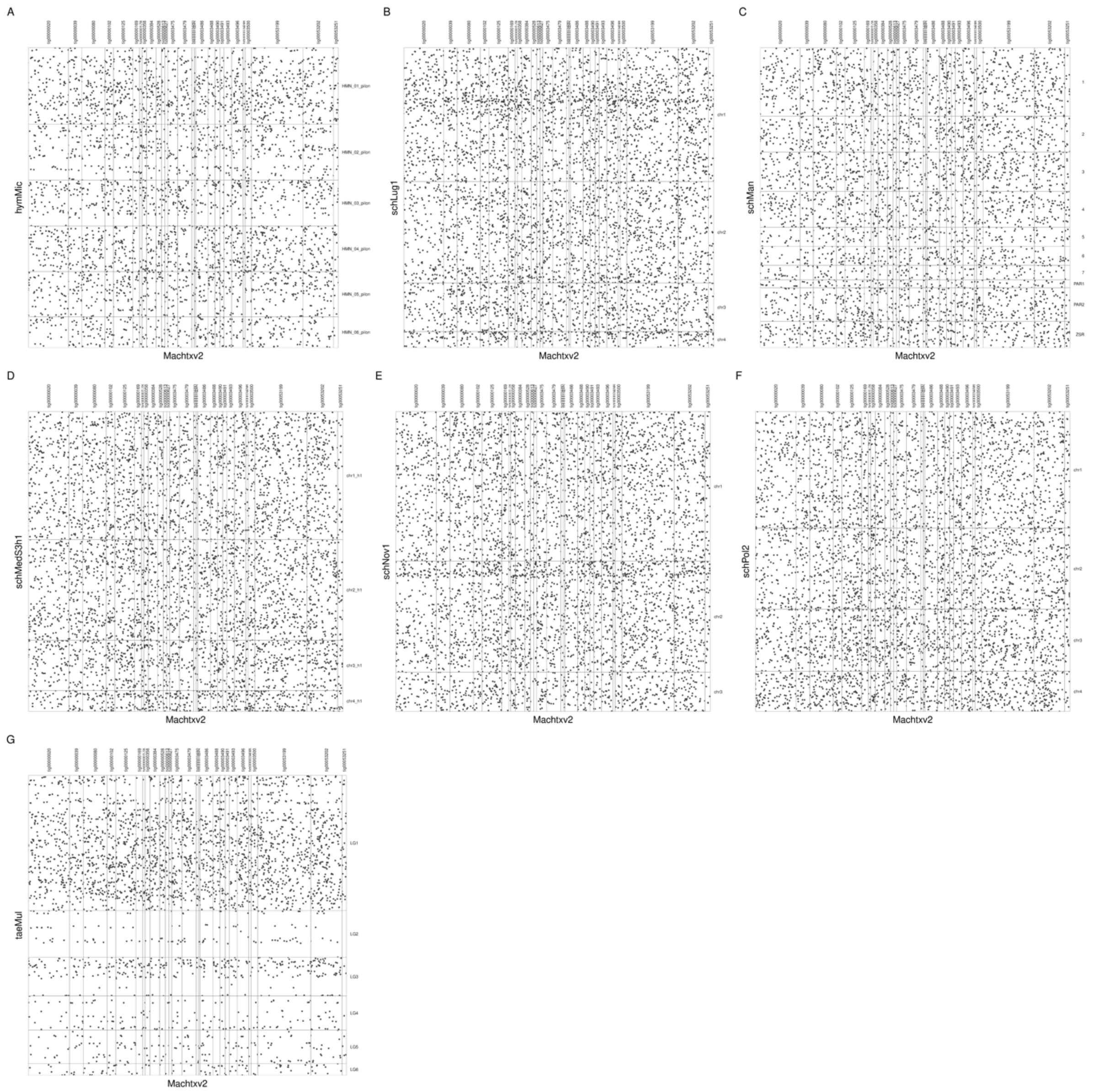


Figure 16 Oxford dotplot between *Macrostomum hystrix* and the *Schmidtea* and parasites species included in this study.

### 4.5 MALG conservation

Here we describe the conservation of the Metazoan ancestral linkage groups (MALG) defined by [3]. Following their notation, we use ⊗ to denote the fusion and mixing of two MALG and describe the equivalence of chromosomes. Note, that none of the MALG were conserved in *S. mediterranea*, *S. polychroa*, *S. nova, S. lugubris* (Figure 17A-D), or *Macrostomum hystrix* (Figure 14I) and therefore they are not included in this description. Table 15 gives the conserved MALGs for each chromosome based on the ODP results visualized in Figure 17E-H.

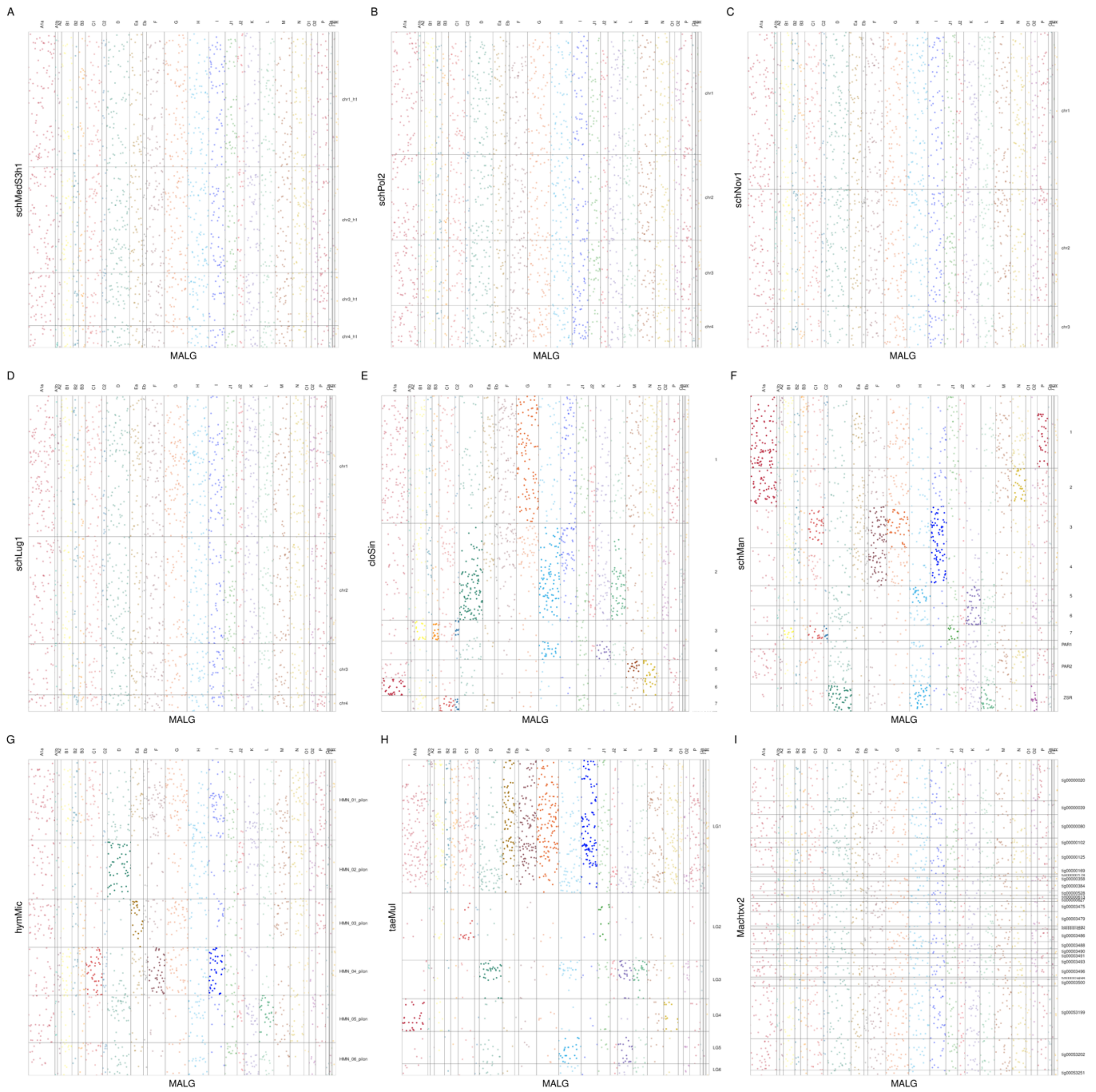


Figure 17 Oxford dotplots between MALG and the species in this study.

Table 15 Description of conservation of metazoan Ancestral Linkage Groups in four parasitic flatworms. Fusion and mixing of Ancestral Linkage Groups are denoted by ⊗.

| species | scaffold | MALG | scaffold name |
|---|---|---|---|
| schMan | 1 | A1a ⊗ P | SM_V9_1 |
| schMan | 2 | A1a ⊗ N | SM_V9_2 |
| schMan | 3 | G ⊗ F ⊗ I ⊗ C1 | SM_V9_3 |
| schMan | 4 | F ⊗ I | SM_V9_4 |
| schMan | 5 | H ⊗ K | SM_V9_5 |
| schMan | 6 | K | SM_V9_6 |
| schMan | 7 | C1 ⊗ C2 ⊗ J1 | SM_V9_7 |
| schMan | PAR1 | — | SM_V9_PAR1 |
| schMan | PAR2 | — | SM_V9_PAR2 |
| schMan | WSR | — | SM_V9_WSR |
| schMan | ZSR | D ⊗ H ⊗ L ⊗ O | SM_V9_ZSR |
| cloSin | 1 | G ⊗ F ⊗ I ⊗ C1 | 1 |
| cloSin | 2 | D ⊗ H ⊗ L | 2 |
| cloSin | 3 | B1 ⊗ B3 ⊗ C2 | 3 |
| cloSin | 4 | H ⊗ K | 4 |
| cloSin | 5 | M ⊗ N | 5 |
| cloSin | 6 | A1a ⊗ N | 6 |
| cloSin | 7 | C1 ⊗ C2 | 7 |
| taeMul | 1 | Ea ⊗ Eb ⊗ G ⊗ F ⊗ I | LG1 |
| taeMul | 2 | C1 ⊗ C2 ⊗ J1 | LG2 |
| taeMul | 3 | D ⊗ K ⊗ L | LG3 |
| taeMul | 4 | A1a ⊗ N | LG4 |
| taeMul | 5 | H ⊗ K | LG5 |
| taeMul | 6 | — | LG6 |
| hymMic | 1 | — | HMN_01_pilon |
| hymMic | 2 | D | HMN_02_pilon |
| hymMic | 3 | Ea | HMN_03_pilon |
| hymMic | 4 | F ⊗ I ⊗ C1 | HMN_04_pilon |
| hymMic | 5 | L | HMN_05_pilon |
| hymMic | 6 | — | HMN_06_pilon |
