## Additional File 3 for "A comparative analysis of planarian genomes reveals regulatory conservation in the face of rapid structural divergence"

### Part I. Transcriptome assembly pipeline

The following workflow describes the complete transcriptome assembly pipeline employed in order to assign the transcript loci of the new genome releases. It describes an evidence-based (*i.e.* relying on RNA-sequencing data), genome-guided approach, and it is summarised in the following scheme. A detailed block diagram of the pipeline is attached at the end of the document.

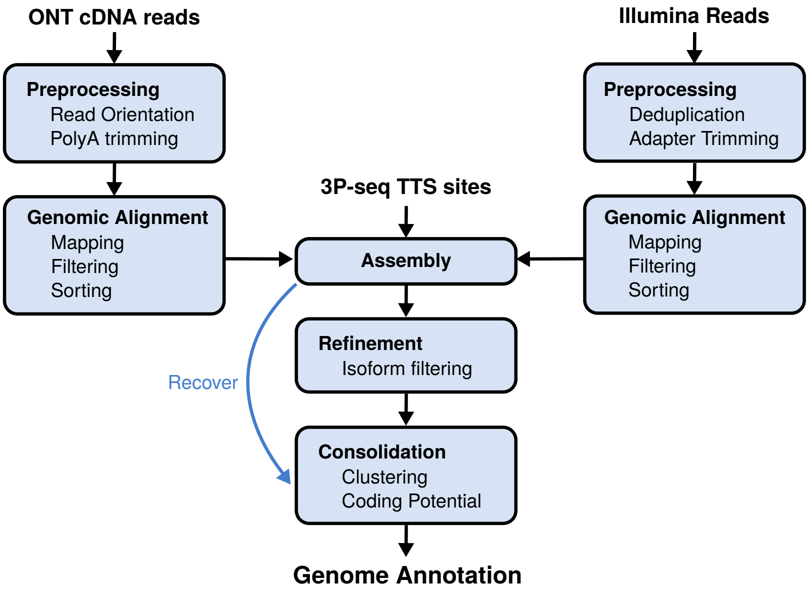


**Figure 1:** Overall workflow

#### 1. Nanopore Read Pre-processing

##### 1.1 Basecalling

ONT reads are obtained by basecalling `fast5` files using **guppy** *(6.2.1)*:

```
guppy_basecaller -i $RUNID/fast5/ \
                 -s $RUNID/fastq/ \
                 --recursive --disable_pings \
                 --trim_strategy none \
                 --compress_fastq \
                 --config $MODEL \
                 --device "cuda:0"
```

The following neural network models are used for basecalling:

- *Super High Accuracy* mode for cDNA reads.

```
MODEL="dna_r9.4.1_450bps_sup_prom.cfg"
```

- *High Accuracy* mode for direct RNA reads.

```
MODEL="rna_r9.4.1_70bps_hac_prom.cfg"
```

Only the *pass* reads of each run (*i.e.* having a Q-score>10 for cDNA and >7 for direct RNA) are retained and merged into a single `fastq` file:

```
find $RUNID/fastq/pass/ -type f -name *.fastq.gz |\
xargs cat > $RUNID".sup.fq.gz"
```

##### 1.2 Read orientation

Raw ONT reads are oriented and adapter-trimmed according to their native strandedness using **py-chopper** *(v2.7.1)* :

```
pychopper -k PCS111 -r $RUNID"_classifier_report.pdf" -t $THREADS \
          $RUNID".sup.fq.gz" $RUNID"_flr.pre.fq"
```

This step is skipped for *direct RNA* reads, that are inherently stranded. In the previous step, as well as throughout this workflow, `$THREADS` indicates the number of available CPU threads.

##### 1.3 Read trimming and filtering

Oriented reads > 150 nt are retained, then poly-A tails are trimmed and low complexity (entropy) reads (arising from sequencing artifacts) are removed using **BBMap** *(38.87)* tools.

```
reformat.sh in=$RUNID.filt_flr.pre.fq out=stdout.fq minlength=150 qin=33 |\
bbduk.sh -Xmx50G in=stdin.fq out=stdout.fq trimpolya=5
         threads=$THREADS qin=33 int=f |\
bbduk.sh -Xmx50G in=stdin.fq out=$RUNID.filt_flr_nopolya.fq.gz entropy=0.2 \
         entropywindow=25 entropytrim=r threads=$THREADS qin=33 int=f
```

Finally, multiple runs are pooled into a single long-read file:

```
cat *.fq.gz > $SPECIES.filt_flr_nopolya.fq.gz
```

##### 1.4 Genomic alignment

Reads are aligned to the genome *fasta* using **minimap2** *(2.24)* and *single mapping* reads are saved in `bam` format. A maximum intron length of 100kb is allowed during mapping.

```
minimap2 -ax splice -G 100k -L -uf -t $THREADS \
          $SPECIES.genome.fa $SPECIES.filt_flr_nopolya.fq.gz |\
samtools view -b -F 2308 -o $SPECIES"_np.pre.bam"
```

While strongly ameliorated by the latest library preparation strategies (e.g. PCS111), it is still possible that a fraction of the mapping reads arise from an internal priming event (*i.e.* oligo-dT cDNA primer annealing to an A-rich sequence within the transcript). Such events are discarded using **seqkit** *(v0.15.0)* by filtering alignments according to the genomic content around the 3' end of the read:

```
seqkit bam -j $THREADS \
           -x -T 'AlnContext: {
                  Ref: $SPECIES.genome.fa,
                  LeftShift: -24,
                  RightShift: 24,
                  RegexEnd: "[Aa]{8,}",
                  Stranded: True,
                  Invert: True }' $SPECIES"_np.pre.bam" > $SPECIES"_np.bam"
```

Finally, alignment files are sorted and indexed with **samtools** *(1.16.1)*:

```
samtools sort -@ $THREADS -o $SPECIES"_np.sort.bam" $SPECIES"_np.bam"
samtools index  $SPECIES"_np.sort.bam"
```

#### 2. Illumina Short Read Pre-processing

##### 2.1 Adapter removal and quality trimming

Illumina Adapters and low quality regions are trimmed away from short reads using **BBduk** (from BBMap tools):

```
bbduk.sh in1=$RUNID"_R1.fq.gz" in2=$RUNID"_R2.fq.gz" \
         out1=$RUNID"_tmp_R1.fq.gz" out2=$RUNID"_tmp_R2.fq.gz" \
         ref=adapters ktrim=r k=23 mink=11 hdist=1 qtrim=rl \
         trimq=10 tpe tbo &> $RUNID.trim.log
```

##### 2.2 PCR duplicate removal

In order to maximise the information content of short reads and avoid redundant mappings, we remove putative PCR duplicates (*i.e.* identical reads) using **BBmap**:

```
clumpify.sh in1=$RUNID"_tmp_R1.fq.gz" in2=$RUNID"_tmp_R2.fq.gz" \
            out1=$RUNID"_clean_R1.fq.gz" out2=$RUNID"_clean_R2.fq.gz" \
            dedupe=t optical=f &> $RUNID.clump.log
```

##### 2.3 Read Length Filtering

FInally, reads > 40bp are filtered using **BBMap**:

```
reformat.sh in1=$RUNID"_R1.fq.gz" in2=$RUNID"_R2.fq.gz" \
            out1=$RUNID"_final_R1.fq.gz" out2=$RUNID"_final_R2.fq.gz" \
            minlength=40
```

In the case of single read libraries, the `in1|in2` and `out1|out2` arguments from paragraphs 2.1-2.3 are replaced with `in=` and `out=`, respectively.

##### 2.4 Read Merging by Library Type

Most of the datasets include:

- `ISR` (Inward paired-end, stranded: `first-strand`)
- `SR` (Single read, stranded: `first-strand`)
- `IU` (Inward paired-end, `unstranded`)
- `U` (Single read, `unstranded`)

Runs of the same type are merged into a single `fastq` file (read files `_R1` and `_R2` are kept separated for paired-end runs):

```
cd $SPECIES

# IU
cat IU/*R1.fq.gz > IU_R1.fq.gz
cat IU/*R2.fq.gz > IU_R2.fq.gz

# ISR
cat ISR/*R1.fq.gz > ISR_R1.fq.gz
cat ISR/*R2.fq.gz > ISR_R2.fq.gz

# U
cat U/*.fq.gz > U.fq.gz

# SR
cat SR/*.fq.gz > SR.fq.gz
```

*S. lugubris*, *S. polychroa* and *S. nova* transciptomes are assembled using `ISR` reads only.

##### 2.5 Short Read Genome Mapping

Short reads are mapped to the genome using **HISAT2** *(2.2.1)*, after generating indexes:

```
hisat2-build -p $THREADS $SPECIES.genome.fa $SPECIES.HISAT2_index
```

Genomic alignment is performed by allowing a maximum intron length of 100kb (identical to long reads), and only single mapping reads are retained in the final `bam` files:

```
cd $SPECIES

## ISR
hisat2 -x $SPECIES.HISAT2_index -1 ISR_final_R1.fq.gz -2 ISR_final_R2.fq.gz \
       --rna-strandness RF --max-intronlen 100000 --threads $THREADS |\
samtools view -h -f 0x2 |\
grep -P "^\@|NH:i:1$" |\
samtools view -h -b -o ISR_final.$SPECIES.bam

## IU
hisat2 -x $SPECIES.HISAT2_index -1 IU_final_R1.fq.gz -2 IU_final_R2.fq.gz \
       --max-intronlen 100000 --threads $THREADS |\
samtools view -h -f 0x2 |\
grep -P "^\@|NH:i:1$" |\
samtools view -h -b -o IU_final.$SPECIES.bam

## SR
hisat2 -x $SPECIES.HISAT2_index -U SR_final.fq.gz --rna-strandness RF \
       --max-intronlen 100000 --threads $THREADS |\
grep -P "^\@|NH:i:1$" |\
samtools view -h -b -o SR_final.$SPECIES.bam

## U
hisat2 -x $SPECIES.HISAT2_index -U U_final.fq.gz \
       --max-intronlen 100000 --threads $THREADS |\
grep -P "^\@|NH:i:1$" |\
samtools view -h -b -o U_final.$SPECIES.bam
```

Finally, alignment files from all library types of the same species are merged into a single file, sorted and indexed:

```
samtools merge -@ $THREADS  All_sreads.$SPECIES.bam \
                            SR_final.$SPECIES.bam ISR_final.$SPECIES.bam \
                            IU_final.$SPECIES.bam U_final.$SPECIES.bam

samtools sort -@ $THREADS -o All_sreads_sorted.$SPECIES.bam \
              All_sreads.$SPECIES.bam

samtools index All_sreads_sorted.$SPECIES.bam
```

#### 3. Preparation of RNA Poly-adenylation Site Data

Poly(A)-position profiling data for *S. mediterranea* (*Lakshmanan et al., 2016*; SRP070102) were re-analysed using the publicly available scripts: https://github.com/VairavanL/3PSeq_analysis.

It is important to notice that this pipeline strictly requires the input `fastq` files to be uncompressed and to have a `.fastq` extension (*i.e.* `.fq` or other suffixes are not accepted). Furthermore, if the chromosome names contain underscore symbols, the resulting `3P-seq_processed_filtered.bedcount` output files will be formatted incorrectly (the program replaces the `_` with a `TAB`), therefore it is necessary to restore the right format using `sed`, as shown below. Briefly, short reads are mapped to the genome using **Bowtie** *(1.3.1)*, and a series of Python scripts extract the 3'-end positions. Only sites with at least 3 supporting reads are considered a valid TTS.

```
# Pre-process short reads
./3Pseq_iniprocess.py -q 3P-seq.fastq

# Prepare Bowtie index
bowtie-build --threads $THREADS $SPECIES.genome.fa $SPECIES.genome

# Align reads
./alignment_trigger.py -q 3P-seq_processed.fastq \
                       -g $SPECIES.genome -c config.ini

# Fix the formatting issues (in this case, for haplotype h1):
sed 's/\th1/_h1/g' 3P-seq_processed_filtered.bedcount |\
sed 's/\tscaffold\t/_scaffold_/g' > 3P-seq_processed_filtered.$SPECIES.bedcount

# Collect read signal into peaks
./detect_peaks.py 3P-seq_processed_filtered.$SPECIES.bedcount 3 \
> "TTS_"$SPECIES.bed
```

Finally, a custom Rscript formats the `.bed` TTS file into a point-feature (.ptf) for subsequent analysis:

```
Rscript ptfFromBed.R "TTS_"$SPECIES.bed  # generates "TTS_"$SPECIES.ptf
```

#### 4. Transcriptome Assembly

##### 4.1 Draft transcriptome construction

Draft genomic annotations are generated by **Stringtie2** *(2.2.0)* using alignments derived either from:

- ONT long-read only

```
stringtie $SPECIES"_np.sort.bam" -v \
          --rf \
          -t -c 1.5 -f 0.02 -g 0 \
          -p $THREADS -m 200 -l N \
          --ptf "TTS_"$SPECIES.ptf \
          -o $SPECIES.NP.gtf
```

- ONT long + Illumina short reads

```
stringtie -v \
          --mix All_sreads_sorted.$SPECIES.bam $SPECIES"_np.sort.bam" \
          --rf \
          -c 1.5 -f 0.05 \
          -s 2.5 -g 0 \
          -p $THREADS \
          -m 200 \
          -l H \
          --ptf "TTS_"$SPECIES.ptf \
          -o $SPECIES.mix_ann.gtf
```

The former approach is better suited for reconstructing an overall robust (albeit slightly less sensitive) exon chaining, the latter for fine-grained gene structures at the expenses of a slightly higher proportion of artifacts (chimaeras, spurious transcripts, etc.). The threshold parameters have been fine-tuned to optimise the sensitivity versus specificity of these specific assemblies. Both steps incorporate Poly(A)-position from 3P-seq experiments, in the form of `.ptf` data.

The output derived by the two Stringtie2 branches is then combined into a low-confidence draft set of annotations `.mix_ann-V5M`:

```
stringtie --merge \
    -G $SPECIES.mix_ann.gtf \
    $SPECIES.NP.gtf \
    -o $SPECIES.mix_ann-V5M.gtf \
    -p $THREADS -F 0 -T 0 -f 0 -g 0
```

##### 4.2 Model refinement

The low confidence set of annotation is refined using **FLAIR** *(v1.5)*: this program is able to remove spurious antisenses, to trim chimaeras, refine the splice junctions and filter out potential artifacts retaining transcripts with a higher confidence. All these steps are performed by relying on ONT read evidence only:

```
# Create input bed12 files from alignments
python bam2Bed12.py -i $SPECIES"_np.sort.bam" > $SPECIES"_np.sort.bed12"

# Correct reads
flair correct -q $SPECIES"_np.sort.bed12" \
              -g $SPECIES.genome.fa \
              -f $SPECIES.mix_ann-V5M.gtf \
              -o $SPECIES \
              -t $THREADS

# Collapse reads and refine loci
flair collapse -g $SPECIES.genome.fa \
               -r $SPECIES.filt_flr_nopolya.fq.gz \
               -q $SPECIES"_all_corrected.bed" \
               -f $SPECIES.mix_ann-V5M.gtf \
               -o $SPECIES.mix_ann-V5f \
               -t $THREADS \
               --temp_dir /tmp/
```

FLAIR produces a higher confidence transcript set called `.mix_ann-V5M.gtf`

##### 4.3 Transcript recovery

`FLAIR` ONT-based refinement of the transcriptome is sometimes too harsh, resulting in cases of gene fragmentation and overall decrease in completeness. Therefore, a recovery strategy is enforced: if a locus was able to produce a longer protein before `FLAIR` filtering (roughly 10-15% of the cases), we include the transcript corresponding to that ORF (*i.e.* a transcript contained in `.mix_ann-V5M.gtf`) back into the high confidence annotation. In order to do so, we first need to obtain the coding products of the two transcript collections, using the `getORF.sh` wrapper script, built around **TransDecoder** *(v5.5.0)*:

```
getORFs.sh $SPECIES.mix_ann-V5M.gtf $SPECIES.genome.fa
getORFs.sh $SPECIES.mix_ann-V5f.isoforms.gtf $SPECIES.genome.fa
```

The script produces a `GFF3` annotation (`.CDS.gff3`), a list of *gene:transcript* name pairs (`.names`), and two `fasta` files containing the nucleotide (`.CDS.cds`) and amino acid (`.CDS.pep`) sequences, respectively, of the predicted Open Reading Frames longer than 100 amino acids.

Then we have to rename the `FLAIR` output transcripts (`V5f`) according to the nomenclature in `V5M` using **gffcompare** *(0.12.6)*:

```
gffcompare -o $SPECIES.MatchMissingID -r $SPECIES.mix_ann-V5M.gtf \
           $SPECIES.mix_ann-V5f.isoforms.gtf
```

Finally, we perform transcript recovery based on coding potential using the custom Rscript `fragmDetectFix.R`. This script:

- Imports both pre- and post-`FLAIR` annotations and ORF predictions, recovering only `FLAIR`'s *de novo* transcripts (`chr:XXXXX`) that map completely internally to a `V5M` transcript;
- Compares side by side the predicted protein products;
- Finds which transcripts have been fragmented after `FLAIR` refinement;
- Inserts back transcripts with longest ORF from `V5M` .

```
Rscript fragmDetectFix.R $SPECIES.mix_ann-V5M.gtf \
                         $SPECIES.mix_ann-V5M.CDS.gff3 \
                         $SPECIES.mix_ann-V5M.names \
                         $SPECIES.mix_ann-V5f.isoforms.gtf \
                         $SPECIES.mix_ann-V5f.isoforms.CDS.gff3
                         $SPECIES.mix_ann-V5f.isoforms.names \
                         $SPECIES.MatchMissingID.$SPECIES.mix_ann-V5f.isoforms.gtf.tmap \
                         $SPECIES.mix_ann-V5MF22.gtf
```

The result is the extended transcriptome annotation file `V5MF22.gtf` .

##### 4.4 Final transcript polishing

Some transcript at this stage still represent subportions of longer, full-length mRNAs. These are removed by collapsing them with **gffread** *(v0.12.7)*:

```
gffread -o $SPECIES.mix_ann-V5MF22C.gff3 \
        --merge -K -Y $SPECIES.mix_ann-V5MF22.gtf
```

Then, transcript are clustered into loci and renamed by genomic position into *gene-like* groups, using a custom script `tidyUpRename.sh` built as a wrapper around `gffread` :

```
#$TXROOT represents the intended suffix of transcript names (e.g. h1Smed).
tidyUpRename.sh $SPECIES.mix_ann-V5MF22C.gff3 \
                $SPECIES.mix_ann-V5MF22CN.gtf $TXROOT
```

### Part II. High-confidence transcript filtering

At this stage, the transcriptome is composed of a set of genes, each of which encompasses a broad collection of different isoforms. While many of these may represent true biological variants, a fraction of these is still made up by artifactual transcripts. We then proceed to filter out the obvious byproducts (e.g. intron retention events, transcript fragments, unlikely exon boundaries) in order to have a more succint, representative snapshot of the gene models.

#### 1. Isoform filtering

As a first step we perform a "soft" isoform filtering, *i.e.* we reduce the isoform number by removing artifactual splicing variants but retaining the ones with good protein coding potential support. In order to proceed, we first obtain the coding potential of the transcriptome:

```
getORFs.sh $SPECIES.mix_ann-V5MF22CN.gtf $SPECIES.fa
```

Then we use the custom script `filterIsoORF.R`, built as a wrapper around **CD-HIT** *(v4.8.1)*, that performs the following tasks:

- Import the genomic annotations and ORF content predictions;
- For each transcript:
  - Consider only its longest ORF and drop the other shorter putative ones;
- For each gene:
  - Run ORF (protein) clustering with `CD-HIT` (90% identity threshold);
  - Retain the top 2 transcripts with the longest ORF for each `CD-HIT` cluster, allowing ties;
  - In case of ties (>1 tx with identical ORF length), keep the 2 shortest transcripts (this is a conservative approach, since longer products may represent chimaeras or retained introns);
- Recover genes for which no ORF was predicted in any of its transcripts and divert them into a separate file (in order to preserve very short monoexonic genes or ncRNAs).

The script it is invoked with:

```
Rscript filterIsoORF.R \
            $SPECIES.mix_ann-V5MF22CN.gtf \
            $SPECIES.mix_ann-V5MF22CN.CDS.pep \
            $SPECIES.mix_ann-V5MF22CN_FiltIso.gtf \
            $SPECIES.mix_ann-V5MF22CN_nc.gtf \
            /path/to/cd-hit-binaries/
```

It produces:

- a filtered coding isoform file `V5MF22CN_FiltIso.gtf`
- a putative ncRNA file `V5MF22CN_nc.gtf`.

After this step, both files require a further round of locus/transcript renaming and clustering. Before running `tidyUpRename.sh` this time we need to parse these `.GTF` files into valid `GFF3` format (since `gffread` does not like the formatting produced by the `rtracklayer` library used in the R script):

```
# Do an intermediate gtf->gff3 conversion
gffread -o $SPECIES.mix_ann-V5MF22CN_FiltIso.gff3 \
           $SPECIES.mix_ann-V5MF22CN_FiltIso.gtf

gffread -o $SPECIES.mix_ann-V5MF22CN_nc.gff3 \
           $SPECIES.mix_ann-V5MF22CN_nc.gtf

# Tidy up and rename gene loci and transcripts
tidyUpRename.sh $SPECIES.mix_ann-V5MF22CN_FiltIso.gff3 \
                $SPECIES.mix_ann-V5MF22CN_FiltIsoN.gtf
                $SPECIES"c"

tidyUpRename.sh $SPECIES.mix_ann-V5MF22CN_nc.gff3 \
                $SPECIES.mix_ann-V5MF22CN_ncN.gtf
                $SPECIES"n"
```

This time, as a last argument, we specify (using the suffix `c` or `n`) whether the transcript belongs to the coding or non-coding set, respectively (e.g. `h1Smedc`, `Slugn`).

#### 2. Chimaeric transcript filtering

While this hybrid transcriptome assembly approach results in a lower proportion of transcript fusions, in the subsequent step we attempt to fix different kinds of remaining issues:

- **Chimaeric Loci**: *i.e.* Bundles of physically different but somewhat overlapping transcripts that are mistakenly assigned to the same Gene. This is the most common event.
- **Chimaeric Transcripts**: Transcripts that encode two distinct ORFs but are assembled as a single fusion transcript.
- **Mix of both**

The script `ChimaeraDetect.R` works as follow:

- For each locus:
  - Create genomic ranges with the `min` and `max` coordinates of all the predicted ORFs;
  - Reduce these coordinates by locus, creating *range_blocks* ;
  - If the locus has 0 or 1 range_block(s), this is a normal locus and it can be discarded from further analysis;

- For each chimaeric locus:
  - Map each range_block(s) to its overlapping transcript(s);
  - Transcripts encompassing more than one range_block are putative chimaeras;
  - When we manage to represent all range_blocks using non-chimeric transcript, we tag the chimeric transcript(s) from the locus as redundant. Otherwise, we keep it/them and flag it/them. In this way, if a locus harbors a chimeric transcript as well as their correct counterparts, only the redundant chimaera is removed. Conversely, in the rare event that the locus only contains a chimaera, and/or the removal of this transcript results in the loss of protein coding information, the transcript is preserved and tagged for future curation.
- Remove redundant chimaeric transcripts and rename loci accordingly (e.g. splitting chimeric loci into distinct ORF genes, according to their ORFs);

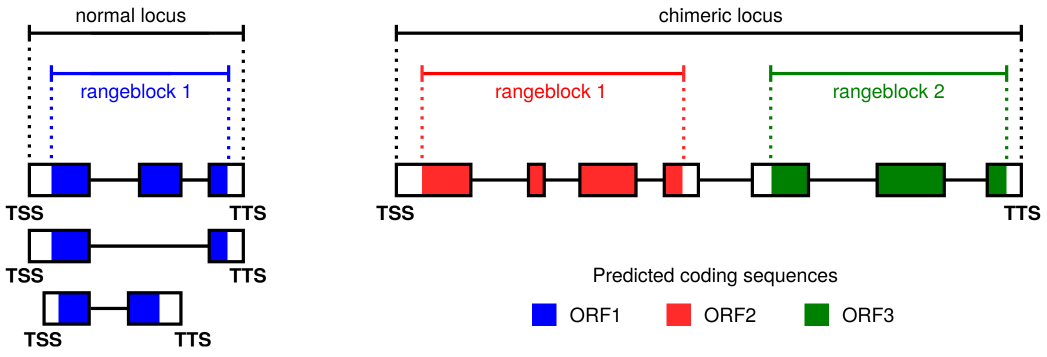


**Figure 2:** Normal and Chimaeric locus

ORF predictions are run before and after this step in order to identify and output the coding regions within the transcripts:

```
# The chimaera filtering steps require intermediate files, normally discarded
# after ORF predictions. With the "keeptmp" flag we retain them
getORFs.sh $SPECIES.mix_ann-V5MF22CN_FiltIsoN.gtf \
           $SPECIES.fa keeptmp

Rscript ChimaeraDetect.R $SPECIES.mix_ann-V5MF22CN_FiltIsoN.gtf \
                         $SPECIES.mix_ann-V5MF22CN_FiltIsoN.CDS.gff3 \
                         $SPECIES.mix_ann-V5MF22CN_FiltIsoN.rawCDS.gff3 \
                         $SPECIES.mix_ann-V5MF22CN_FiltIsoN_chfx.gtf \
                         $SPECIES"c"

# Coding predictions after chimaera removal
getORFs.sh $SPECIES.mix_ann-V5MF22CN_FiltIsoN_chfx.gtf \
           $SPECIES.fa
```

The script returns the `V5MF22CN_FiltIsoN_chfx.gtf` file containing an amended coding transcript annotation.

##### 3. Fix partial 5' ORF annotations

The ORF annotation predictions returned by TransDecoder classify the putative peptides as `complete`, `5-prime partial`, `3-prime partial`, or `internal`. However, most of the `5-prime partial` sequences contain a methionine relatively close to the predicted start of the ORF, and manual inspection (e.g. using BLAST on the full-length protein) confirms that a vast proportion of these are actual `complete` ORFs misassigned as partial fragments. In order to fix this problem we apply the following strategy:

- Import ORF annotations;
- Calculate the 5' UTR length distribution for `complete` ORFs (as a ground-truth reference);
- Estimate a threshold for 5' UTR length by fitting a Gaussian Mixture Model assuming a bimodal distribution with a majority of real, complete ORFs and a smaller, artifactual proportion of misassignments: the longest allowed lenght is therefore defined as 3 standard deviations above the mean of the lower population, as shown in the following picture;

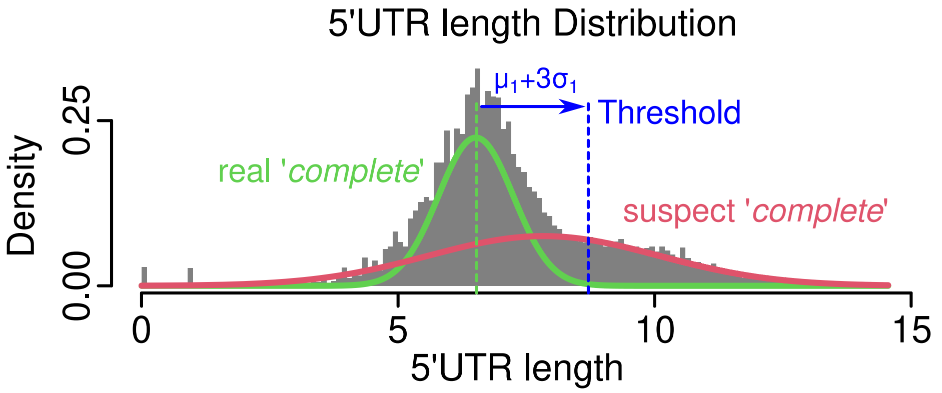


- `5-prime partial` ORFs with an in-frame methionine in their 5' UTR falling within the threshold distance are reclassified as `complete`, and their starting sites are corrected accordingly.

The above approach is implemented by the script `Fix5ORF.R`:

```
Rscript Fix5ORF.R $SPECIES.mix_ann-V5MF22CN_FiltIsoN_chfx.CDS.gff3 \
                  $SPECIES.fa \
                  $SPECIES.mix_ann-V5MF22CN_FiltIsoN_chfx.fix5utr.gff3
```

The script returns a `V5MF22CN_FiltIsoN_chfx.fix5utr.gff3` file containing an amended ORF prediction annotation.

In order to produce a compact ORF set, since TransDecoder may return multiple coding sequences per transcript, we finally run the script `filter_longestORFperTx.R`:

```
# Keep only 1 ORF (the longest) per tx
Rscript filter_longestORFperTx.R \
                $SPECIES.mix_ann-V5MF22CN_FiltIsoN_chfx.fix5utr.gff3 \
                $SPECIES.mix_ann-V5MF22CN_FiltIsoN_chfx.fix5utr.uniqueCDS.gff3

# Remove the ORF number (e.g. ".p1") in place
sed -i -E 's/\.p[0-9]+//g' \
    $SPECIES.mix_ann-V5MF22CN_FiltIsoN_chfx.fix5utr.uniqueCDS.gff3
```

#### 4. Final formatting

As a final step, we want to merge the coding and non-coding annotations into a single set:

```
# Convert non-coding set to gff3
gffread -F --keep-exon-attrs $SPECIES.mix_ann-V5MF22CN_ncN.gtf > \
                             $SPECIES.mix_ann-V5MF22CN_ncN.gff3

# Merge it with coding set
cat $SPECIES.mix_ann-V5MF22CN_FiltIsoN_chfx.fix5utr.uniqueCDS.gff3 \
    $SPECIES.mix_ann-V5MF22CN_ncN.gff3 |\
grep -v "#" > $SPECIES.ENCODE_hybrid_annot.tmp.gff3
```

The resulting `$SPECIES.ENCODE_hybrid_annot.tmp.gff3` annotation is then parsed using the `IDpatch.R` script. This command polishes the `GFF3` file by resolving format inconsistencies (e.g. adds explicit `gene` entries, renames the file `source`, fixes the `type` field layout):

```
Rscript IDpatch.R $SPECIES.ENCODE_hybrid_annot.tmp.gff3 \
                  $SPECIES.ENCODE_hybrid_annot.gff3
```

The following figure represents a description the overall naming scheme used in this annotation:

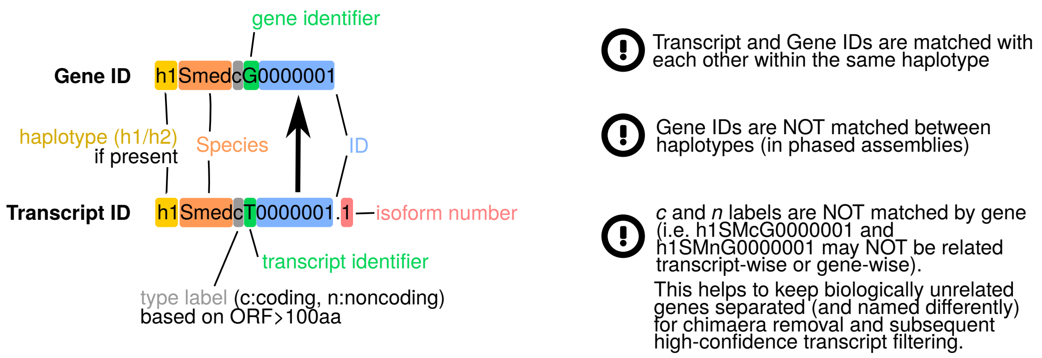


**Figure 3:** Gene naming scheme

In conclusion the file is sorted:

```
gffread --sort-alpha -F --keep-exon-attrs --keep-genes \
        $SPECIES.ENCODE_hybrid_annot.gff3 |\
grep -v "#" > $SPECIES.ENCODE_hybrid_annot.sort.gff3
```

This results in the final annotation file `ENCODE_hybrid_annot.sort.gff3`.

| Species | Sample | Reads (M) | Lib. Type | Chemistry | Flow Cell |
|---|---|---|---|---|---|
| *S.mediterranea S2F2* | Whole | 20.5 | cDNA | PCS109 | FLO-MIN106D |
| *S.mediterranea S2F2* | Heads | 16.7 | cDNA | PCS109 | FLO-MIN106D |
| *S.mediterranea S2F2* | Reg. Pool (7d) | 11.5 | cDNA | PCS109 | FLO-MIN106D |
| *S.mediterranea S2F2* | Reg. Pool (7d) | 13.9 | cDNA | PCS109 | FLO-MIN106D |
| *S.mediterranea S2F2* | Whole | 11.4 | dRNA | RNA002 | FLO-PRO002 |
| *S.mediterranea CIW4* | Heads | 12.3 | cDNA | PCS109 | FLO-MIN106D |
| *S.mediterranea CIW4* | Reg. Pool (7d) | 16.3 | cDNA | PCS109 | FLO-MIN106D |
| *S.mediterranea CIW4* | Heads | 5.9 | cDNA | PCS109 | FLO-MIN106D |
| *S.mediterranea CIW4* | Heads | 1.6 | dRNA | RNA002 | FLO-MIN106D |
| *S.mediterranea CIW4* | Whole | 18.8 | cDNA | PCS109 | FLO-MIN106D |
| *S.mediterranea CIW4* | Sorted Cells | 10.7 | cDNA | PCS109 | FLO-MIN106D |
| *S. lugubris* | Whole | 50.9 | cDNA | PCS111 | FLO-PRO002 |
| *S. lugubris* | Reg. Pool (7d) | 47.1 | cDNA | PCS111 | FLO-PRO002 |
| *S. lugubris* | Whole+Regen. | 7.3 | dRNA | RNA002 | FLO-PRO002 |
| *S. nova* | Whole | 54.9 | cDNA | PCS111 | FLO-PRO002 |
| *S. nova* | Reg. Pool (7d) | 40.1 | cDNA | PCS111 | FLO-PRO002 |
| *S. nova* | Whole+Regen. | 9.2 | dRNA | RNA002 | FLO-PRO002 |
| *S. polychroa* | Whole | 41.6 | cDNA | PCS111 | FLO-PRO002 |
| *S. polychroa* | Reg. Pool (7d) | 34.3 | cDNA | PCS111 | FLO-PRO002 |
| *S. polychroa* | Whole+Regen. | 10.1 | dRNA | RNA002 | FLO-PRO002 |

**Figure 4:** Summary of ONT reads used for the transcriptome assembly

| Run | BioProject | Center Name | Lab | Type | Rlen | Bases | Tissue | strain | LibType |
|---|---|---|---|---|---|---|---|---|---|
| SRR8162692 | PRJNA503908 | GEO | Kuhn | PAIRED | 300 | 2.6E+10 | whole animal | CIW4 | IU |
| SRR5876398 | PRJNA395927 | WRIGHT STATE UNIVERSITY | Rouhana | PAIRED | 250 | 3.5E+09 | heads | S2F2 | IU |
| SRR5876399 | PRJNA395927 | WRIGHT STATE UNIVERSITY | Rouhana | PAIRED | 250 | 3.0E+09 | heads | S2F2 | IU |
| SRR5876401 | PRJNA395927 | WRIGHT STATE UNIVERSITY | Rouhana | PAIRED | 250 | 3.0E+09 | heads | S2F2 | IU |
| SRR5876402 | PRJNA395927 | WRIGHT STATE UNIVERSITY | Rouhana | PAIRED | 250 | 4.0E+09 | heads | CIW4 | IU |
| SRR5876403 | PRJNA395927 | WRIGHT STATE UNIVERSITY | Rouhana | PAIRED | 250 | 3.5E+09 | heads | CIW4 | IU |
| SRR5876400 | PRJNA395927 | WRIGHT STATE UNIVERSITY | Rouhana | PAIRED | 250 | 3.6E+09 | heads | CIW4 | IU |
| SRR496276 | PRJNA167022 | GEO | Pearson | PAIRED | 202 | 4.4E+09 | FACS purified X1s | CIW4 | IU |
| SRR496277 | PRJNA167022 | GEO | Pearson | PAIRED | 202 | 4.3E+09 | FACS purified X1s | CIW4 | IU |
| SRR496278 | PRJNA167022 | GEO | Pearson | PAIRED | 202 | 2.1E+10 | FACS purified X2s | CIW4 | IU |
| SRR496279 | PRJNA167022 | GEO | Pearson | PAIRED | 202 | 2.1E+10 | FACS purified X2s | CIW4 | IU |
| SRR496280 | PRJNA167022 | GEO | Pearson | PAIRED | 202 | 4.2E+09 | D7 γ-irradiated | CIW4 | IU |
| SRR496281 | PRJNA167022 | GEO | Pearson | PAIRED | 202 | 4.2E+09 | D7 γ-irradiated | CIW4 | IU |
| SRR5680794 | PRJNA390190 | STOWERS INSTITUTE FOR MEDICAL RESEARCH | Alvarado | PAIRED | 200 | 8.7E+09 | D5RNA-R3 | S2F2 | IU |
| SRR5680795 | PRJNA390190 | STOWERS INSTITUTE FOR MEDICAL RESEARCH | Alvarado | PAIRED | 200 | 3.9E+09 | D0RNA-R1 | S2F2 | IU |
| SRR5680796 | PRJNA390190 | STOWERS INSTITUTE FOR MEDICAL RESEARCH | Alvarado | PAIRED | 200 | 7.0E+09 | D3RNA-R2 | S2F2 | IU |
| SRR5680797 | PRJNA390190 | STOWERS INSTITUTE FOR MEDICAL RESEARCH | Alvarado | PAIRED | 200 | 7.9E+09 | D3RNA-R3 | S2F2 | IU |
| SRR5680798 | PRJNA390190 | STOWERS INSTITUTE FOR MEDICAL RESEARCH | Alvarado | PAIRED | 200 | 1.1E+10 | D5RNA-R1 | S2F2 | IU |
| SRR5680799 | PRJNA390190 | STOWERS INSTITUTE FOR MEDICAL RESEARCH | Alvarado | PAIRED | 200 | 6.9E+09 | D5RNA-R2 | S2F2 | IU |
| SRR5680800 | PRJNA390190 | STOWERS INSTITUTE FOR MEDICAL RESEARCH | Alvarado | PAIRED | 200 | 9.2E+09 | D7RNA-R1 | S2F2 | IU |
| SRR5680801 | PRJNA390190 | STOWERS INSTITUTE FOR MEDICAL RESEARCH | Alvarado | PAIRED | 200 | 8.6E+09 | D7RNA-R2 | S2F2 | IU |
| SRR5680802 | PRJNA390190 | STOWERS INSTITUTE FOR MEDICAL RESEARCH | Alvarado | PAIRED | 200 | 9.6E+09 | D7RNA-R3 | S2F2 | IU |
| SRR5680803 | PRJNA390190 | STOWERS INSTITUTE FOR MEDICAL RESEARCH | Alvarado | PAIRED | 200 | 9.9E+09 | D3RNA-R1 | S2F2 | IU |
| SRR5680804 | PRJNA390190 | STOWERS INSTITUTE FOR MEDICAL RESEARCH | Alvarado | PAIRED | 200 | 6.6E+09 | D0RNA-R2 | S2F2 | IU |
| SRR5680805 | PRJNA390190 | STOWERS INSTITUTE FOR MEDICAL RESEARCH | Alvarado | PAIRED | 200 | 6.6E+09 | D0RNA-R3 | S2F2 | IU |
| SRR5680806 | PRJNA390190 | STOWERS INSTITUTE FOR MEDICAL RESEARCH | Alvarado | PAIRED | 200 | 4.9E+09 | Juv-R1 | S2F2 | IU |
| SRR5680807 | PRJNA390190 | STOWERS INSTITUTE FOR MEDICAL RESEARCH | Alvarado | PAIRED | 200 | 7.1E+09 | Juv-R2 | S2F2 | IU |
| SRR5680808 | PRJNA390190 | STOWERS INSTITUTE FOR MEDICAL RESEARCH | Alvarado | PAIRED | 200 | 5.3E+09 | Juv-R3 | S2F2 | IU |
| SRR748892 | PRJNA79997 | MAX DELBRUECK CENTER FOR MOLECULAR MEDICINE | Rajewsky | PAIRED | 152 | 5.0E+09 | whole animal | CIW4 | IU |
| SRR125365 | PRJNA79997 | MAX DELBRUECK CENTER FOR MOLECULAR MEDICINE | Rajewsky | PAIRED | 152 | 7.6E+03 | whole animal | CIW4 | IU |
| SRR125527 | PRJNA79997 | MAX DELBRUECK CENTER FOR MOLECULAR MEDICINE | Rajewsky | PAIRED | 152 | 2.1E+09 | whole animal | CIW4 | IU |
| SRR955099 | PRJNA215411 | STOWERS INSTITUTE FOR MEDICAL RESEARCH | Alvarado | PAIRED | 200 | 1.4E+08 | Pooled | CIW4 | ISR |
| SRR955511 | PRJNA215411 | STOWERS INSTITUTE FOR MEDICAL RESEARCH | Alvarado | PAIRED | 200 | 2.0E+08 | Pooled | S2F2 | ISR |
| SRR5408394 | PRJNA379262 | MPI OF MOLECULAR CELL BIOLOGY AND GENETICS | Rink | SINGLE | 91 | 4.6E+10 | Pooled | S2F2 | SR |
| SRR3465425 | PRJNA319973 | UNIVERSITY OF ILLINOIS, URBANA-CHAMPAIGN | Newmark | SINGLE | 100 | 1.4E+09 | Whole 1 | CIW4 | U |
| SRR3465432 | PRJNA319973 | UNIVERSITY OF ILLINOIS, URBANA-CHAMPAIGN | Newmark | SINGLE | 100 | 1.0E+09 | Whole 2 | CIW4 | U |
| SRR3465433 | PRJNA319973 | UNIVERSITY OF ILLINOIS, URBANA-CHAMPAIGN | Newmark | SINGLE | 100 | 2.3E+09 | Whole 3 | CIW4 | U |
| SRR3465434 | PRJNA319973 | UNIVERSITY OF ILLINOIS, URBANA-CHAMPAIGN | Newmark | SINGLE | 100 | 1.2E+09 | Cut 1 | CIW4 | U |
| SRR3465435 | PRJNA319973 | UNIVERSITY OF ILLINOIS, URBANA-CHAMPAIGN | Newmark | SINGLE | 100 | 2.1E+09 | Cut 2 | CIW4 | U |
| SRR3465502 | PRJNA319973 | UNIVERSITY OF ILLINOIS, URBANA-CHAMPAIGN | Newmark | SINGLE | 100 | 2.7E+09 | Cut 3 | CIW4 | U |
| SRR3465511 | PRJNA319973 | UNIVERSITY OF ILLINOIS, URBANA-CHAMPAIGN | Newmark | SINGLE | 100 | 1.2E+09 | 12h 1 | CIW4 | U |
| SRR3465512 | PRJNA319973 | UNIVERSITY OF ILLINOIS, URBANA-CHAMPAIGN | Newmark | SINGLE | 100 | 1.6E+09 | 12h 2 | CIW4 | U |
| SRR3465513 | PRJNA319973 | UNIVERSITY OF ILLINOIS, URBANA-CHAMPAIGN | Newmark | SINGLE | 100 | 1.3E+09 | 12h 3 | CIW4 | U |
| SRR3465518 | PRJNA319973 | UNIVERSITY OF ILLINOIS, URBANA-CHAMPAIGN | Newmark | SINGLE | 100 | 1.6E+09 | 36h 1 | CIW4 | U |
| SRR3465519 | PRJNA319973 | UNIVERSITY OF ILLINOIS, URBANA-CHAMPAIGN | Newmark | SINGLE | 100 | 9.8E+08 | 36h 2 | CIW4 | U |
| SRR3465520 | PRJNA319973 | UNIVERSITY OF ILLINOIS, URBANA-CHAMPAIGN | Newmark | SINGLE | 100 | 1.3E+09 | 72h 1 | CIW4 | U |
| SRR3465521 | PRJNA319973 | UNIVERSITY OF ILLINOIS, URBANA-CHAMPAIGN | Newmark | SINGLE | 100 | 1.7E+09 | 36h 3 | CIW4 | U |
| SRR3465522 | PRJNA319973 | UNIVERSITY OF ILLINOIS, URBANA-CHAMPAIGN | Newmark | SINGLE | 100 | 1.6E+09 | 72h 2 | CIW4 | U |
| SRR3465527 | PRJNA319973 | UNIVERSITY OF ILLINOIS, URBANA-CHAMPAIGN | Newmark | SINGLE | 100 | 9.7E+08 | 72h 3 | CIW4 | U |

**Figure 5:** Summary of short read datasets used for the transcriptome assembly

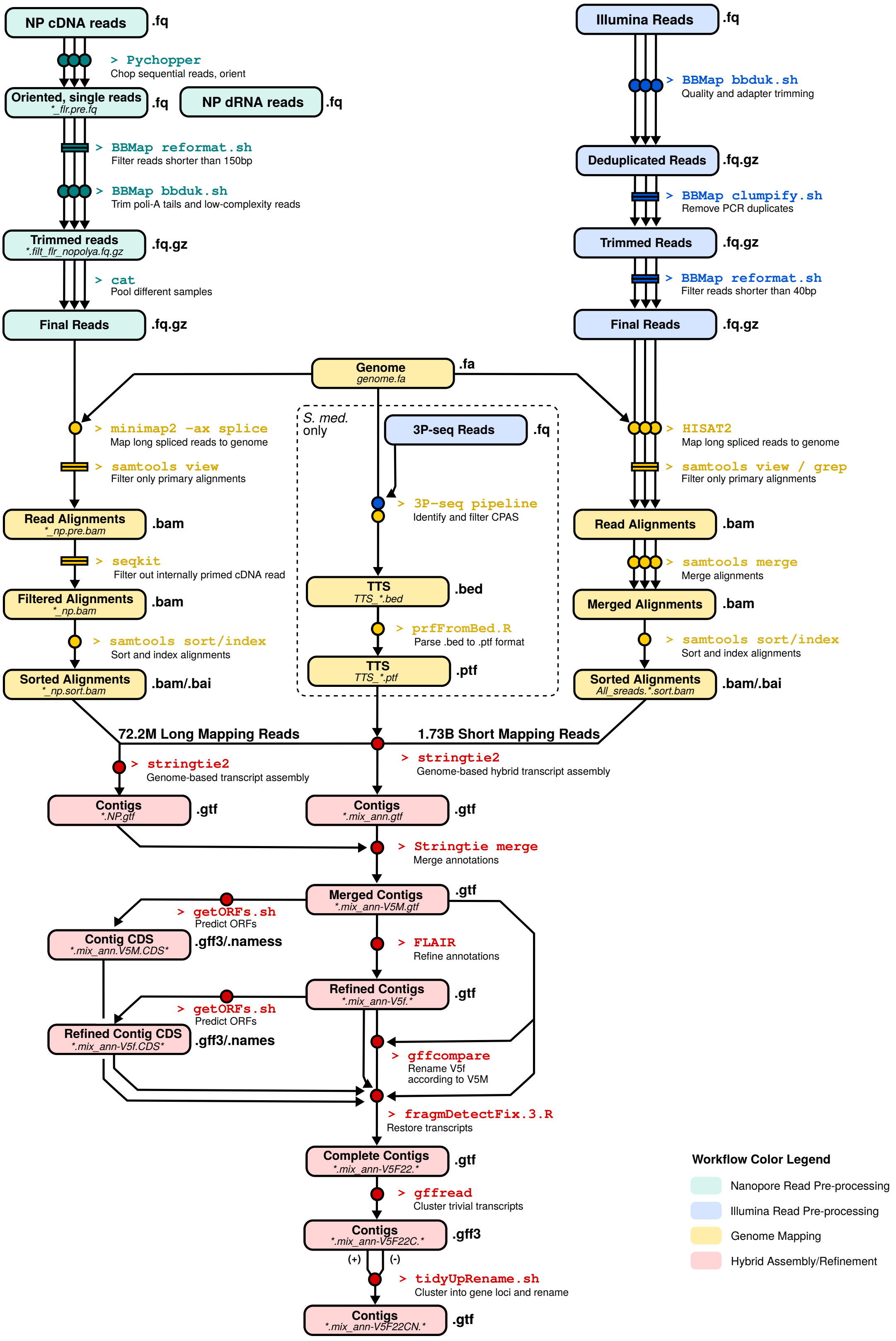


**Figure 6:** Detailed workflow of Part I

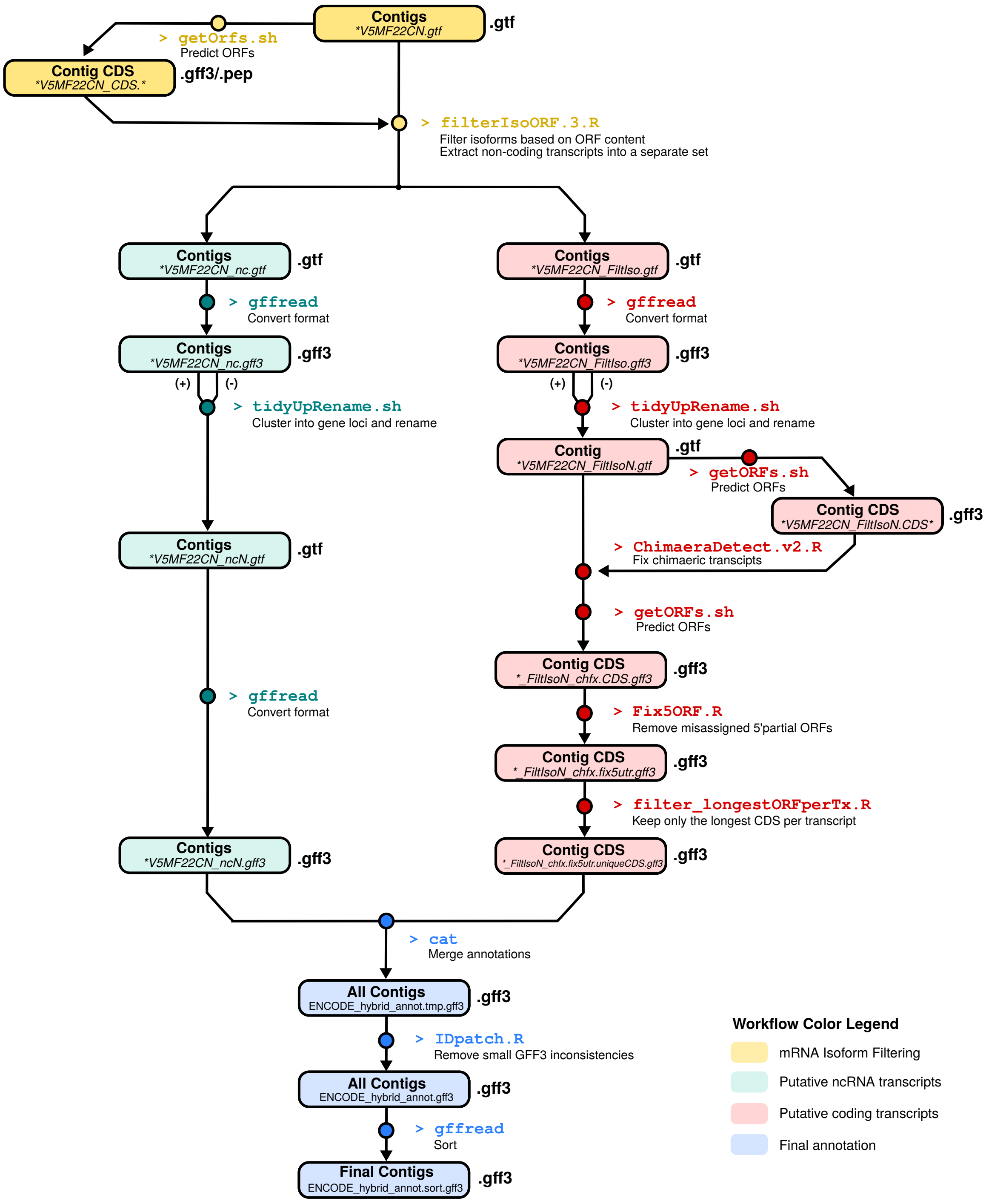


**Figure 7:** Detailed workflow of Part II
